# Oxygen-sensing regulatory architecture structures mammalian diversification

**DOI:** 10.64898/2026.07.21.739880

**Authors:** JB Smaers, A Gil-Gomez, REM Rickaby, CW Pugh, C West, P Aggarwal, A Hecker, A Chen, C Wen, M Riessland, JS Rest

## Abstract

The HIF oxygen-sensing pathway traces to the last metazoan common ancestor ∼800 million years ago and is conventionally viewed as a conserved cellular stress-response module. Whether this ancestral system has contributed to mammalian diversification at macroevolutionary timescales remains unexplored. We analyzed sequence-encoded TF-gene regulatory architecture for 34 transcription factors and 705 genes in 10 oxygen-sensing pathways across 239 mammalian species.

Oxygen-sensing regulatory architecture carries strong clade-structured evolutionary signal. The primary axis of variation tracks a fast-slow life history gradient, marked by rewiring of growth-control and tumor suppressor hub genes. A second axis recovers the monotreme-marsupial-placental transition and aligns with the decline in atmospheric O_2_ from the Permo-Carboniferous maximum toward present-day levels^1^. Orthogonal axes encode distinct ecological regulatory strategies; two later axes separately resolve HIF-compatible binding-site architecture and dominant TF-family assignment, identifying regulatory strategies associated with powered flight and hibernation. This multidimensional space also informs Peto’s paradox, suggesting that relative cancer resistance tracks the combination of tumor-suppressor enrichment and coordinated HIF-complex assignment.

Together, these results indicate that regulatory configurations arise at major evolutionary transitions and persist coherently across descendant lineages through a punctuated mode of regulatory evolution, providing genomic-level evidence for Simpson’s adaptive zones and a mechanism for evolutionary stasis. These findings reframe oxygen sensing as a regulatory hub in mammalian diversification, with stable patterns of TF-family assignment configurations emerging as a structuring force in macroevolution.

**Brief:** Ancient molecular processes such as oxygen-sensing, whose HIF-pathway dates to the origin of animals ∼800 million years ago, are typically regarded as ‘conserved’ across lineages. How such deeply ancestral systems have contributed to mammalian diversification remains largely unexplored. By analyzing the oxygen-sensing regulatory architecture (which transcription factors regulate which genes) across 239 mammalian species, we find that oxygen-sensing regulatory rewiring tracks placental evolution, atmospheric O_2_, life history evolution, ecological specializations, and cancer resistance. Major radiations occupy discrete, heritable configurations established at key phylogenetic transitions and subsequently retained across descendant lineages through near-neutral within-regime drift, revealing regulatory architecture lock-in as a structuring force in macroevolution.

## Background

Oxygen sensing is among the most ancient molecular systems in animals. The hypoxia-inducible factor (HIF) pathway can be traced to the last metazoan common ancestor approximately 800 million years ago, predating the Cambrian explosion by over 250 million years, and its core components (HIF1A, PHD, FIH, and VHL) are conserved across eumetazoan phyla^2^. Despite this antiquity and ubiquity, oxygen sensing is conventionally regarded as a relatively circumscribed cellular stress-response module: a system that detects hypoxia, stabilizes HIF-α subunits, and activates a cassette of target genes involved in erythropoiesis, angiogenesis, and glycolytic metabolism. Whether and how oxygen-sensing regulation has contributed to animal diversification at macroevolutionary timescales remains largely unexplored.

A separate but convergent problem concerns the evolutionary informativeness of transcription factor binding sites (TFBS). King and Wilson^3^ proposed that morphological evolution proceeds primarily through changes in gene regulation rather than protein sequence, and fifty years of evidence from evolutionary developmental biology broadly supports this thesis. Yet at the molecular level, individual TFBS undergo rapid evolutionary turnover, with estimated site-level half-lives ranging from approximately 5 to 150 million years^4^ (the time for half of all binding sites present in an ancestral genome to be lost or replaced). This turnover means that functionally conserved regulatory elements can show low sequence similarity even among closely related species, and that the majority of experimentally identified binding sites in any one genome are not conserved across mammals. Transcriptional networks themselves appear to evolve at a common rate across metazoan lineages regardless of genome-wide divergence rates^5^, further complicating efforts to extract adaptive signal from binding site data. This has raised a persistent question: does TFBS architecture retain adaptive evolutionary signal at macroevolutionary timescales, or does the noise of binding site gain and loss overwhelm phylogenetic and functional information?

We addressed both questions simultaneously by analyzing whole-genome TFBS architecture of 34 TFs and 705 genes spanning 10 KEGG pathways directly involved in oxygen sensing (Table ED1), across 239 mammalian species. Our approach differs from conventional comparative genomics in a critical respect. Previous studies of cis-regulatory evolution have tended toward two broad strategies: comparing orthologous regulatory elements or alignable sites to infer constraint and turnover^6,7^, or comparing TF occupancy across a small number of species and tissues using ChIP-seq^6^. Both become difficult to extend across deep divergence and hundreds of genomes because individual TFBS have half-lives of approximately 5-150 million years and enhancers turn over at comparable rates^4,7–9^. We instead analyze the regulatory architecture as a species-by-gene-by-TF map of sequence encoded binding site potential. Each species thereby produces a unique regulatory map encoding its sequence-level regulatory potential, allowing us to compare predicted TF-gene associations across species (see Methods). Crucially, this does not require identifying or aligning specific orthologous binding sites across species and is therefore robust to the turnover that confounds site-level and occupancy-based approaches.

## Results

To summarize regulatory variation across the oxygen-sensing gene set in 239 species, we applied principal component analysis (PCA) to the matrix of GC-corrected TFBS abundance values for 12,919 gene-TF pairs. This yielded a multi-dimensional regulatory architecture space in which each species occupies a position determined by its pattern of predicted TF-gene regulatory architecture across oxygen-sensing genes (Fig. 1). Each axis of this space represents an orthogonal dimension of cross-species variation in regulatory architecture, and species proximity within it reflects similarity in the overall pattern of TF-gene regulatory architecture, rather than conservation of any individual binding site.

**Figure 1.**
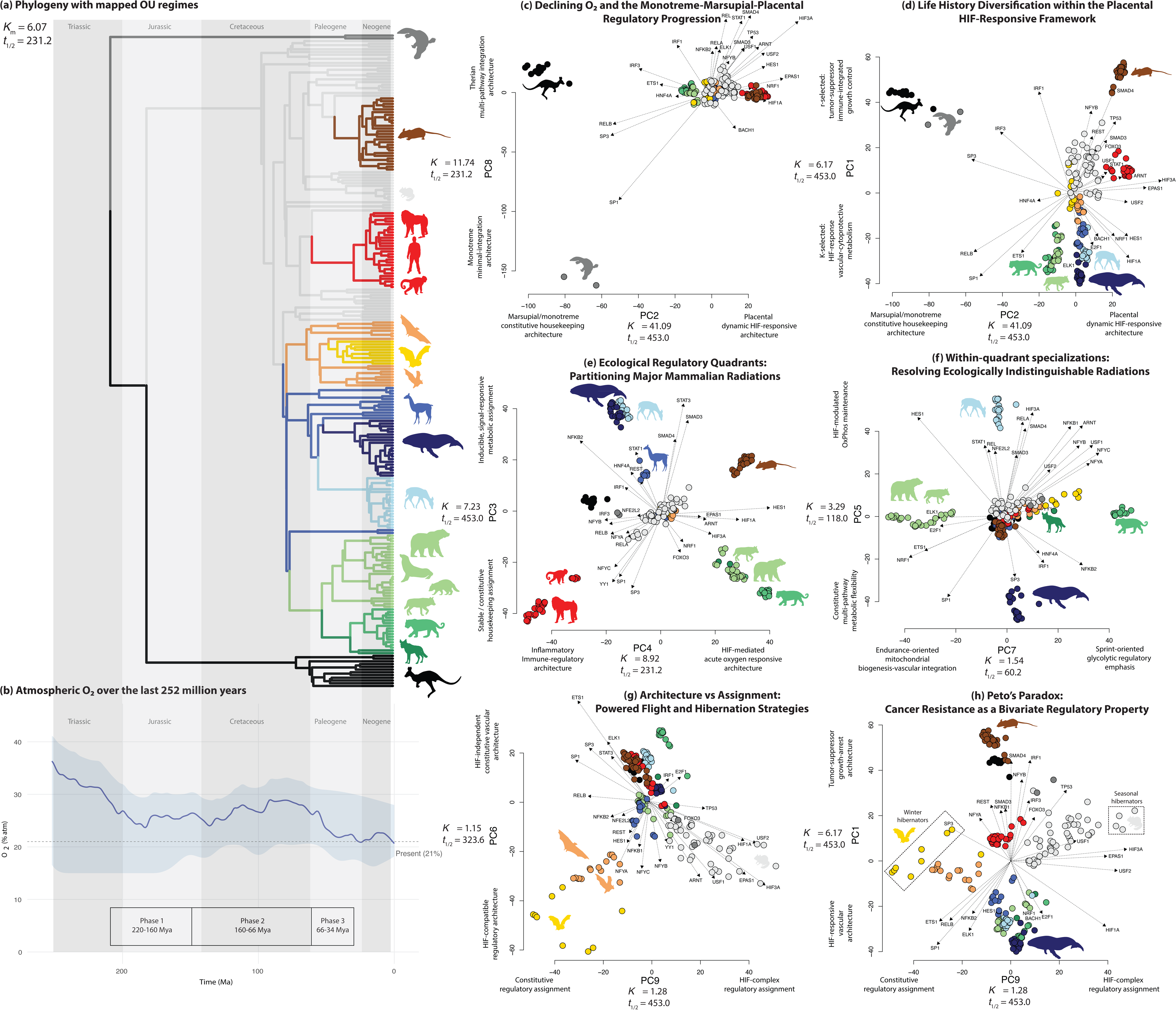
Principal component analysis of oxygen-sensing regulatory architecture across 239 mammals. (a) Phylogeny of the 239 mammalian species included in the study. Colors represent different OU-inferred regulatory regimes, each with a distinct multivariate mean across 9 PC axes; clade colors match species colors in panels (c)-(h). Phylogenetic half-life (*t*_1/2_) and phylogenetic signal (*K*_m_) from the joint 9-axis scOU model indicate near-neutral within-regime dynamics: regulatory configurations are determined by shift inheritance at major transitions rather than by continuous attraction toward a fixed optimum. (b) Atmospheric O_2_ levels over the last 252 million years (consensus curve from Mills et al.^1^). Shaded envelope reflects inter-model uncertainty. (c) PC8 vs PC2 recovers the monotreme-marsupial-placental transition (PC2: SP1-to-HIF assignment transition at core metabolic genes; PC8: degree of multi-pathway integration), showing a three-stage transition from ancestral constitutive architectures calibrated to post-Carboniferous hyperoxia to dynamically HIF-responsive placental architectures as O_2_ declined toward present-day levels. *t*_1/2_ and *K* indicate axis-specific phylogenetic half-life phylogenetic signal. (d) PC1 vs PC2 maps the diversification of placentals within this framework, with PC1 capturing the r-K life-history and body-size gradient and PC2 capturing the shift from ancestral constitutive control to dynamic HIF-responsive regulation, yielding four non-overlapping quadrants corresponding to major placental radiations. (e) PC3 vs PC4 partitions mammals into four ecological regulatory philosophies (constitutive vs inducible control and HIF-mediated vs inflammatory oxygen responsiveness) separating primates, carnivorans, cetaceans-plus-ruminants, and myomorph rodents into distinct adaptive zones. (f) PC5 vs PC7 resolves within-quadrant specializations that are invisible on PC3-PC4, distinguishing rumen fermenters from cetaceans by their contrasting solutions to chronic hypoxia and separating feliform from caniform carnivorans by sprint-focused versus endurance-focused metabolic assignment. (g) PC6 vs PC9 separates HIF-independent from HIF-compatible architectures and, within the latter, distinguishes lineages with HIF-complex assignment from those with constitutive assignment, revealing divergent regulatory strategies associated with powered flight and deep hibernation in bats versus ground squirrels. (h) PC1 vs PC9 defines a two-dimensional regulatory space in which tumor-suppressor investment (PC1) and HIF-complex assignment (PC9) jointly predict relative cancer protection, providing a regulatory perspective on Peto’s paradox by showing that species differ not by body size alone but by their position in this bivariate regulatory landscape.

Notably, phylogenetic signal^10^ across all individual gene-TF pairs is weak (mean Blomberg’s *K*_m_ = 0.177), consistent with the rapid turnover of individual binding sites and supporting the use of standard rather than phylogenetic PCA (see Methods). By concentrating the dispersed phylogenetic covariance shared across thousands of individually labile traits into composite axes, PCA allows regulatory architecture to be analyzed as a network-level property rather than a collection of independent sites. The resulting first nine PC axes show strong phylogenetic signal (*K* > 1 on all nine axes, *P* = 0.001; Fig. 1), meaning closely related species are substantially more similar in their composite TF-gene regulatory architecture than expected under Brownian motion. Axes beyond PC9 did not carry significant phylogenetic signal and were not retained for subsequent analyses.

To characterize the mode of evolution generating this signal, we fitted multivariate Ornstein-Uhlenbeck shift models (PhyloEM^11^; scalar OU process across all nine axes jointly) to the PC scores. Estimated phylogenetic half-lives, the time required for a lineage displaced from its regulatory configuration to return halfway toward it, met or exceeded the tree height on every axis (joint 9-axis: 231 Myr; per-axis range 60-453 Myr; tree height 217 Myr; Fig. 1), with the selection-strength parameter α converging to the lower bound of the search grid in all fits. These values are therefore lower bounds indicating uniformly weak within-regime attraction (α indistinguishable from zero) rather than discriminating axis-specific tempos. Taken together, the long phylogenetic half-lives and supra-BM signal (*K*>1) indicate that this pattern does not reflect ongoing stabilizing selection within clades, which would reduce K toward or below 1 by constraining within-clade variance, but rather the inheritance of discrete regime shifts at clade boundaries. Each shift establishes a new regulatory baseline that all descendant lineages inherit and retain, producing the between-clade clustering that elevates K above 1 without requiring continuous stabilizing pull. Under a multi-optimum (shift) OU process, K > 1 is the expected signature of divergent, coherently inherited clade optima, not a departure from the OU framework: between-regime separation inflates K above the Brownian expectation regardless of the within-regime restoring force.

This contrast between weak signal at the level of individual gene-TF pairs (*K*_m_ = 0.177) and strong signal at the level of composite PC axes (*K* > 1) is the key property that both motivates and validates our approach. Individual binding sites are labile and individually near-neutral, but their collective configuration is architecturally conserved and inherited as a coherent unit through punctuated shifts; the signature of punctuated regulatory evolution. Regulatory architecture is thus a network-level emergent property whose evolutionary coherence cannot be detected, or explained, at the level of individual binding-site features.

We then used a complementary TF-switching analysis to interpret the retained PC axes biologically. For each axis, this analysis identifies genes represented in both the positive and negative loading tails but with different TF partners, thereby inferring which TFs lose or gain regulatory prominence across that dimension of regulatory-architecture variation. Thus, PCA defines the major axes of regulatory architecture, while switching analysis identifies the hub genes and transitions in dominant TF-family assignment that provide their functional interpretation. We use dominant TF-family assignment as shorthand for the individual TFs or broader functional TF classes preferentially associated with one end of a PC axis, as inferred from TF-specific loading patterns and supported by the directionality of switching events. At the gene level, TF assignment refers to the TF partner or partners represented with that gene in a given extreme loading tail; a switch occurs when the same gene is represented in opposite tails with different TF partners. This differs from regulatory architecture, which describes the multivariate pattern TF abundance across all retained oxygen-sensing gene-TF pairs. Both describe sequence-encoded regulatory potential rather than realized transcription.

Before interpreting these axes biologically, we tested whether the regulatory architecture space depended on scanning window, coordinate anchor, or genome-annotation pipeline. The primary 500-bp ATG-anchored solution was highly congruent with alternative promoter-window definitions, including wider ATG-anchored windows and a TSS-anchored window (Procrustes r ≥ 0.984, Mantel r ≥ 0.973 across three alternative window and two anchor conditions). Recomputing the analysis on species and gene-TF pairs shared between the NCBI/OrthoFinder and TOGA annotation^12^ sets also reproduced the regulatory architecture space (Procrustes r ≥ 0.977, Mantel r ≥ 0.969), with per-axis congruence strongest for the principal within-placental axes and moderate for the PC3/PC4 ecological axes (Supplementary information). Because the TOGA comparison lacks monotremes and marsupials, it cannot evaluate the deep PC2/PC8 monotreme-marsupial-placental transition.

Using this framework, we recover a staged transition from constitutive regulation toward HIF-centered assignment aligned with the monotreme-marsupial-placental transition and long-term changes in atmospheric O_2_, orthogonal ecological structuring of regulatory architectures across mammalian radiations, and covariation between regulatory architecture and cancer resistance across species. Within this context, we use ‘adaptive signal’ to denote the phylogenetic pattern in which regulatory configurations diverge between clades at key transitions and are subsequently conserved within clades across ecological diversification, a pattern quantified below using Blomberg’s K and multivariate OU regime-shift models.

### 1. Declining O_2_ and the evolution of dynamic HIF-responsive architecture in placentals

The monotreme-marsupial-placental transition recovered by PC2 (capturing SP1-to-HIF assignment transition at core metabolic genes) and PC8 (capturing the degree of multi-pathway integration) maps onto the long-term trajectory of atmospheric O_2_ from Permo-Carboniferous hyperoxia (∼28-31% at 220 Ma) toward present-day levels (∼21%), with O_2_ having already declined to ∼23-27% at the marsupial-placental split (∼160 Ma), temporarily rising to ∼28-29% during the mid-to-late Cretaceous before declining to ∼22% by the late Eocene^1^ (Fig. 1a-d). All atmospheric O_2_ estimates cited here are derived from the COPSE biogeochemical model (Mills et al.^1^) and carry substantial uncertainty, particularly for pre-Cretaceous intervals, where inter-model ranges can span ±5-8 percentage points; the qualitative trajectory, elevated Permo-Carboniferous levels declining toward modern values, is more robustly supported than any individual estimate. This non-monotonic trajectory provides the mechanistic context for a proposed three-phase regulatory transition characterizing mammalian evolution that is jointly captured by PCs 1, 2 and 8.

#### Phase 1: Constitutive lock-in under declining post-Carboniferous O_2_ (∼220-160 Ma)

The monotreme-therian divergence (217 Ma in the phylogeny used here) and the marsupial-eutherian divergence (∼160 Ma) both occurred as atmospheric O_2_ was declining from its Permo-Carboniferous maximum (estimated at ∼31-37% at ∼263 Ma), with modeled values around ∼28-31% at 220 Ma and ∼23-27% at 160 Ma (Fig. 1b; uncertainty ranges reflect inter-model variation)^1^. Although tissue PO_2_ is always substantially lower than atmospheric PO_2_, with the gradient from atmosphere to mitochondria spanning ∼10-20 kPa in contemporary mammals across different tissues^13^, higher atmospheric O_2_ calibrates the upper bound of the oxygen delivery system^14^, shifting the full physiological O_2_ range upward. The catalytic and structural properties of PHD enzymes that confer O_2_-sensing function are conserved from the simplest animal, *Trichoplax adhaerens*, to *C. elegans* to humans^15,16^, indicating that the kinetic basis of graded HIF stabilization was present in the ancestors of early mammalian lineages. At Permo-Carboniferous atmospheric levels (estimated ∼31-37%), this upper-bound shift would have maintained tissue PO_2_ at values closer to the upper portion of the PHD-sensitive range, where graded HIF stabilization is reduced, providing limited selective advantage for dynamic HIF-responsive programs. In effect, this left the oxygen-sensing system with little usable dynamic range: because PHDs were already working at or near their maximum capacity, small changes in tissue oxygen likely could no longer translate into graded changes in HIF stabilization. Constitutive SP1/SP3/RELB-mediated transcription of oxygen-sensing genes (which is the architecture retained by monotremes and marsupials on negative PC2) could be argued to be the energetically efficient strategy under these conditions because there is little selective advantage to dynamic HIF-responsive programs when ambient O_2_ keeps PHDs permanently saturated and HIF-α permanently destabilized. The difference in O_2_ conditions at the two divergences, ∼28-31% at the monotreme-therian split versus ∼23-27% at the marsupial-eutherian split, may explain the greater extremity of the monotreme regulatory architecture (PC8 = −158.5) relative to marsupials (PC8 = +18.6): monotremes diverged when PHDs were operating closer to full saturation (∼31% O_2_), while marsupials acquired SMAD4/STAT1/NF-κB pathway integration at ∼23-27% O_2_, a level near the boundary of the PHD-sensitive range. The coherence of marsupial clustering on PC2 (SD = 6.19 across 12 ecologically diverse species) confirms early phylogenetic fixation (Table ED2). This lock-in has broad phenotypic consequences: constitutive SP1-mediated control of PRKAA1 (AMPK; 48 PC2 switches), SLC2A4 (GLUT4; 35 switches), RHEBL1 (mTOR; 24 switches), ENO4 (glycolysis; 15 switches), and UQCRC1 (Complex III; 12 switches) produces a metabolic architecture (Fig. 2a, ED1a, Table ED3) that cannot dynamically couple energy sensing to oxygen availability or support hemochorial placentation, consistent with the marsupial BMR deficit^17^ and absence of thermogenic brown fat^18^.

**Figure 2.**
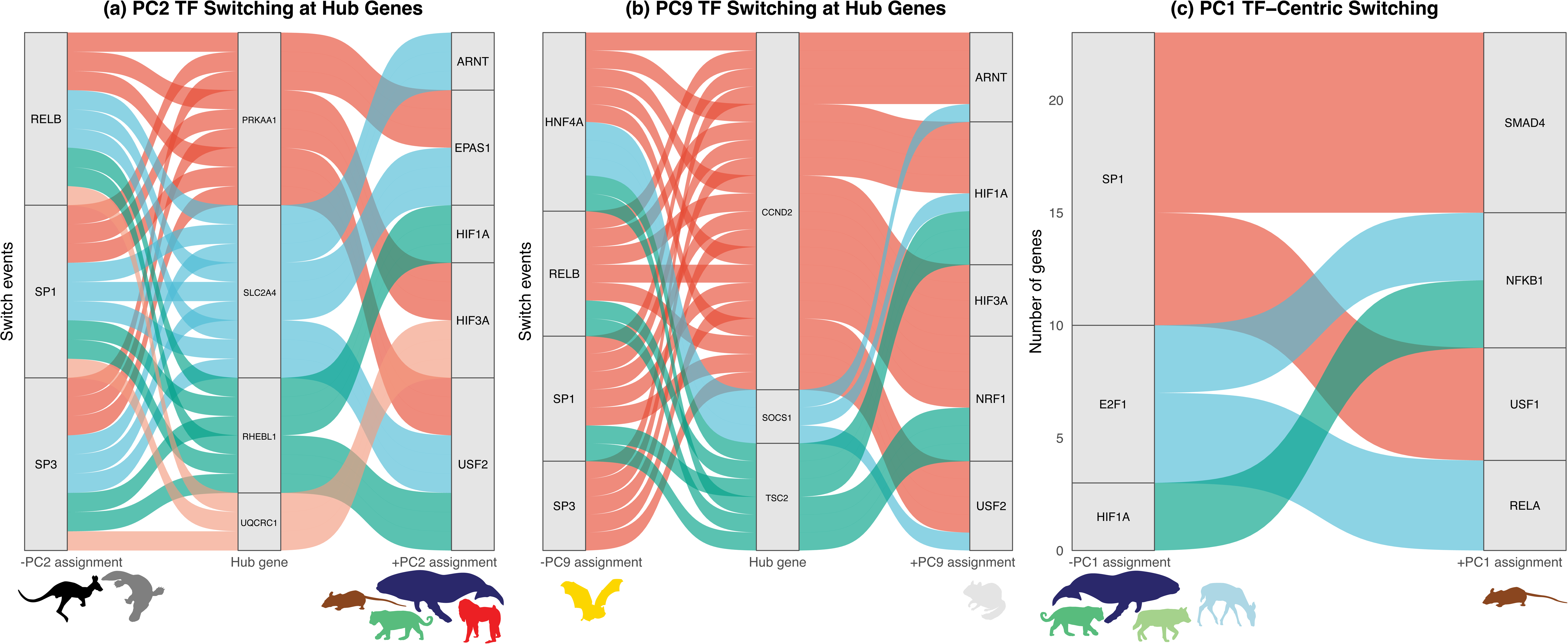
TF switching identifies systematic regulatory rewiring at hub genes across major evolutionary transitions. Alluvial plots show directed TF switching events along three independent PC axes, where ribbons connect TFs losing assignment (left) to those gaining assignment (right) through individual hub genes (center, gene-centric panels A-B) or as aggregated TF pairs (TF-centric panel C). **(A)** Gene-centric switching on PC2 (marsupial → placental axis). Four metabolic hub genes [PRKAA1 (AMPK, energy sensing), SLC2A4 (GLUT4, glucose transport), RHEBL1 (mTOR signaling), and UQCRC1 (Complex III, OXPHOS)] switch from constitutive SP1/SP3/RELB assignment (retained in marsupials) to HIF-responsive regulation by HIF1A, HIF3A, EPAS1, ARNT, and USF2 (acquired in placentals). This transition corresponds to the Phase 2 transition described in the text. Only the core constitutive and HIF-family TFs are shown (full switching landscape in Extended Data Fig. ED1A). **(B)** Gene-centric switching on PC9 (bat ↔ ground squirrel hibernation axis). CCND2 (cell-cycle arrest, 28 switches, 58.8× enrichment), TSC2 (mTOR inhibition, 8 switches), and SOCS1 (immune suppression, 12 switches) switch from constitutive/metabolic assignment (SP1, SP3, HNF4A, RELB) toward bats to HIF-complex assignment (HIF1A, HIF3A, ARNT, NRF1, USF2) toward ground squirrels. These hub genes serve dual roles in the cancer framework (§4): the same regulatory rewiring that distinguishes hibernation strategies also calibrates cell-cycle and growth-control checkpoints to species-specific cancer protection. Only the principal constitutive and HIF-associated TFs are shown (full switching landscape in Extended Data Fig. ED1B). **(C)** TF-centric switching on PC1 (r-K life-history axis). Ribbon width represents the number of genes sharing each directed TF_neg → TF_pos pair. The dominant switch, SP1 → SMAD4 (8 genes, 8.1× enrichment, *P BH* = 0.0003), captures transition from HIF-responsive/vascular architecture (negative PC1, K-selected ferungulates) to tumor suppressor-enriched architecture (positive PC1, r-selected myomorphs). Together, Panels B and C visualize the two independent regulatory layers of the bivariate PC1 × PC9 cancer framework: tumor suppressor investment (PC1) and HIF-complex assignment (PC9), whose intersection structures species-level cancer vulnerability.

We acknowledge that two caveats apply to this argument. First, ancestral haemoglobin precursors likely differed from contemporary Hb in oxygen affinity and cooperativity, making precise reconstruction of ancestral tissue PO_2_ unavailable. Second, the apparent K_m_ for O_2_ of ancestral PHD-like hydroxylases is unknown; it is therefore possible that the HIF system was functionally active at the PHD-kinetic level during periods of high atmospheric O_2_, and that constitutive regulatory architecture in early mammalian lineages reflects other evolutionary factors. The argument from atmospheric O_2_ to PHD saturation should therefore be understood as a directional hypothesis, higher atmospheric O_2_ reduces the selective advantage for dynamic HIF-responsive programs, rather than a precise mechanistic reconstruction of ancestral PHD activity.

#### Phase 2: HIF-responsive architecture innovation on the placental stem lineage (∼160-66 Ma)

Along the placental stem lineage, as atmospheric O_2_ fluctuated around ∼25-29%, already substantially below the ∼31-37% Permo-Carboniferous maximum that had calibrated the constitutive architecture of monotremes and marsupials, PHD enzymes began operating further below their saturation ceiling, creating conditions under which dynamic HIF stabilization in response to physiological oxygen gradients became selectively meaningful^1^ (Fig. 1b). This is evidenced by the SP1-to-HIF regulatory transition captured by PC2: 116 SP1, 82 SP3, and 62 RELB switches, overwhelmingly unidirectional (96-99% bias) from constitutive to HIF-responsive architecture (Fig. 4A, Table ED4). This transition rewired the same conserved hub genes (PRKAA1, SLC2A4, RHEBL1, ENO4) from SP1/SP3 to HIF1A/EPAS1/ARNT assignment, creating a unified oxygen-energy sensing network (Fig. 2A). Three placental-exclusive innovations that enabled the evolution of the placenta emerged: HIF3A/IPAS negative feedback (47 exclusive positive switches), enabling transient HIF activation without pathological overactivation; EPAS1 chronic sensing (36 exclusive positive switches), supporting sustained adaptive responses during prolonged gestation; and USF2 HIF-cooperative modules (39 exclusive positive switches), enabling tissue-specific metabolic responses (Fig. 4A, ED1c, Table ED4). Together, these innovations enabled hemochorial placentation by coupling energy sensing (PRKAA1), glucose metabolism (SLC2A4, ENO4), and growth signaling (RHEBL1) to oxygen availability across the oxygen gradients of early pregnancy; gradients whose physiological informativeness was enhanced relative to the hyperoxic conditions that had constrained the ancestral architecture. Superordinal placental divergence (∼102-66 Ma) coincided with a relatively elevated but stable O_2_ (∼27-29% during the mid-to-late Cretaceous^1^), driven by continental fragmentation providing geographic isolation. The HIF-responsive architecture evolved in Phase 2 was exapted to function across this range, enabling reproductive diversification^19^ without requiring further regulatory innovation.

#### Phase 3: Paleogene O_2_ decline and adaptive radiation (66-34 Ma)

The post-K-Pg interval saw atmospheric O_2_ resume its long-term decline from ∼25-29% at the Cretaceous-Paleogene boundary toward ∼22% by the late Eocene^1^ (approaching modern levels), a ∼5.5 percentage-point drop over 32 million years that deepened the PHD-sensitive regime established in Phase 2 and coincided with the most dramatic radiation of placental mammals in the fossil record. The PC1 gradient (80.9 units from myomorphs to ferungulates, Table ED2) represents diversification of regulatory architecture within the HIF-responsive framework established in Phase 2 (Fig. 1d). This gradient, correlating with body size and the r-K continuum, reflects systematic rewiring of growth-control hub genes (WNT1, TSC2, BAD, CDKN1B, PLK1) between oxygen-linked cytoprotective control in K-selected lineages and inflammatory-immunometabolic control in r-selected lineages (Fig. ED1b, Table ED3). The continued descent of O_2_ into the range where tissue-level hypoxia becomes increasingly likely during metabolic activity and embryonic development favored this diversification: it enabled the ordinal radiation of artiodactyls, perissodactyls, and early cetaceans into the constitutively high-capacity architecture of negative PC1, lineages whose large body size and high absolute metabolic demands require robust oxygen delivery, and supported the cerebral metabolic demands of primates, whose enhanced HIF-responsive architecture (PC2 = +22.8) enabled neurovascular coupling in expanding brains. Most living placental orders appear in the early Eocene, and the four-quadrant PC1 × PC2 space maps directly onto this radiation: HIF-responsive with immune integration (myomorphs, primates), HIF-responsive with constitutive high capacity (cetaceans, ruminants), reduced HIF with constitutive high capacity (carnivorans), and ancestral constitutive (marsupials) (Fig. 1d).

### 2. Ecological structuring of oxygen-sensing regulatory architecture across mammalian radiations

The life-history and placental framework represented in PCs 1, 2, and 8, however, explains only part of the diversity in oxygen-sensing regulation. Other axes (PC3-PC7) reorganize the same genes and TFs to address a different question: how mammalian lineages with this shared genomic toolkit deploy distinct regulatory architectures to meet their ecological oxygen challenges (Fig. 1e,f,g).

This wiring operates at two hierarchical levels, orthogonal to the PC1/PC2 life history and placentation axes. PC3 and PC4 establish broad regulatory philosophies: PC3 separates constitutive housekeeping control (SP1/SP3/NFY; negative) from inducible, signal-responsive control (STAT3/SMAD3/HNF4A; positive), while PC4 separates NF-κB/IRF-dominated architecture (NFKB2/IRF3/RELB; negative) from HIF-mediated acute oxygen responsiveness (HIF1A/ARNT/HES1; positive). Together they partition mammals into four non-overlapping quadrants, each occupied by a major radiation (Fig. 1e, Table ED2). PC5 and PC7 resolve within-quadrant specializations: PC5 separates cetaceans from Pecora, indistinguishable on PC3, via different regulatory mechanisms at HIF-pathway genes, while PC7 separates feliformes from caniformes, co-occupying PC3−/PC4+, via NRF1-mitochondrial versus NFYC-governed regulatory emphasis (Fig. 1f). That the same two lineages show near-identity on one axis and the largest divergence in the entire dataset (91.6 units) on another, and that this analytical logic repeats independently for carnivorans (shared PC3−/PC4+, 75.0-unit PC7 divergence), argues that the PCA decomposes biologically coherent regulatory modules rather than partitioning variance arbitrarily.

The PC3/PC4 space highlights network-level crosstalk between HIF and NF-κB signaling. While this relationship is complex and context-dependent, extensive evidence supports their co-regulation and mutual crosstalk at the transcriptional level; the full mechanistic details of their interaction remain an active area of research^20^. In this space primates occupy the constitutive/NF-κB-enriched quadrant (PC3−/PC4−), showing a directional gradient from strepsirrhines through platyrrhines to catarrhines, placing oxygen-sensing genes under stable SP1/NFY assignment enriched for NF-κB/IRF elements rather than canonical HIF pathways (Fig. 1e). This architecture is consistent with a lineage whose brains consume up to 20-25% of resting oxygen^21^. TF switching confirms this: HIF1A shows the strongest directional bias on PC4 (26/29 switches gained toward PC4+, P = 0.0003, Table ED4), meaning genes systematically lose HIF1A architecture toward primates, while NFYC (15/18 retained, P = 0.041) and RELB (15/19 retained, P = 0.080) persist. Cetaceans and Pecora converge in the inducible/NF-κB-enriched quadrant (PC3+/PC4−), their PC3 means differing by only 0.49 units despite 55 million years of independent evolution (Table ED2). Both face predictable chronic hypoxia (diving apnea and ruminal fermentation) and share STAT3/SMAD3 inducible architecture with NF-κB/IRF-enriched rather than HIF-enriched control. Non-ruminant ungulates (pigs, hippos, camelids; PC3 = +14.9) occupy intermediate positions confirming that the cetaceans and Pecora extremes are derived. HNF4A shows the strongest PC3 bias (24/27 switches toward PC3+, P = 0.001), and the directed switch RELA → HNF4A (P = 0.031) captures transition from NF-κB to metabolic assignment (Fig. ED2a, Table ED5). Carnivorans occupy the constitutive/HIF-enriched quadrant (PC3−/PC4+), combining stable SP1/NFY baseline with HIF1A/ARNT architecture. Feliformes (PC4 = +36.2, Table ED2) are more extreme than caniformes (+26.3), consistent with sprint versus endurance predation. Myomorphs occupy the maximally flexible quadrant (PC3+/PC4+; SD < 2 on both axes), combining inducible control with HIF-enriched architecture, consistent with small mammals whose extreme metabolic rates and fast life histories demand maximal regulatory versatility.

PC5 resolves how cetaceans and Pecora, despite indistinguishable PC3 positions, employ fundamentally different regulatory strategies (Fig. 1f). Their 91.6-unit (Table ED2) divergence reveals that Pecora show increased HIF3A binding architecture (strength = +0.00166; Table ED6) with NF-κB elements (RELA = +0.00151), with OXPHOS complex genes (Complexes I, II, IV) prominent among the most extreme positive loadings, consistent with HIF3A dominant-negative modulation maintaining oxidative phosphorylation for volatile fatty acid oxidation under fermentation hypoxia. Cetaceans show SP1 architecture enrichment (strength = −0.00130) with individual metabolic genes from glycolysis (PGK2), TCA cycle (DLAT, PDHB, ACO2), and OXPHOS (NDUFB11, ATP5F1B) among the most extreme negative loadings, drawing from multiple metabolic modules rather than any single pathway. This pattern is itself consistent with metabolic flexibility: cetacean regulatory architecture does not lock onto a single metabolic pathway but instead positions individual genes from diverse pathways under SP1-mediated assignment, enabling rapid substrate switching during dive-recovery cycles. SP1 switching confirms this (25 neg, 6 pos; P = 0.022, Table ED4), and hub genes FZD4 and PGK2 (each 15 switches, P = 0.005, Table ED3) identify vascular and glycolytic regulation as the principal sites of flexibility (Fig. ED2b).

PC7 resolves how feliformes and caniformes differ in metabolic emphasis despite sharing the carnivoran quadrant (Fig. 1f). Caniformes show coordinated NRF1 mitochondrial biogenesis architecture (51 switches, 43 neg; P = 6 × 10⁻□, Table ED4) and ETS1 vascular architecture (41 switches, 36 neg; P = 6 × 10⁻□). TCA cycle and OXPHOS complex genes (Complexes I, III, IV) appear among caniform extreme-loading genes, consistent with NRF1-governed mitochondrial integration linking oxygen delivery to substrate oxidation. Feliformes show NFY complex enrichment (NFYC 24 switches, 23 pos; P = 2.2 × 10⁻□, Table ED4), alongside stress-response genes (ATF4, PIAS3), consistent with sprint predation physiology. NFYC assignment preferentially associates with metabolic and signaling targets at the feliform end. The directed switch NRF1 → NFYC (P = 0.021) captures this mechanistic core, with PDGFD (24 switches, P = 0.003) as the master hub gene, and PIAS3 (15 switches, P < 1×10⁻□) as a co-regulated target, shifting from HIF1A/NRF1-linked regulation in caniformes to NFYC/NFYA-linked regulation in feliformes (Fig. 3a,b, Table ED5,ED2).

**Figure 3.**
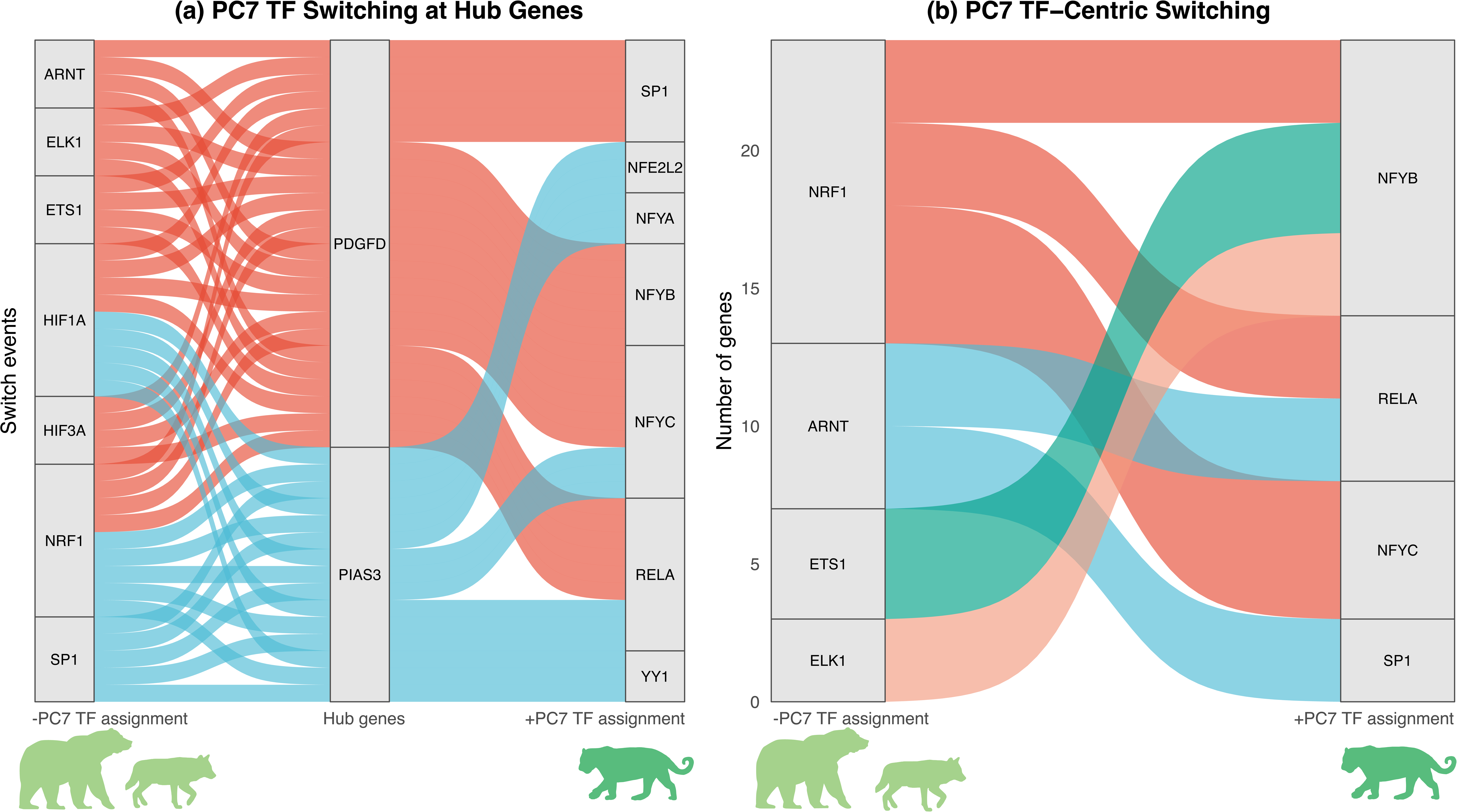
Gene-centric and TF-centric switching resolve the regulatory basis of the caniform-feliform ecological divergence. PC7 separates caniformes (endurance predation, negative PC7) from feliformes (sprint predation, positive PC7) within the carnivoran radiation that shares an indistinguishable PC3-PC4 quadrant position. **(A)** Gene-centric switching at PDGFD and PIAS3, the two most enriched hub genes on PC7. PDGFD (24 switches, 97.2× enrichment, P_BH = 0.006) switches from coordinated HIF1A-ETS1-NRF1 assignment (caniform end, consistent with NRF1-governed mitochondrial biogenesis linking oxygen delivery to substrate oxidation for endurance predation) to NFYC-NFYB assignment (feliform end, consistent with NFY-mediated glycolytic-emphasis control supporting sprint predation physiology). PIAS3 (15 switches, 36.1× enrichment, P_BH = 0.006), a STAT3 inhibitor and stress-response gene, switches from HIF1A-NRF1-SP1 assignment (caniform end) to NFYA-NFYC-NFE2L2 assignment (feliform end), reinforcing the NRF1-to-NFYC transition at a co-regulated target consistent with sprint predation physiology. Both genes converge on the directed switch NRF1 → NFYC captured by Panel B. **(B)** TF-centric switching on PC7, showing directed TF pairs with ≥3 shared genes. Ribbon width represents the number of genes per pair. The dominant directed switch NRF1 → NFYC (5 genes: FZD9, MLST8, PARP1, PDGFD, PIAS3; 7.3× enrichment, P_BH = 0.021) captures the mechanistic core of this divergence, representing a transition from NRF1-coordinated mitochondrial integration to NFYC-mediated metabolic assignment. Together, Panels A and B demonstrate the complementarity of gene-centric (single-gene resolution, all TF connections) and TF-centric (population-level TF pair frequencies across multiple genes) switching analyses. Pinnipeds retain caniform NRF1-mitochondrial architecture on PC7 despite 25 million years of aquatic evolution and shared diving ecology with cetaceans, demonstrating that regulatory architecture constrains the space of accessible ecological solutions.

### 3. Powered flight and different strategies for hibernation

Where PCs 3, 4, 5 and 7 resolve how ecological strategies are encoded in oxygen-sensing regulatory architecture, PCs 6 and 9 separate two independently varying layers of the same regulatory system: HIF-compatible binding-site architectures (PC6) and dominant TF-family assignment (PC9). The joint PC6-PC9 space reveals how regulatory architecture and dominant TF-family assignment combine in adaptations involving extreme oxygen demand or oxygen limitation, illustrated here by powered flight in bats and deep hibernation in ground squirrels (Fig. 1g).

PC6 differentiates HIF-independent from HIF-compatible regulatory architecture. The positive end represents constitutive, always-on regulatory architecture mediated by ETS1 and ELK1, ETS-family factors that can cooperate with HIF-2α at specific vascular targets (e.g., Flk-1/VEGFR2) but whose primary vascular regulatory function operates independently of hypoxia signaling, combined with SP-family housekeeping factors (SP1/SP3). The dominance of ETS1 in PC6+ is confirmed in the TF switching analysis where it emerged as the defining switch target: 28 gene-TF switches towards PC6+ and 0 towards PC6-(*P*=1.1×10^−7^) (Fig. ED3b, Table ED4). ETS1 is a master vascular endothelial regulator that drives angiogenic gene expression independently of hypoxia, providing constitutive high-capacity oxygen delivery infrastructure. This architecture is characteristic of groups requiring predictable, sustained high metabolic rates: sprint predators (felids), foregut fermenters (ruminants), and large-brained primates. The negative end is defined by HIF transcriptional machinery, including all three HIF-α isoforms (HIF1A, EPAS1, HIF3A) plus the obligate dimerization partner ARNT (HIF1B). The presence of USF1/2 (upstream stimulatory factors that cooperate with HIFs) and the NFY complex (NFYA/B/C, mitochondrial biogenesis regulators) indicates this architecture supports oxygen-sensing-linked mitochondrial regulation. Consistent with this interpretation, NFYA/B/C and USF1 to ETS1 and SP3 emerge as the most dominant TF switching strategies along the PC6 axis (Table ED5), with the metabolic enzyme ACLY (ATP citrate lyase, 10 switches, 21-fold enrichment, Table ED3) exemplifying this rewiring by switching from NFY/REST assignment to ETS1 assignment (Fig. ED3a,b).

PC9 is characterized by an orthogonal differentiation of species with HIF-compatible regulatory architecture (PC6-), separating hibernating ground squirrels from bats (Fig. 1g). The positive end is defined by coordinated HIF-family regulatory assignment (HIF3A, EPAS1, HIF1A, USF1/2), while the negative end is defined by constitutive regulatory assignment (SP1, ETS1, RELB, SP3).

Ground squirrels, facing severe tissue hypoxia during long, stable torpor bouts, show HIF regulatory assignment (PC9+). Bats, including the hibernating vesper bats Myotis, show constitutive/non-HIF regulatory assignment (PC9-). This divergence is confirmed through systematic rewiring of TF logic: the dominant PC9 switches from negative to positive are SP1 to ARNT (6 switches, *P*=0.0002, Table ED5), HNF4A to HIF1A (5 switches, *P*=0.002), reflecting transitions from constitutive to inducible HIF control at shared targets (Fig. 2b, 4b). A striking directional bias in each of the three HIF-α isoforms (Fig. 4b, Table ED4) is consistent with this interpretation.

**Figure 4.**
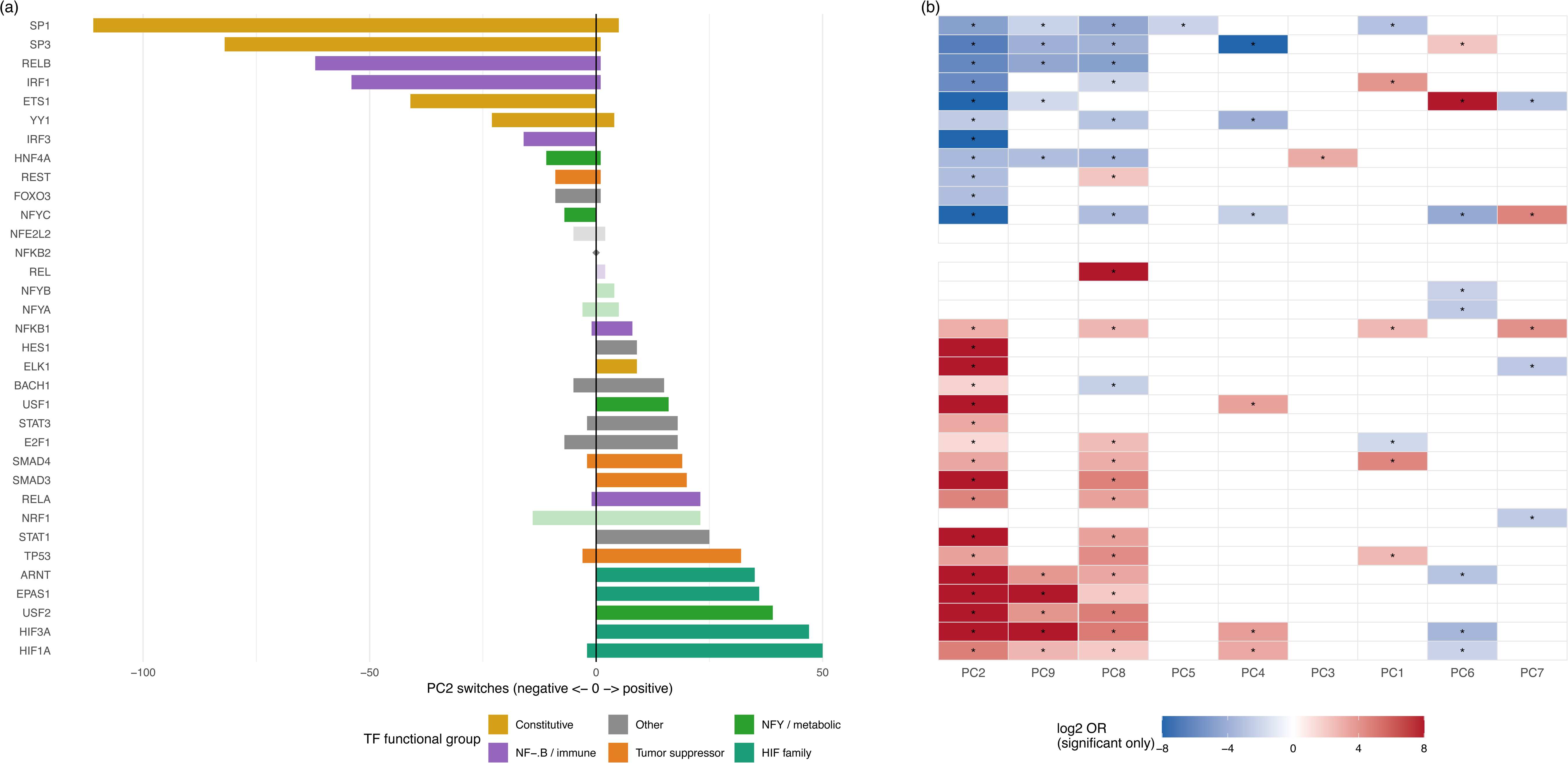
TF directional bias reveals the aggregate logic of switching across evolutionary axes (PC1-PC9). Where alluvial plots (Figures 2 and 3) show specific switching events at individual genes or TF pairs, TF involvement analysis quantifies the overall directional bias of each TF across all switching events on an axis. For each TF, a Fisher’s exact test compares the number of switching events placing that TF on the negative versus positive side, testing whether participation is significantly skewed toward one evolutionary pole. Both panels display all 34 TFs in a shared y-axis ordering derived from PC2 bias direction and magnitude: negative-biased TFs (losing assignment in placentals) appear in the upper rows, ordered by the number of negative-side switches (largest at top); TFs with no recorded switching events on PC2 occupy a middle zone; positive-biased TFs (gaining assignment) appear in the lower rows, ordered by the number of positive-side switches (largest at bottom). This three-zone shared ordering ensures perfect row-level alignment between panels, allowing horizontal visual tracking from a TF’s PC2 switching profile (Panel A) to its behavior across all nine axes (Panel B). **(A)** PC2 two-sided waterfall: all 34 TFs. TFs with switching events show two bars, one extending left for the number of negative-side switches and one extending right for the number of positive-side switches, so that both the dominant direction and the magnitude of any counter-directional switches are visible. TFs with no PC2 switches are marked by a grey diamond (−) at zero in the middle zone between the two bias groups. Bars are colored by TF functional class using a palette (teal, goldenrod, purple, orange, green, grey) chosen to be fully distinguishable from the heatmap’s blue-white-red diverging scale; opaque bars indicate significant directional bias (BH-corrected *P* < 0.05) and translucent bars indicate non-significant bias. Twenty-eight of 33 TFs with switches are significantly biased, with a sharp boundary separating constitutive TFs (goldenrod; SP1: 111 neg / 5 pos; SP3: 82 / 1; ETS1: 41 / 0) and NF-κB/immune TFs (purple; RELB: 62 / 1; IRF1: 54 / 1) in the upper rows from HIF-family TFs (teal; HIF3A: 0 / 47; EPAS1: 0 / 36; ARNT: 0 / 35; HIF1A: 2 / 50) and tumor suppressors (orange; TP53: 3 / 32) in the lower rows. The near-absence of opposite-direction bars for significantly biased TFs confirms the near-complete unidirectionality of the constitutive-to-HIF transition described in §1 Phase 2. The grey diamond placeholders identify TFs that do not participate in the PC2 transition but whose cross-axis activity is visible in Panel B, demonstrating axis-specific TF deployment. **(B)** TF × PC heatmap (PC1-PC9): Directional bias for all 34 TFs across the nine main axes. Cells are colored by log2OR only when bias is statistically significant (BH-corrected P < 0.05, marked with asterisks); non-significant cells are rendered white to prevent low-count TFs with extreme but unreliable log2OR values (e.g., 2/0 switches yielding log2OR = ∞ but P = 0.51) from receiving the same color intensity as high-count TFs with equivalent directionality (e.g., 47/0, P < 0.0001). Blue indicates significant negative bias (TF loses assignment toward positive PC end); red indicates significant positive bias (TF gains assignment toward positive PC end). Rows use the same three-zone PC2-derived TF order as Panel A; the leftmost column is PC2, with remaining columns ordered by hierarchical clustering (Ward’s D2 method on Euclidean distances) of their mutual similarity. A colored annotation strip between the panels indicates TF functional class (HIF-family, constitutive, NF-κB/immune, tumor suppressor, NFY/metabolic, other). The shared row order reveals that TFs which lose assignment on PC2 (upper rows: SP1, SP3, RELB, ETS1) also lose on PC1, PC5, PC8, and PC9 (blue cells) but *gain* assignment on PC6 (red cells), while TFs that gain assignment on PC2 (lower rows: HIF1A, HIF3A, EPAS1, ARNT) show the mirror pattern. TFs in the middle placeholder zone, absent from the PC2 transition, may show strong directional bias on other axes, confirming that TF participation in switching is axis-specific rather than genome-wide. This cross-axis sign inversion, visible as color reversals within the same row, demonstrates that regulatory architecture evolution operates through modular, axis-specific reorganization of a shared TF toolkit rather than fixed TF hierarchies, and supports the conclusion that HIF-compatible regulatory architecture (PC6) and dominant HIF-family assignment (PC9) are evolutionarily separable dimensions. All *P*-values are two-sided Fisher’s exact tests with BH correction within each PC axis.

The cell cycle regulator CCND2 emerges as the principal hub gene (28 switches, 58.8-fold enriched, Table ED3), switching from constitutive regulators (SP1, HNF4A, RELB) toward bats and to HIF machinery (HIF1A, HIF3A, ARNT, NRF1) toward ground squirrels, consistent with divergent cell-cycle arrest mechanisms during torpor (Fig. 2b). Similarly, TSC2, the mTOR inhibitor, switches from constitutive assignment (HNF4A, RELB, SP1) toward bats to HIF/NRF1 assignment toward ground squirrels (8 switches, 15.1-fold enriched, Table ED3), linking oxygen-sensing assignment to growth suppression during hibernation (Fig. 2b). SOCS1 (12 switches, 25.2-fold enriched, Table ED3) likewise switches from constitutive to HIF1A-mediated control, coupling oxygen sensing to immune suppression during hibernation (Fig. 2b). The regulatory divergence between bats and ground squirrels at these shared hub genes provides a mechanistic explanation for why pan-placental convergence in torpor is absent at the protein-coding level^22,23^: regulatory architecture converges where sequence does not, consistent with the network-level emergence of TF assignment configurations from individually labile binding sites.

Although bats indicate HIF-compatible regulatory architecture (occupying -PC6 space), they contrast with ground squirrels by demonstrating constitutive/non-HIF TF-family assignment (SP1, ETS1, RELB) rather than HIF-family assignment. The co-occurrence of HIF-compatible architecture (-PC6) with constitutive assignment (-PC9) suggests that bats evolved a dual-layered architecture (Fig. 1g) where oxygen-sensing genes maintain HIF-compatible binding structure while delegating primary TF assignment to constitutive factors. This architecture may constitute the mechanistic basis of bats’ flight-driven innovation, allowing flexible switching between constitutive and oxygen-responsive regulatory control without permanently committing to either. By maintaining high-capacity oxidative metabolism via SP1/ETS1 assignment (-PC9) and NFY/HIF binding site infrastructure (-PC6), bats uniquely combine sustained aerobic capacity for high-activity flight with the ability to rapidly transition to metabolic depression during daily torpor. That Myotis species, which possess the highest-activity flight mode and daily enter torpor, penetrate deepest into −PC6/−PC9 space supports this interpretation (Fig. 1g).

### 4. Size-dependent cancer resistance and Peto’s paradox

Peto’s paradox describes the absence of correlation between body size and cancer incidence across species. Although individual mechanisms (TP53 copy-number expansion in elephants^24^, high-molecular-mass hyaluronan in naked mole-rats^25^, and telomere length constraints^26^) have been identified in specific lineages, a unifying framework explaining how cancer resistance scales with body size and ecology across mammals has remained elusive. The joint PC1-PC9 space offers a regulatory-level contribution towards resolving the paradox by revealing that cancer suppression covaries with a bivariate property of regulatory architecture, not a uniform genomic solution scaled to body size (Fig. 1h, Table ED2).

PC1 separates HIF-responsive architecture (HIF1A, SP1, ELK1) from tumor suppressor enrichment (TP53, IRF1, SMAD4, REST). PC9 introduces a second dimension separating hibernating ground squirrels from hibernating vesper bats. The positive end is defined by full HIF machinery (HIF3A, EPAS1, HIF1A and ARNT), while the negative end is characterized by constitutive regulation via SP1, ETS1, and RELB. Together, these two axes create a regulatory space in which cancer vulnerability is determined by quadrant position: because HIF pathway activation, while adaptive for acute hypoxia, can become oncogenic when chronically unregulated, promoting glycolytic metabolism and VEGF-mediated angiogenesis, species lacking both tumor suppressor enrichment (PC1-) and HIF-complex assignment (PC9-) are predicted to be most cancer-vulnerable, while species combining both protective layers (PC1+, PC9+) are predicted to be most cancer-resistant (Fig. 1h).

TF switching confirms these strategies operate through systematic rewiring. The dominant PC1 switch is SP1 to SMAD4 (8 occurrences, 8.1-fold enriched, Table ED5), representing transition from vascular responsiveness to tumor suppressor-mediated growth arrest (Fig. 2c). Along PC9, dominant switches are SP1 to ARNT (6 occurrences, 14.1-fold enriched, Table ED5) and HNF4A to HIF1A (5 occurrences, 8.5-fold enriched, Table ED5), reflecting ground squirrels’ adoption of HIF regulation during torpor while bats maintain constitutive control (Fig. 2b). This adoption extends across all three HIF-α isoforms: HIF1A (5 negative, 30 positive switches, P = 1.2×10⁻□), EPAS1 (0 negative, 10 positive, P = 0.002), and HIF3A (0 negative, 20 positive, P < 0.0001), together contributing 60 positive against only 5 negative HIF-α switching events (Fig. 4b, Table ED4). The complete unidirectionality of EPAS1 and HIF3A, combined with the near-complete bias of HIF1A, indicates that ground squirrels deploy the full HIF-α repertoire as a coordinated HIF-assignment module, consistent with a self-regulating architecture in which HIF3A/IPAS negative feedback may limit pathological HIF overactivation during repeated torpor-arousal cycles.

The same hub genes (CCND2, TSC2) identified in §3 also calibrate cancer protection, while WNT1 functions along PC1 as the dominant hub whose shift from HIF-pathway assignment in ferungulates to TP53/immune-surveillance assignment in myomorphs redirects this pleiotropic signaling node from direct HIF-pathway control toward an integrated immune-surveillance and tumor-suppressor program that enforces proliferation checkpoints (Fig. 2b,c, ED1b, Table ED3).

This bivariate regulatory framework of cancer vulnerability generates testable predictions for individual species with regards to their cancer protection (Table 1, Fig. 1h). The domestic ferret (*Mustela putorius furo*), which has the highest documented neoplasia prevalence among mammals (63% of necropsied zoo individuals)^27^, occupies deep negative PC1 (−28.8) and moderate negative PC9 (−4.53), placing it in the HIF-responsive, tumor suppressor-depleted quadrant lacking HIF-complex protective regulation. Conversely, the naked mole-rat (*Heterocephalus glaber*), a fossorial rodent that does not hibernate but experiences chronic hypoxia (1.5-8% O_2_) in its burrows and exhibits constitutive HIF1A expression even under atmospheric normoxia, occupies positive PC1 (+15.46) and positive PC9 (+22.4), squarely in the tumor suppressor-enriched, HIF-complex-governed quadrant. Its near-complete cancer resistance, recently shown to require simultaneous loss of both p53 and pRb for tumor initiation^28^, places it in the predicted protective quadrant of this regulatory framework: TP53/IRF1/SMAD4 regulatory emphasis (PC1+) combined with coordinated HIF-complex assignment (PC9+). Three phylogenetically and ecologically disparate lineages independently converge in this protective quadrant: the Damaraland mole-rat (*Fukomys damarensis*; PC1 = +26.06, PC9 = +31.61), which shares the fossorial hypoxic niche; elephants (*Loxodonta africana*; PC1 = +3.14, PC9 = +14.15), whose body mass exceeds the naked mole-rat’s by five orders of magnitude yet which exhibit comparably low cancer mortality^27^; and hibernating ground squirrels (PC1 = +12-20, PC9 = +32-51), which face severe torpor-induced hypoxia. This reinforces that animal cancer risk is not solely a function of cell number but is substantially shaped by position in the bivariate PC1-PC9 regulatory space, with species-specific combinations of tumor suppressor investment and HIF pathway control calibrated to life history, body size, and ecological oxygen demands.

**Table 1.**
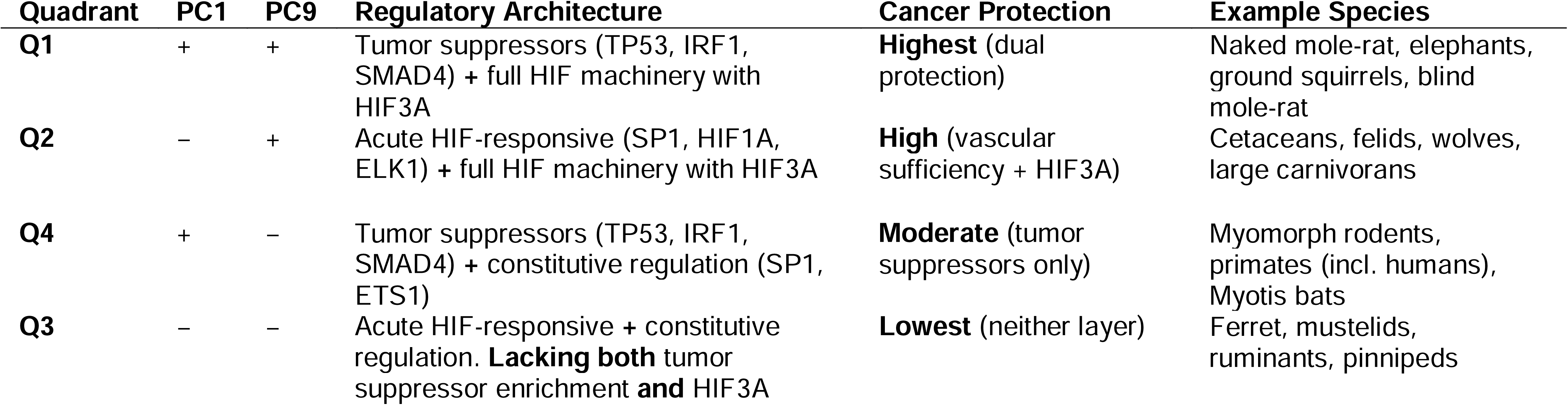
Bivariate regulatory architecture generates predictions for cancer protection across mammals. The four quadrants of the PC1-PC9 regulatory space defined by tumor suppressor investment (PC1) and HIF-complex assignment (PC9) generate distinct predictions for relative cancer protection. Quadrant position reflects the combination of regulatory layers present: tumor suppressor enrichment (TP53, IRF1, SMAD4; positive PC1) and coordinated HIF-complex assignment including HIF3A/IPAS negative feedback (positive PC9). Species with both protective layers (Q1) are predicted most cancer-resistant; species lacking both (Q3) are predicted most cancer-vulnerable. Cancer protection levels are qualitative predictions derived from regulatory architecture position, not direct cancer incidence measurements.

Lineage-specific mechanisms, including somatic TP53 copy-number expansion in elephants^24^, high-molecular-mass hyaluronan production in naked mole-rats^25^, telomere dynamics contributing to cancer resistance through reduced replicative potential in longer-lived species^26^, and variation in DNA repair^29^ and immune surveillance efficiency^30^, are consistent with, and may be downstream consequences of, the regulatory architectures identified here.

### 5. Regulatory architecture lock-in and the pattern of macroevolution

A striking structural feature of the regulatory space described above is that major mammalian radiations occupy discrete, non-overlapping regions across independent regulatory axes (PC2, PC3/PC4, and PC7). Preliminary analyses of metabolism and immunity pathways recover similar clade-level configurations (unpublished data), suggesting the architectures described here reflect a broader multi-dimensional physiological regulatory identity of each radiation.

Two independent lines of evidence support that this pattern reflects punctuated regulatory evolution; architectures established abruptly at key transitions and subsequently inherited with near-neutral within-regime drift, rather than either ongoing stabilizing selection (which would reduce K below 1) or unstructured neutral drift (which would produce K ≈ 1 under Brownian motion). First, Blomberg’s K exceeds 1 on all nine PC axes (p < 0.001; Fig. 1), confirming that within-clade regulatory similarity is substantially greater than neutral drift predicts. Second, multivariate OU shift models (PhyloEM^11^) fitted jointly across all nine PC axes recover discrete, clade-specific optima with phylogenetic half-lives that meet or exceed the total tree height (joint 9-axis model: 231 Myr; tree height 217 Myr). Half-lives at or near the total tree height indicate that the within-regime OU restoring force is weak, which means that architectures do not continuously track their clade’s optimum but instead are established abruptly at major phylogenetic transitions and subsequently drift near-neutrally within clades. The discrete shift structure of the model, not ongoing stabilizing pull, accounts for the K > 1 signal: regulatory architecture is inherited coherently because shifts are rare, not because lineages are continuously corrected back toward a fixed target.

The discrete shift structure documented above, with configurations inherited coherently across all descendants, has consequences that reach beyond oxygen biology.

First, they provide genomic-level evidence for Simpson’s adaptive zones^31^: stable ecological-evolutionary configurations within which lineages diversify but between which transitions are rare. Adaptive zones have historically been defined morphologically and ecologically, without a molecular substrate. The non-overlapping regulatory quadrants (e.g., inter-quadrant distances of 50-90 units on PC3/PC4) give these zones a molecular identity as distinct configurations of TF assignment over the same conserved gene set, defining each lineage’s metabolic and physiological capabilities and constraints.

Second, the coherence of regulatory architecture within radiations provides a genomic mechanism for evolutionary stasis, the dominant but poorly explained pattern in punctuated equilibrium^32^. Erwin and Davidson^33^ proposed that developmental gene regulatory networks contain deeply conserved subcircuits (“kernels”) that resist modification after formation, but this concept has been restricted to body-plan developmental genes. Our findings extend kernel-like behavior to physiological regulatory networks. Key oxygen-sensing genes (PRKAA1, SLC2A4, RHEBL1, TSC2) are metabolic and signaling genes rather than developmental regulators, yet their regulatory architectures show the same clade-level canalization that Erwin and Davidson described for body-plan specification. The marsupial cluster, for example, twelve species from kangaroos to Tasmanian devils sharing essentially identical regulatory wiring despite >60 million years of ecological diversification, is a concrete genomic instance of the active stasis that Eldredge and Gould^32^ envisioned but could not molecularly characterize (Fig. 4a, 2A). The OU shift models establish that this stasis operates through shift rarity rather than continuous stabilizing pull: once the marsupial regulatory configuration was fixed, it persisted not because lineages were actively corrected back toward an optimum, but because no further architectural shifts occurred. The marsupial cluster’s K value on PC2 (K = 41-60 across window conditions; Supplementary Information Figure S7) is the most extreme in the dataset, quantifying a within-clade coherence that far exceeds any plausible neutral expectation and that persists across 60 million years of ecological diversification.

Third, these findings bridge the fifty-year King-Wilson gap. King and Wilson^3^ proposed that morphological evolution proceeds primarily through regulatory change, but most empirical support has come from microevolutionary comparisons, while comparative genomics have emphasized rapid binding-site turnover^4^ that erodes site-level signal over deep time^6,7^. By shifting from site conservation to regulatory architecture identity, we recover structured adaptive signal across 220 million years, providing macroevolutionary-scale genomic evidence for the King-Wilson hypothesis. This resolution parallels Siepel and colleagues’^34^ demonstration that cis-regulatory codes can be conserved across >500 Myr despite near-total enhancer replacement, implying that regulatory codes rather than individual sequences are the evolutionarily stable unit.

Fourth, regulatory architecture functions as a heritable macroevolutionary unit that constrains accessible physiological states. For example, pinnipeds retain caniform regulatory architecture despite 25 million years of aquatic selection, consistent with Wagner and Altenberg’s^35^ prediction that the genotype-phenotype map itself evolves and constrains later evolution.

These observations fit a hierarchical view of evolution in which properties at higher organizational levels (clade-level regulatory architecture) constrain evolution at lower levels in ways not predictable from individual gene regulatory changes alone^36,37^. The lock-in of marsupial regulatory architecture, constraining ecologically diverse species to a narrow region of regulatory space despite tens of millions of years of selection for diverse ecological niches, exemplifies emergent constraint above the level of individual genes.

Taken together, these results show that mammalian macroevolution proceeds through a punctuated mode in which regulatory architectures are established at major transitions, inherited as coherent units, and subsequently resist modification. This lock-in channels diversification within bounds set by ancestral configurations, a pattern reflected in within-regime half-lives that meet or exceed the tree height across regulatory axes (Fig. 1), indicating attraction too weak to erase inherited configurations. While this mode has been theorized based on morphological and developmental observations^33,38,39^, the present findings provide the first genomic-level evidence for it across deep evolutionary time, establishing regulatory architecture lock-in as a fundamental structuring force in macroevolution.

## Acknowledgements

AH was supported by an international exchange program between Stony Brook University and the University of Tübingen.

## Notes

### Competing Interest Statement

The authors have declared no competing interest.

