## Supplementary material for "Oxygen-sensing regulatory architecture structures mammalian diversification": Methods

**Materials and Methods**

***Genomic data and regulatory architecture assembly.***

Genomes, genome annotations, and proteomes were obtained for 239 mammalian species (all mammalian RefSeq genomes available from [NCBI](https://www.ncbi.nlm.nih.gov/datasets/genome/) in September 2025). For each gene, when multiple isoforms were present, the longest isoform was selected. Proteins were classified into orthogroups using OrthoFinder v2.5.5^1^. The density of transcription factor binding sites (TFBS) was quantified across 886 vertebrate TF motifs defined in the JASPAR 2026 database^2^ using TFBSTools version 1.46^3^. We applied the same JASPAR position weight matrices across all 239 species, treating them as a reasonable approximation of sequence-encoding binding potential because TF DNA-binding domains and core sequence specificities are highly conserved across mammals^4^. The resulting motif counts were converted to promoter-level frequencies and treated as a reasonable approximation of sequence-encoded binding potential across species. For each gene in each species, we extracted the 500 base pairs immediately 5′ of the annotated start codon (strand-aware) for the selected isoform and scanned both strands for motif matches, and the quantity of each TFBS type recorded at a 60% minimum relative match score. The 500 bp ATG window was selected as the most consistently defined gene-proximal sequence window across the 239 assemblies, while minimizing annotation-dependent noise from more distal sequence. This ATG-proximal design necessarily excludes distal enhancers. Enhancers are important components of the regulatory network, but they are distributed across large and variable genomic distances, often tens to hundreds of kilobases from their targets, in either orientation, and sometimes within introns of neighboring genes, and cannot be confidently assigned to the gene they regulate without locus-specific chromatin-conformation data (e.g., 3C/Hi-C), which are unavailable for all but a handful of the 239 species analyzed here. Incorporating enhancers into a consistent cross-species gene–TF matrix at this phylogenetic scale is therefore not currently tractable. Our analysis should therefore be understood as characterizing ATG-proximal, sequence-encoded regulatory architecture rather than the complete cis-regulatory landscape. Sensitivity analyses across three alternative window widths and two anchor positions confirm robustness of all major results (Supplementary Information; Procrustes r ≥ 0.984, Mantel r ≥ 0.973). Raw TFBS counts were converted to frequency per 1,000 TFBS for each observed upstream sequence *o* and TFBS type *i*:

$$f_{TFBS_{i,o}}=1000\cdot\frac{n_{TFBS_{i,o}}}{\sum_{j=1}^{886} n_{TFBS_{j,o}}}$$

To correct for interspecific GC content variation, 100 random DNA sequences of 500 bp were simulated at each of 101 GC content levels (0–100%) using SimRAD^5^. TFBS frequencies were calculated for each simulated sequence, and a LOESS regression of simulated TFBS frequency as a function of GC content was fitted for each TFBS type. The residual difference between observed TFBS frequencies and the expectation given the GC content of each upstream region yielded GC-corrected frequencies used in all subsequent analyses.

Predicted binding sites were then filtered against experimental evidence: orthogroups were excluded for a given TFBS unless the ReMap 2022 database^6^ provided experimental support for that TF-gene association in the human regulatory atlas (>600 cell and tissue types). The ReMap database is an atlas of approximately 165 million non-redundant binding regions from >11,000 ChIP-seq and DAP-seq experiments across four species (Human, Mouse, Drosophila, *Arabidopsis thaliana*), of which the human regulatory atlas contains the binding regions used as an evidence filter in the present study. Because ReMap integrates across the union of all cell types and tissues studied, the resulting binding landscape captures pan-tissue regulatory potential rather than the chromatin state of any single tissue. ReMap was used only as an evidence filter, reducing false positives but biasing the analysis toward TF–gene relationships represented in existing human datasets, and not as a tissue-specific chromatin map.

*Pruning to Oxygen-Sensing Regulatory Architecture.*

The full genome-wide dataset of gene–TF binding site pairs was pruned to retain only transcription factors and KEGG pathways directly involved in oxygen sensing. Pathways were assigned to a two-tiered classification based on their mechanistic relationship to the HIF oxygen-sensing system. Tier 1 comprised the core HIF-1 signaling pathway (KEGG 04066), the master transcriptional module for hypoxic response. Tier 2 comprised nine pathways with direct metabolic, regulatory, or cooperative relationships to HIF signaling: three direct metabolic outputs whose gene expression is controlled by HIF-1 (glycolysis/gluconeogenesis, TCA cycle, oxidative phosphorylation); three HIF cooperator or target pathways (JAK-STAT, FoxO, AMPK signaling); and three direct HIF regulators (NF-κB, PI3K-Akt, mTOR signaling). This yielded 10 KEGG pathways encompassing 705 genes (after filtering gene-TF pairs for ReMap support).

Transcription factors were similarly assigned to two tiers. Tier 1 (Essential) comprised the four core HIF pathway members: HIF1A, EPAS1 (HIF-2α), ARNT (HIF-1β), and HIF3A. Tier 2 (Direct regulators/cooperators) comprised 30 transcription factors with experimentally established direct regulatory or cooperative relationships to the HIF pathway, including the NF-κB family (NFKB1, RELA, REL, RELB, NFKB2), SP family (SP1, SP3), ETS-related factors (ELK1, ETS1), STAT factors (STAT1, STAT3), NF-Y complex (NFYA, NFYB, NFYC), SMAD factors (SMAD3, SMAD4), bHLH factors (USF1, USF2, HES1), and additional regulators (NFE2L2, BACH1, REST, HNF4A, FOXO3, YY1, E2F1, IRF1, IRF3, NRF1, TP53). Only Tier 1 and Tier 2 TFs (34 total) and Tier 1 and Tier 2 KEGG pathways (10 total) were retained for subsequent analyses. The promoter-level frequencies for the intersection of these 34 TFs with 705 pathway genes across 239 species produced the gene–TF regulatory architecture matrix analyzed in all subsequent steps.

***Principal Component Analysis.***

*Missing Data Imputation and Phylogenetic Signal.*

A small proportion of gene–TF pairs (2.53%) had missing values due to the absence of orthologous genes in particular species. Missing values were estimated using the function phylo_impute_BM, which imputes missing entries under a Brownian motion model of evolution along the phylogeny. To verify that imputation did not introduce systematic bias, Blomberg's *K^7^* was calculated for all 12,919 traits both with and without imputed values. The phylogenetic signal of the complete (non-imputed) data was *K* = 0.177. Comparison between imputed and non-imputed *K* values showed near-perfect concordance (Pearson *r* = 0.9959; Spearman *r* = 0.9974), with a negligible mean difference (mean Δ*K* = 0.0031; median Δ*K* = 0.0013) and only 63 of 12,919 traits (0.49%) showing |Δ*K*| > 0.1. These results confirm that Brownian motion imputation of 2.53% missing data does not materially alter phylogenetic signal estimates. Blomberg's *K* values reported here reflect the raw gene–TF trait matrix and are used solely to validate imputation fidelity; phylogenetic signal of the summarized regulatory architecture is reported separately on PC axis scores (see *Phylogenetic Signal of PC Axes*, below).

*Choice of Standard PCA over Phylogenetic PCA.*

Standard principal component analysis was applied to the imputed species × gene–TF trait matrix (239 species × 12,919 traits). The case for standard over phylogenetic PCA (pPCA) in this setting is grounded in the fact that, in the presence of evolutionary shifts, pPCA preprocessing is actively counterproductive because it removes the phylogenetically structured signal that constitutes the primary biological signal under investigation. Five independent lines of evidence support this choice.

First, the weak phylogenetic signal across traits (mean Blomberg's *K* = 0.177) indicates that phylogenetic covariances contribute minimally to the structure of the data matrix. The statistical artifacts that pPCA is designed to correct, inflation of early-burst patterns and phylogenetic sorting into the leading principal components, occur primarily when phylogenetic signal is strong (*K* approaching 1.0)^8^. With *K* = 0.177, the data approximate a star phylogeny for which PCA and pPCA converge to the same solution^8,9^.

Second, Bastide et al.^10^ proved formally that pPCA fails to decorrelate traits in the presence of evolutionary shifts; precisely the scenario this study investigates. The pPCA estimator of the trait covariance matrix is biased whenever shifts are present: E[**R̂**] = **R** + **B**, where the additive bias term **B** = (n−1)⁻¹**G**ᵀ**C**⁻¹**G** (with **G** being the matrix of group mean deviations induced by the shifts) is identically zero only when no shifts exist. Because **R̂** is biased, the PC scores produced by pPCA remain correlated with one another; defeating the entire purpose of the preprocessing step. This problem is exacerbated under an OU process: Bastide et al. note explicitly that although bias in **R̂** is moderate when shifts occur on long branches (where the bias term **B** is small relative to **R**), it becomes large, comparable in magnitude to **R** itself, when shifts occur on short internal branches, as expected for clade-defining transitions in mammalian regulatory architecture. The authors conclude: "we show that the standard practice of decorrelating traits using phylogenetic principal component analysis (pPCA) before using a method designed for independent traits can be misleading in the presence of shifts.

Third, Bastide et al.^10^ demonstrated numerically that pPCA preprocessing actively harms shift detection accuracy when the number or amplitude of shifts is large. In simulations with correlated traits (between-trait correlation ρ_d = 0.4), pPCA is neutral at best when only a few small shifts are present, but degrades performance materially as shifts become more numerous or larger; precisely the conditions expected across a 239-species mammalian phylogeny spanning 217 Myr. Furthermore, in their Anolis lizard case study, Bastide et al. found that the number of shifts recovered by a pPCA-based method depended entirely on the arbitrary choice of how many pPC axes to retain (ranging from 5 to 20 shifts, the maximum allowed), and that even after pPCA transformation, the retained axes had off-diagonal correlations up to |0.29|, confirming that pPCA does not achieve the independence it is intended to produce.

Fourth, the research question explicitly concerns phylogenetically structured adaptive evolution: the diversification of oxygen-sensing regulatory architecture across mammalian radiations. Polly et al.^9^ caution that "when the phylogenetic changes are adaptive, then environmental components of variation are identical to phylogenetic components. Removing phylogenetic correlations from data will remove precisely the component of shape variation that is relevant to adaptation." Applying pPCA would therefore risk eliminating the adaptive phylogenetic signal targeted by the analysis.

Fifth, the downstream analytical framework (PhyloEM, scOU model) was specifically designed to render pPCA preprocessing unnecessary. Bastide et al.^10^ state that the scOU model "spares us from doing a preliminary pPCA" by accommodating residual between-trait correlations through an unconstrained full diffusion matrix **R**. Standard PCA axes, while not phylogenetically orthogonalised, are analytically orthogonal in the sample space; their residual phylogenetic covariances are handled explicitly by **R** in the scOU likelihood, making the trait-independence assumption that pPCA attempts to enforce both unnecessary and potentially harmful for this specific downstream application. Uyeda et al.^8^ additionally demonstrated that when the leading eigenvector explains a large share of variation, PC scores are not appreciably distorted, and that multivariate datasets with low effective dimensionality show no systematic distortion of contrasts through time.

A sensitivity analysis addressing concerns about analytical choices more generally is presented in the Supplementary Information. The PCA configuration (biological interpretation of axes, phylogenetic signal, quadrant structure) is robust to scanning window width and anchor position across four conditions (Procrustes r ≥ 0.984, Mantel r ≥ 0.973 across ATG 500bp, ATG 1000bp, ATG 1500bp, and TSS 500bp windows), confirming that the results are not artefacts of the primary methodological choices. The choice between standard and phylogenetic PCA cannot be addressed by a comparable sensitivity analysis because the two approaches are designed to detect different properties of the data: pPCA removes the between-clade phylogenetic variance that, under the shifts model documented here, constitutes the adaptive signal, making the approaches non-comparable rather than exchangeable.

*TF-Specific Loading Strengths.*

Because the raw PCA loadings matrix contains one value per gene–TF pair per PC axis, loadings were aggregated to characterize the contribution of each transcription factor to each axis of variation. Each row of the loadings matrix was labelled as a gene–TF pair (e.g., "PRKAA1.SP1"). The TF identity was extracted from each label, and the mean loading across all gene–TF pairs sharing the same TF was calculated for each PC axis (Table ED6), yielding a TF centroid loading per axis. The overall strength of each TF's contribution to a given bivariate PC space was then calculated as the Euclidean norm of its centroid loadings across the selected axes:

$$\text{TF strength}=\sqrt{\sum_{k\in\text{axes}} \bar{L}_{TF,k}^{2}}$$

where $\bar{L}_{TF,k}$ is the mean loading for a given TF on PC axis *k*. For individual axes, signed strengths were calculated to preserve directionality, defined as $\text{sign}(\bar{L}_{TF,k})\times|\bar{L}_{TF,k}|$. These TF strengths indicate which transcription factors dominate each axis of regulatory variation (for example, whether an axis primarily separates species by HIF-family versus SP-family assignment) and their signs indicate the direction of that dominance (Fig. 4b, Table ED6).

*Phylogenetic signal of PC axes*

To characterize the degree to which the PCA-summarized regulatory architecture reflects shared evolutionary history, phylogenetic signal was quantified on the PC scores in two complementary ways.

For each individual PC axis (PC1–PC9), univariate phylogenetic signal was assessed using Blomberg's *K^7^*, implemented in the phytools R package^11^ (function phylosig, method = "K") with significance evaluated against a null distribution of 1,000 random tip-label permutations. Blomberg's *K* equals 1 under a Brownian motion model, is greater than 1 when closely related species are more similar than expected under BM (phylogenetic conservatism), and is less than 1 when trait similarity decays faster than expected under BM (phylogenetic lability). Statistical significance (*P* < 0.05) indicates that the observed *K* value is unlikely under the null hypothesis of no phylogenetic signal.

For the joint 9-dimensional space (PC1–PC9 simultaneously), multivariate phylogenetic signal was assessed using *K*_mult, the multivariate extension of Blomberg's *K* implemented in the geomorph R package^12^ (function physignal), with significance evaluated against 1,000 random permutations. *K*_mult summarizes the degree to which the full multivariate regulatory architecture covaries with phylogenetic relatedness, and provides a single overall measure of phylogenetic structure in the nine-axis trait space.

Blomberg's *K* (univariate) and *K*_mult (multivariate) were computed using the same dated phylogeny (217 Myr) used for all other comparative analyses. Results are reported alongside phylogenetic half-life estimates in Fig. 1.

***TF Switching Analysis.***

A complementary switching analysis was developed to identify, for each gene, which transcription factors lose assignment and which gain it along each axis of variation, providing gene-level wiring diagrams of regulatory evolution. Two perspectives on switching were implemented: TF-centric switching (which TF pairs co-occur as directional switches across genes?) and gene-centric switching (which genes are regulatory switching hubs?) (Fig. 2,3, Tables ED3, ED5).

I*dentification of Switching Events.*

For each PC axis, gene–TF pairs were ranked by their loading values and the top 2.5% most positive and bottom 2.5% most negative loadings were identified. A switching event was defined when the same gene appeared in both tails with different TF partners: the gene was paired with TF_pos in the positive tail and TF_neg in the negative tail. This indicates that along the axis of variation, the gene’s regulatory identity shifts from TF_neg assignment (at one end of the species distribution) to TF_pos assignment (at the other end). The magnitude of each switch was quantified as the absolute difference (delta) between the positive and negative loading values.

The 2.5% tail threshold was chosen to identify the most extreme loading values while retaining sufficient gene-TF pairs per axis for statistical enrichment testing; it represents a standard extreme-quantile approach in loading-based analyses. The qualitative pattern of switching events (which TF pairs switch, which hub genes are involved, and the directionality of enrichment) is robust to threshold choices in the range 1–5% (assessed by examining the concordance of top-ranked switching pairs across threshold values in preliminary analyses). Stricter thresholds (1%) reduce the number of detectable switches and lower statistical power without altering the identity of the most enriched pairs; more permissive thresholds (5%) increase noise without changing principal findings.

*TF-Centric Switching and Enrichment.*

TF-centric switching quantifies how often a specific directed TF pair (TF_neg → TF_pos) recurs across different genes on a given PC axis (Fig. 2c, 3b, Tables ED5). For each TF pair on each PC, the number of genes exhibiting that specific switch was counted. Statistical significance was assessed using a hypergeometric test for co-occurrence. The hypergeometric framework tests whether genes paired with TF_neg in the negative tail and with TF_pos in the positive tail overlap more than expected by chance. Specifically, for each TF pair, the universe N was defined as the total number of unique genes, K as genes appearing with TF_neg in the negative tail, n as genes appearing with TF_pos in the positive tail, and the observed overlap X as the number of genes exhibiting both conditions. The enrichment was calculated as X / (Kn/N), and the P-value was obtained from the upper tail of the hypergeometric distribution: P(X ≥ observed). Odds ratios and Fisher's exact test P-values were also computed from the resulting 2 × 2 contingency table. P-values were corrected for multiple testing within each PC axis using the Benjamini–Hochberg (BH) procedure (Table ED5).

*Gene-Centric Switching and Enrichment.*

Gene-centric switching quantifies whether individual genes serve as regulatory hubs by participating in more switching events than expected by chance (Fig. 2a,b, 3a, Table ED3). For each gene on each PC axis, the total number of distinct TF_neg–TF_pos combinations (i.e., distinct switches) was counted. To assess whether the observed number of switches was enriched, a global switch probability was first estimated across all genes on that PC axis. For each gene, the number of TFs appearing in the positive tail (n_TF_pos) and negative tail (n_TF_neg) were counted, and the total potential switches were defined as n_TF_pos × n_TF_neg. The global switch probability p̂ was estimated as the ratio of total observed switches to total potential switches across all genes with TFs in both tails. Statistical significance for each gene was then assessed through Monte Carlo simulation (n = 10,000 iterations). In each iteration, each of the gene's m TF partners was independently assigned to the positive tail (with probability q_pos), negative tail (probability q_neg), or neither, where q_pos and q_neg were the global tail-entry frequencies for that PC axis. The number of simulated switches was recorded, and the empirical P-value was calculated as the proportion of simulations yielding at least as many switches as observed. Enrichment was computed as observed switches divided by the mean of simulated switches. P-values were BH-corrected within each PC axis (Table ED3).

*TF Tail Bias.*

An additional analysis tested whether individual transcription factors show directional bias in their switching involvement. For each TF on each PC axis, the number of switching events in which it appeared on the positive side versus the negative side was counted (Fig. 4, Table ED4). A 2 × 2 contingency table was constructed contrasting positive-side and negative-side event counts for that TF against the corresponding counts for all other TFs, and Fisher's exact test was applied to obtain an odds ratio and two-sided P-value. TFs were classified as "positive-biased" (odds ratio > 1), "negative-biased" (odds ratio < 1), or unbiased. P-values were BH-corrected within each PC axis. This analysis reveals whether a TF systematically gains assignment (positive bias) or loses assignment (negative bias) along a given axis of regulatory variation (Table ED4).

***Ornstein–Uhlenbeck Regime-Shift Analysis.***

*Theoretical Framework*

To formally identify evolutionary shifts in regulatory architecture across the mammalian phylogeny, multivariate Ornstein–Uhlenbeck shift models were fitted jointly to species scores across all nine PC axes simultaneously using the scalar OU (scOU) process.

The scOU process is obtained from the general multivariate OU SDE dX_t = A(β − X_t) dt + R dW_t, where A is the selection strength matrix, β the primary optimum, and R the drift rate matrix, by restricting A to a scalar form A = αI_p. This single shared selection strength α simplifies computations while allowing a full unconstrained p × p diffusion matrix R to capture residual between-trait covariances. The conditional distribution of the trait vector X_i at node i given its parent X_pa(i) is then:

X_i | X_pa(i) ~ N( e^(−αt_i) X_pa(i) + (1 − e^(−αt_i)) β_i, (1/(2α))(1 − e^(−2αt_i)) R )

where β_i = β_pa(i) + Δ_i is the optimum inherited by node i after any shift Δ_i on the branch of length t_i leading from pa(i) to i. The joint distribution of observed tip traits is:

vec(Y) ~ N( vec(T W(α) Δ), R ⊗ F(α) )

where W(α) is a diagonal matrix of regime weighting factors and F(α) is the scaled phylogenetic correlation matrix; this is the likelihood expression that PhyloEM maximizes. The parameter α ≥ 0 controls the strength of the deterministic pull toward the optimum; α ≈ 0 reduces the process to multivariate Brownian motion. The phylogenetic half-life, t₁/₂ = ln(2)/α, gives the time required for a lineage displaced from its regime optimum to return halfway toward it, and provides the primary metric for characterizing the mode of evolutionary change (see Results).

The OU model with multiple optima was first proposed by Hansen^13^ and developed by Butler and King ^14^. As Butler and King^14^ established, the selective optima of the OU process provide an explicit mathematical formalization of Simpson's^15^ concept of adaptive zones; the stable ecological-evolutionary configurations within which lineages diversify but between which transitions are rare. In the multi-optima OU model, each optimum $\beta_{k}$ represents a distinct adaptive zone, and each shift in $\beta$ along a branch of the phylogeny represents a transition between zones. This provides a direct quantitative link between the macroevolutionary pattern of clade-level regulatory architecture lock-in observed in the PCA (where major radiations occupy discrete, non-overlapping regions of regulatory space) and a formal evolutionary model in which these configurations are established by discrete shifts in the optimal state at specific phylogenetic transitions, and subsequently maintained through near-neutral within-regime drift rather than by ongoing stabilizing selection.

As Khabbazian et al.^16^ noted, even when the optimum value is estimated to be constant within a given clade, this value may represent a broad adaptive zone around which the true local optimum fluctuates, making it prudent to interpret shifts in $\beta$ as transitions between adaptive zones rather than literal estimates of a single optimal phenotype. Hansen^13^ proposed that the macroevolutionary OU process operates on a timescale distinct from within-population microevolutionary dynamics, with the optimum $\beta$ reflecting the balance of multiple conflicting selective demands, and with shifts corresponding to Simpson's "quantum evolution": rare, rapid transitions between adaptive zones. The OU framework thus provides both a statistical tool for detecting regime shifts and a conceptual bridge between molecular regulatory data and classical macroevolutionary theory.

*Implementation via PhyloEM.*

Regime shifts were detected using the PhylogeneticEM package (PhyloEM)^10^, which implements a phylogenetic Expectation–Maximization (EM) algorithm to simultaneously estimate the number, phylogenetic position, and magnitude of shifts in trait optima without prespecified hypotheses^17^. Unlike penalized-regression approaches, PhyloEM treats the locations and magnitudes of shifts as latent variables recovered through iterative E and M steps, enabling exact likelihood computation under the multivariate scOU model. Critically, the scOU framework is specifically designed to handle correlations among input traits through the unconstrained diffusion matrix **R**, making any prior pPCA decorrelation step unnecessary and, in the presence of shifts, actively counterproductive (see Choice of Standard PCA over Phylogenetic PCA, above; Bastide et al.^10^).

The EM algorithm alternates between two steps. The E-step performs ancestral trait reconstruction at all internal nodes conditional on the observed tip data, using an upward–downward pruning recursion on the phylogeny (upward pass: compute conditional means and variances from leaves to root; downward pass: update ancestral estimates conditioning on the full taxon set). The M-step uses the reconstructed ancestral values to identify shift positions via a lasso penalty on whitened ancestral increments, and updates the diffusion matrix **R** in closed form. The selection strength α is not estimated within the EM loop but is instead treated as a fixed outer parameter optimized over a grid of values; a complete EM algorithm is run separately for each α on the grid, and the (K, α) combination yielding the best model selection criterion is adopted. The EM algorithm is initialized using a LASSO-based sparse regression on the whitened data to obtain candidate shift positions and magnitudes, following the approach described in Bastide et al.^10,17^.

Because the EM algorithm maximizes the likelihood for a fixed number of shifts K, the procedure was run across a grid of 20 α values, equally spaced on a log scale and calibrated to span t₁/₂ values from approximately 1% to 300% of total tree height (217 Myr), covering the range from near-star-tree behavior (t₁/₂ ≪ tree height, strong selection) to near-Brownian motion (t₁/₂ ≫ tree height, weak selection), and across all values of K from 0 to K_max = 15. The upper limit was set to K_max = 15, a conservative cap below the package default (√n + 5 ≈ 21 for n = 239) chosen to reduce computational load while remaining above the number of major mammalian radiations expected a priori.

Model selection, the choice of optimal (K, α) combination, was performed using three penalized criteria implemented in PhyloEM. The primary criterion, LINselect, is a theoretically grounded penalised least-squares criterion^10,18^ whose penalty accounts explicitly for the identifiability of shift positions on the tree (the number of parsimonious, non-equivalent shift configurations C(2n−2−K, K) for a fully resolved tree). DDSE and Djump are slope-heuristic alternatives^19,20^ that infer the penalty empirically from the profile of the criterion across K values rather than from theoretical bounds. For the joint 9-axis analysis (p = 9), Djump was adopted as the primary criterion because the multivariate residual tr[R̂(K, α̂)] aggregates signal across all nine traits, and Djump's response to abrupt profile discontinuities is more robust to dimensionality-driven smoothing of the criterion than LINselect's theory-derived penalty in high-dimensional settings. For the nine univariate per-axis analyses (p = 1), LINselect is not defined: the BGHml penalty term underlying LINselect requires the estimation of a between-trait penalty constant that is degenerate for a single trait. Djump was therefore used as the primary criterion for all univariate fits. Sensitivity of univariate half-life estimates to the choice of criterion was assessed by comparing Djump and DDSE results across all nine axes; estimates were equivalent across both criteria, confirming robustness of the reported values. Concordance across all applicable criteria was evaluated as a general sensitivity check.

The root state was modelled as a random draw from the OU stationary distribution (random.root = TRUE, stationary.root = TRUE), appropriate for a deep mammalian phylogeny spanning 217 Myr in which the ancestral regulatory state is unknown.

*Application to PC Scores.*

PhyloEM was applied in two complementary configurations. First, a joint analysis was run across all nine PC axes simultaneously, fitting the multivariate scOU model to a 239 x 9 trait matrix (rows = species, columns = PC axes). Because PhyloEM accommodates residual covariances among axes through the unconstrained diffusion matrix **R**, and because pPCA decorrelation is both unnecessary and biased in the presence of shifts^10^, the PC scores were supplied directly to PhyloEM without further preprocessing. This joint analysis captures covariances among PC axes within each regime and estimates shift vectors in the full 9-dimensional regulatory architecture space. Second, nine univariate analyses were run, one per PC axis, each fitting the scOU model to a single trait vector of length 239. These univariate fits complement the joint K_mult signal estimate with univariate Blomberg's K values and are not used for shift detection (which requires the full multivariate configuration to recover co-directional shifts). On individual axes with K > 1, the scalar α converges to the lower bound of the search grid, so the per-axis half-lives are boundary values reflecting uniformly weak within-regime attraction and are not interpreted as discriminating tempos among axes.

In both configurations, each identified shift partitions the 239 species into groups (regimes) sharing a common regulatory optimum. Each regime corresponds to a clade or set of clades inheriting a distinct configuration of TF assignment. The joint shift configuration across all nine axes defines the complete regulatory identity of each mammalian radiation and provides the molecular substrate for Simpson's adaptive zones.

The scalar selection strength parameter α and the associated phylogenetic half-life $t_{1/2}=\ln(2)/\alpha$ were extracted from the best-fitting (K, α) combination for each analysis configuration. For the joint 9-axis analysis, Djump was the primary criterion for all analyses. In all configurations, α is estimated in units of 1/Myr (consistent with the Myr-scaled branch lengths of the input chronogram), so that $t_{1/2}$ is expressed directly in millions of years without further rescaling. Because α converged to the lower bound of the search grid in all configurations, reported half-lives are treated as lower bounds (α indistinguishable from zero) rather than point estimates; the inference of weak within-regime attraction depends on the half-life being long relative to tree height, not on its precise value. Phylogenetic half-life and Blomberg's *K* for the joint 9-axis space and for each individual PC axis are reported in Fig. 1.

The phylogeny was taken from Smaers et al.^21^, who provided a consensus tree from the fully resolved mammalian supertree compiled by Faurby et al.^22^. The tree was confirmed to be ultrametric prior to analysis, with total root-to-tip depth of 217 Myr.

***Software and Reproducibility.***

Orthology inference used OrthoFinder v2.5.5^1^. TFBS scanning and motif matching used TFBSTools v1.46^3^ with JASPAR 2026 position weight matrices^2^. GC content simulation used SimRAD^5^. Experimental validation filtering used ReMap 2022^6^. Phylogenetic signal was calculated using Blomberg's *K^7^*. PCA was performed using the prcomp function in R with centering. OU regime-shift detection and phylogenetic half-life estimation used the PhylogeneticEM package (v. 1.8.1); Djump model selection was adopted as the primary criterion for the joint 9-axis analysis and for all nine univariate per-axis analyses. PhyloEM was run with rescale_OU = TRUE (OU-to-BM tree rescaling for exact likelihood computation) and light_result = FALSE (full ancestral trait imputation retained for diagnostic checks). Univariate Blomberg's *K* was calculated using the phytools package (function phylosig, 1,000 permutations). Multivariate *K*_mult was calculated using the geomorph package (function physignal, 1,000 permutations). All switching analyses, TF strength calculations, and enrichment tests were implemented in custom R functions using parallel computation via the parallel and furrr packages.

1	Emms, D. M. & Kelly, S. OrthoFinder: phylogenetic orthology inference for comparative genomics. *Genome biology* **20**, 238 (2019).

2	Ovek Baydar, D. *et al.* JASPAR 2026: expansion of transcription factor binding profiles and integration of deep learning models. *Nucleic acids research* **54**, D184-D193 (2026).

3	Tan, G. & Lenhard, B. TFBSTools: an R/bioconductor package for transcription factor binding site analysis. *Bioinformatics* **32**, 1555-1556 (2016).

4	Nitta, K. R. *et al.* Conservation of transcription factor binding specificities across 600 million years of bilateria evolution. *elife* **4**, e04837 (2015).

5	Lepais, O. & Weir, J. T. Sim RAD: an R package for simulation‐based prediction of the number of loci expected in RAD seq and similar genotyping by sequencing approaches. *Molecular ecology resources* **14**, 1314-1321 (2014).

6	Hammal, F., De Langen, P., Bergon, A., Lopez, F. & Ballester, B. ReMap 2022: a database of Human, Mouse, Drosophila and Arabidopsis regulatory regions from an integrative analysis of DNA-binding sequencing experiments. *Nucleic acids research* **50**, D316-D325 (2022).

7	Blomberg, S. P., Garland Jr, T. & Ives, A. R. Testing for phylogenetic signal in comparative data: behavioral traits are more labile. *Evolution* **57**, 717-745 (2003).

8	Uyeda, J. C., Caetano, D. S. & Pennell, M. W. Comparative analysis of principal components can be misleading. *Systematic Biology* **64**, 677-689 (2015).

9	Polly, P. D., Lawing, A. M., Fabre, A.-C. & Goswami, A. Phylogenetic principal components analysis and geometric morphometrics. *Hystrix* **24**, 33 (2013).

10	Bastide, P., Ané, C., Robin, S. & Mariadassou, M. Inference of adaptive shifts for multivariate correlated traits. *Systematic Biology* **67**, 662-680 (2018).

11	Revell, L. J. phytools: an R package for phylogenetic comparative biology (and other things). *Methods in Ecology and Evolution* **3**, 217-223 (2012). <https://doi.org/10.1111/j.2041-210X.2011.00169.x>

12	Adams, D. C. & Otárola‐Castillo, E. geomorph: an R package for the collection and analysis of geometric morphometric shape data. *Methods in Ecology and Evolution* **4**, 393-399 (2013).

13	Hansen, T. F. Stabilizing selection and the comparative analysis of adaptation. *Evolution* **51**, 1341-1351 (1997).

14	Butler, M. A. & King, A. A. Phylogenetic comparative analysis: A modeling approach for adaptive evolution. *American Naturalist* **164**, 683-695 (2004).

15	Simpson, G. G. *Tempo and mode in evolution*. (Columbia University Press, 1944).

16	Khabbazian, M., Kriebel, R., Rohe, K. & Ané, C. Fast and accurate detection of evolutionary shifts in Ornstein–Uhlenbeck models. *Methods in Ecology and Evolution* **7**, 811-824 (2016).

17	Bastide, P., Mariadassou, M. & Robin, S. Detection of adaptive shifts on phylogenies by using shifted stochastic processes on a tree. *Journal of the Royal Statistical Society Series B: Statistical Methodology* **79**, 1067-1093 (2017).

18	Baraud, Y., Giraud, C. & Huet, S. Gaussian model selection with an unknown variance. *Annals of Statistics* **37**, 630-672 (2009).

19	Arlot, S. & Massart, P. Data-driven Calibration of Penalties for Least-Squares Regression. *Journal of Machine learning research* **10** (2009).

20	Baudry, J.-P., Maugis, C. & Michel, B. Slope heuristics: overview and implementation. *Statistics and Computing* **22**, 455-470 (2012).

21	Smaers, J. B., Turner, A. H., Gómez-Robles, A. & Sherwood, C. C. A cerebellar substrate for cognition evolved multiple times independently in mammals. *eLife* **7**, e35696 (2018).

22	Faurby, S. & Svenning, J.-C. A species-level phylogeny of all extant and late Quaternary extinct mammals using a novel heuristic-hierarchical Bayesian approach. *Molecular phylogenetics and evolution* **84**, 14-26 (2015).
