## Supplemental Information for "Oxygen-sensing regulatory architecture structures mammalian diversification"

**Supplementary Information**

### Choice of scanning window and anchor position

#### Overview

All primary analyses in this study were performed on transcription factor binding site (TFBS) counts extracted from the 500 bp region immediately upstream of the start codon (ATG) for each gene across 239 mammalian species (the ATG 500 bp window). To assess whether the biological conclusions depend on this choice, we conducted a sensitivity analysis comparing the ATG 500 bp solution against three alternatives: a 1,000 bp ATG-anchored window (ATG 1000 bp), a 1,500 bp ATG-anchored window (ATG 1500 bp), and a 500 bp window anchored at the transcription start site (TSS 500 bp). The ATG-window comparisons test sensitivity to window width; the ATG vs. TSS comparison tests sensitivity to anchor position.

#### Part 1: Justification for the ATG 500 bp Window

##### The TSS as the biological ideal

Promoters are canonically defined relative to the transcription start site, and functional promoter-proximal transcription factor binding is concentrated near the TSS^1^. For this reason, the TSS is the biologically natural anchor for investigating regulatory evolution. In well-annotated genomes, a window of 500 bp upstream of the TSS captures the core promoter and the majority of proximal regulatory elements with high fidelity.

We do note that functional TFBS are not exclusively located upstream of the TSS: core promoter elements and TF binding sites downstream of the TSS, including the initiator element (INR), downstream core promoter element (DPE), and binding sites for SP1, NF-Y, and ETS-family factors in the early transcribed region, are well-documented features of mammalian promoters^1^. A window strictly upstream of the TSS therefore does not capture the full complement of promoter-proximal regulatory elements. An ATG-anchored window, by design, samples a region that in many genes spans or overlaps the TSS, and that in genes with short 5' UTRs (where the ATG lies within a few hundred base pairs of the TSS) also captures sequence downstream of the TSS. The ATG window thus encompasses a biologically heterogeneous but functionally relevant promoter-proximal region that is not strictly equivalent to a TSS-upstream window and, in this respect, may more completely represent the full regulatory neighborhood of the gene in species where TSS annotation is reliable.

##### Why ATG was used as the cross-species anchor

In a macroevolutionary analysis spanning 239 mammalian genomes drawn from NCBI assemblies^2^, TSS and corresponding 5’ untranslated region (UTR) annotations are substantially less uniform than coding sequence (CDS) annotations. TSS coordinates depend on full-length transcript evidence (RNA-seq coverage, CAGE data, or manual curation) which varies enormously in depth and quality across assemblies. The CDS start codon (ATG), by contrast, is defined by the protein-coding sequence and is therefore one of the most consistently and reliably annotated features across all 239 species in this dataset.

The practical consequence is severe: across the 239 NCBI mammalian species used here, species-level median TSS-to-ATG distances range from 0 to 223 bp. Eight species have a median of 0 bp, 58 species have a first quartile of 0 bp, and 101 species have third quartiles of at least 1,000 bp (Figure S1). These values are incompatible with a uniform biological distribution of mammalian 5′ features; they are exactly what would be expected if transcript-end annotation quality varies substantially among assemblies. Using the TSS as the cross-species anchor would therefore introduce systematic, assembly-quality-driven variation into the comparative matrix; noise that is entirely unrelated to regulatory biology. Current reference resources explicitly acknowledge that transcript models continue to be improved as new long-read transcriptomic data accumulate, and that many non-model species lack or have incomplete annotations upstream of the CDS^2^.

The ATG anchor is therefore not chosen because it is biologically equivalent to the TSS. It is chosen because it is the only anchor that can be applied consistently and reliably across all 239 species.

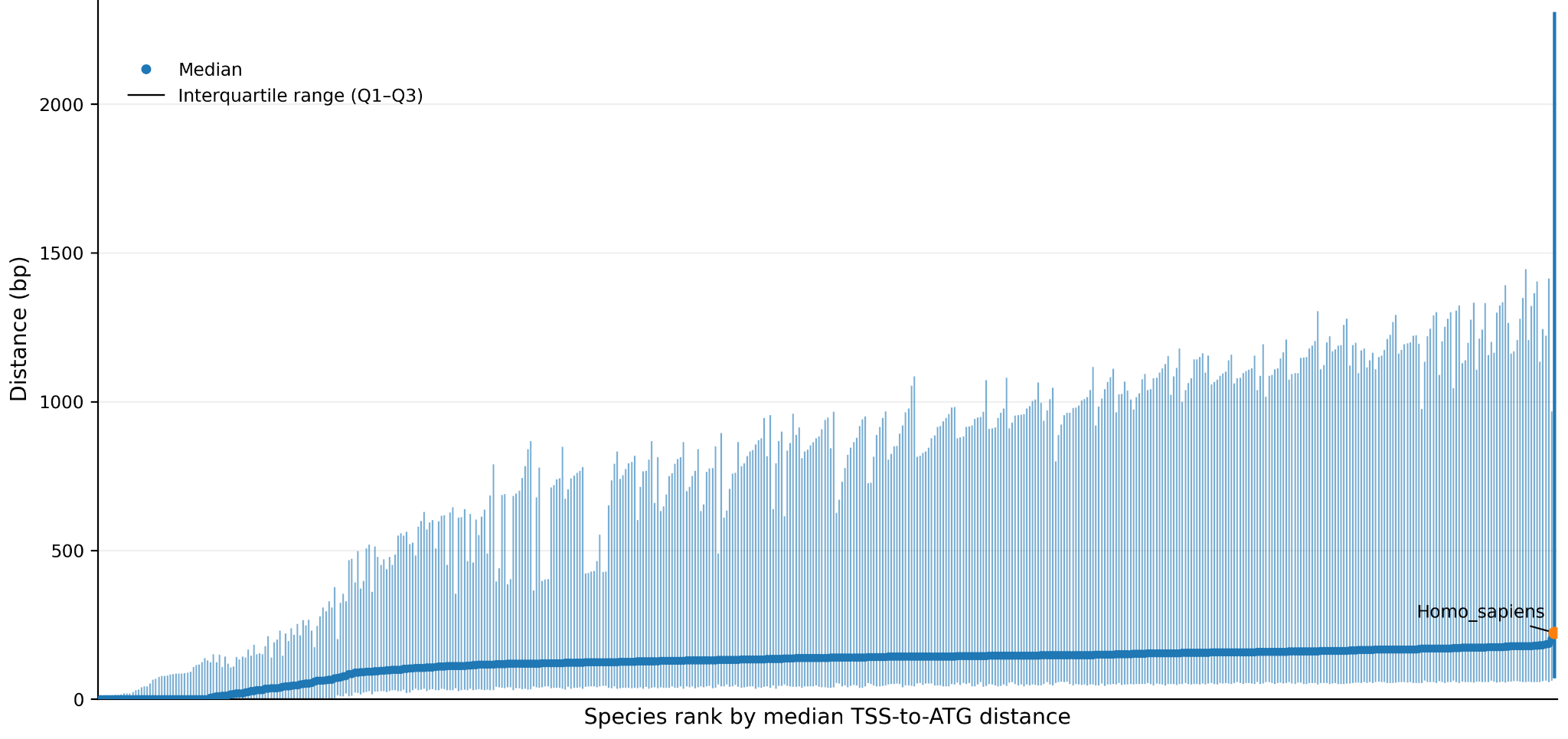

**Figure S1.** Cross-species inconsistency in NCBI TSS-to-ATG summaries. Species are ranked by the median TSS-to-ATG distance; whiskers show the interquartile range (Q1 to Q3). The medians vary from 0 to 223 bp; 8 species have median = 0 bp, 58 have Q1 = 0 bp, and 101 have Q3 ≥ 1000 bp. This argues against treating NCBI TSS coordinates as a uniformly reliable macroevolutionary anchor. Note: 223 bp is the maximum *species-level median* TSS-to-ATG distance; individual gene-level TSS-to-ATG distances within species can be substantially larger, as indicated by the Q3 ≥ 1,000 bp values for 101 species.

##### Quantifying the Cost of the ATG Choice in the Best-Annotated Reference

The cost of using an ATG anchor, in terms of how often the annotated TSS is actually captured, can be estimated most honestly in human, where transcript annotation is the deepest available (including MANE harmonisation between RefSeq and Ensembl/GENCODE) and where the ReMap evidence filter used in this study is itself based^3^.

Among the 705 oxygen-sensing genes used in this analysis, the median human TSS-to-ATG distance is 203 bp, close to the 209 bp genome-wide human median. The annotated TSS lies within the ATG 500 bp window for 477/705 genes (67.7%), within the ATG 1,000 bp window for 533/705 genes (75.6%), and within the ATG 1,500 bp window for 562/705 genes (79.7%) (Figures S2-3). The ATG 500 bp window thus captures the annotated TSS for approximately two-thirds of the oxygen-sensing gene set in the best-annotated reference genome; a meaningful but imperfect approximation.

Approximately 35% of human genes contain at least one 5′ UTR intron^4^. For these genes, the sequence sampled upstream of the ATG includes 5′ UTR and first-intron sequence. Such sequence is not biologically inert, first introns are enriched for regulatory information^5^, but it is not a clean TSS-anchored promoter. This heterogeneity is acknowledged as an inherent limitation of ATG-anchored scanning at macroevolutionary scales.

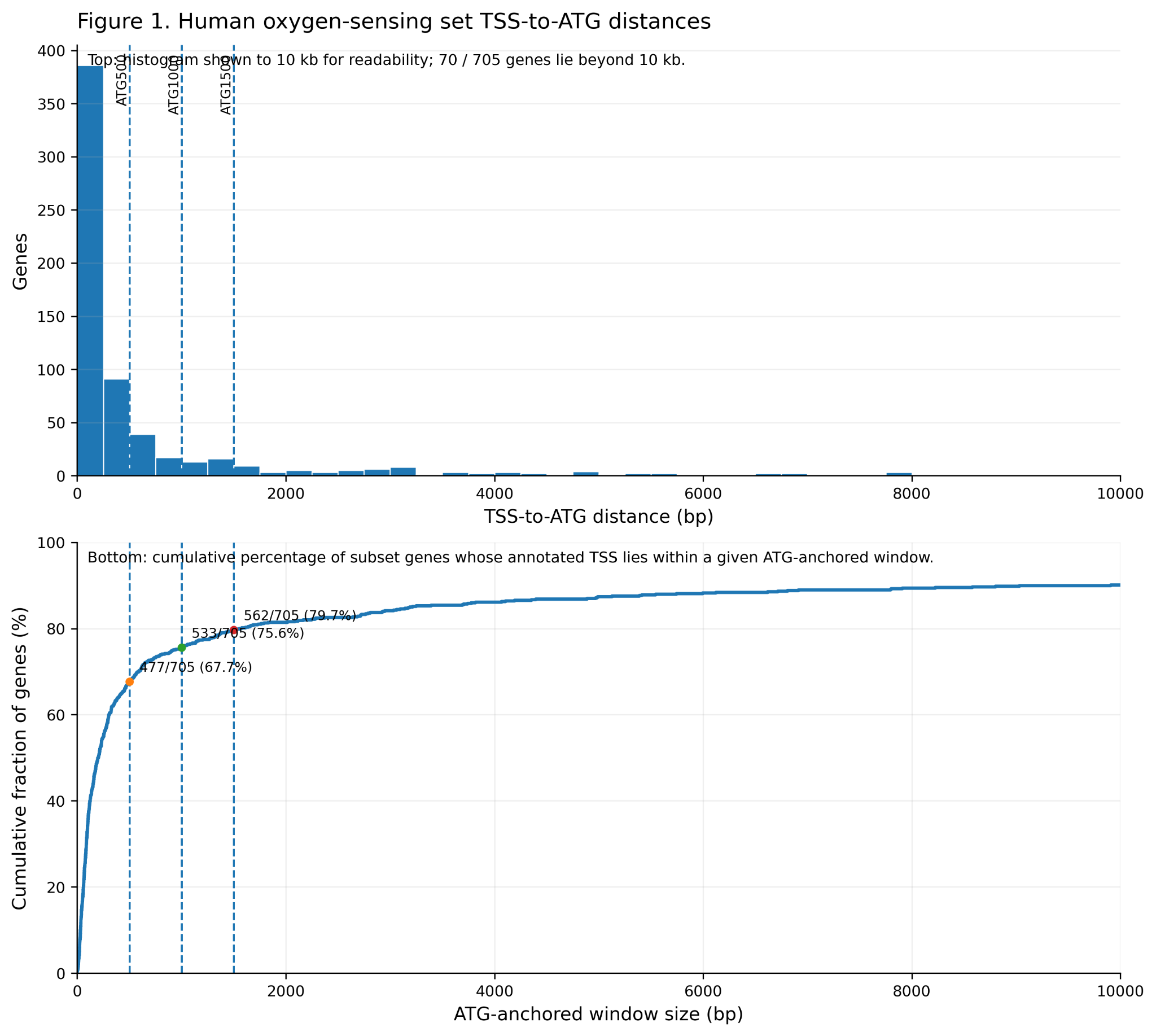

**Figure S2.** Human oxygen-sensing set TSS-to-ATG distances. Top: distribution of annotated human TSS-to-ATG genomic distances in the 705 representative genes in OGs containing oxygen sensing genes. Vertical dashed lines mark ATG500, ATG1000, and ATG1500. Bottom: cumulative fraction of genes whose annotated TSS lies within a given ATG-anchored window. In this subset, 477/705 genes (67.7%) lie within 500 bp, 533/705 (75.6%) within 1000 bp, and 562/705 (79.7%) within 1500 bp.

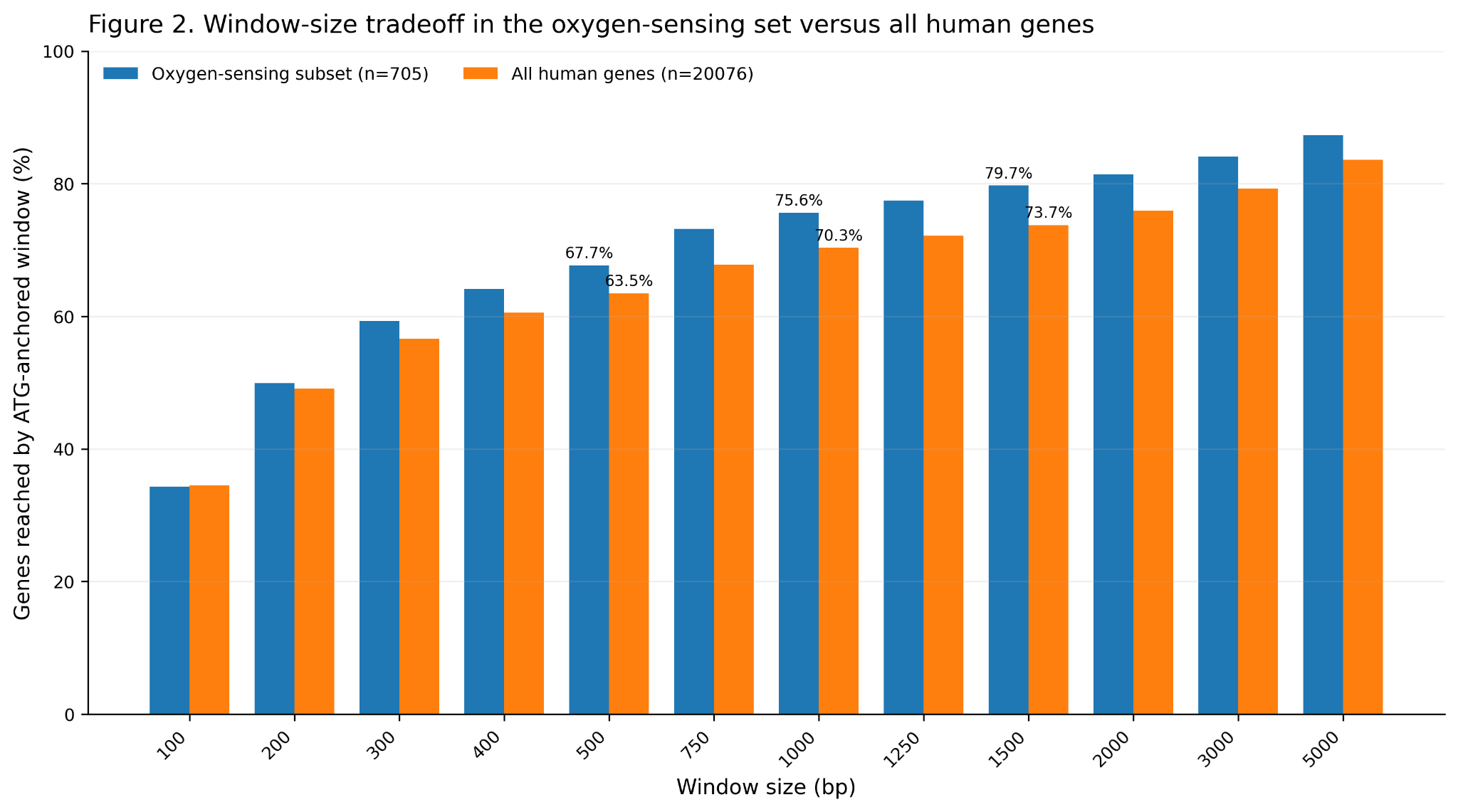

**Figure S3**. Window-size tradeoff in the oxygen-sensing set versus all human genes. Bars show the fraction of genes whose annotated TSS is reached by ATG-anchored windows of different sizes for the oxygen-sensing subset (n = 705) and for all human genes (n = 20,076). This is the direct estimate of the cost of the ATG choice in the best-annotated reference species.

##### Justification for 500 bp as the Window Width

The choice of 500 bp is motivated by the intersection of three considerations. First, the proximal promoter, the region with the highest density of functional TF binding and the strongest ChIP-seq signal, is concentrated within ~500 bp of the TSS in mammals. Second, as shown above, 500 bp upstream of the ATG captures the annotated TSS for the majority (67.7%) of the oxygen-sensing gene set in human. Third, the sensitivity analysis (Part 2 below) demonstrates that widening the window to 1,000 or 1,500 bp progressively inflates inter-species distances by adding lineage-specific distal TFBS that contribute noise rather than shared regulatory signal, without improving the recovery of biologically coherent structure. The 500 bp window therefore maximises signal-to-noise within the constraints of the cross-species annotation landscape.

##### Validity of the ReMap Evidence Filter Under ATG Anchoring

TFBS–gene pairs predicted by motif scanning were filtered against experimental evidence from ReMap 2022^6^, an atlas of over 165 million experimentally validated TF-binding regions from >600 human cell and tissue types. An edge is retained when at least one ReMap MACS peak interval^7^ overlaps the ATG 500 bp window. This is not a strict claim that the ChIP-seq summit lies within the ATG 500 bp window; it is evidence that a TF binds in the promoter-proximal neighbourhood of the gene in human. The median overlap between the ATG 500 bp window and retained TF MACS peak intervals is 367 bp, confirming that the majority of retained associations reflect substantive, not trivial, overlap. The ReMap filter thus operates as a conservative false-positive screen that remains valid under ATG anchoring.

##### Scope of the window: proximal-promoter architecture and distal enhancers

An ATG-anchored window samples a consistently defined gene-proximal region that typically overlaps promotor-proximal sequence but does not capture distal enhancers. We emphasize that this is a deliberate scoping decision rather than a claim that enhancers are unimportant: enhancers are major contributors to mammalian gene regulation and, like promoter-proximal sites, undergo rapid evolutionary turnover. They are, however, intractable to include in a comparative gene-TF matrix of this breadth for two reasons. First, enhancers are distributed diffusely across the genome, frequently residing tens to hundreds of kilobases from the genes they regulate, on either side of the gene and in either orientation, and often embedded within the introns of the target gene or of unrelated neighboring genes. Second, and consequently, an enhancer cannot be assigned to its target gene from sequence position alone; confident enhancer–gene assignment requires locus- and cell-type-specific chromosome-conformation evidence (3C, 4C, Hi-C, or comparable assays), which does not exist for the overwhelming majority of the 239 species in this dataset and cannot be inferred reliably across such deep divergence. Attributing distal binding sites to genes without such data would introduce assignment errors that scale with genomic distance and would vary unpredictably with assembly contiguity across species. We therefore restrict the analysis to the proximal-promoter region, where the sampled sequence can be associated consistently with the focal gene comparably across all species, and interpret our results as describing the evolution of proximal-promoter regulatory architecture. The turnover and distal-element considerations that motivate this scope are the same ones that make site-level and enhancer-based comparative approaches difficult to extend across hundreds of genomes (main text).

#### Part 2: Sensitivity analysis. Robustness to Window size and anchor position

To assess whether the biological conclusions depend on the choice of scanning window, the ATG 500 bp primary solution was compared against three alternatives:

- **ATG 1,000 bp**: tests sensitivity to window width (doubling)
- **ATG 1,500 bp**: tests sensitivity to window width (tripling)
- **TSS 500 bp**: tests sensitivity to anchor position (ATG vs. TSS, same width)

Four complementary statistical tests were applied, each targeting a distinct level of the analysis hierarchy. Prior to all tests, the PCA solutions for the alternative windows were aligned to the ATG 500 bp reference solution through axis matching, using the Hungarian algorithm (via clue::solve_LSAP) to find the globally optimal one-to-one mapping between axes that maximizes Tucker congruence, followed by sign correction of each matched axis. This step is necessary because PCA axes are defined only up to sign and permutation; without it, positional comparisons across solutions would confound genuine biological differences with artefactual axis-slot mismatches.

##### Test 1: Procrustes Analysis of PC Score Matrices

###### Method

Procrustes analysis (implemented via vegan::protest, symmetric = TRUE) superimposes two multi-dimensional species configurations by finding the optimal combination of rotation, reflection, and isotropic scaling that minimizes the sum of squared species displacements. The residual sum of squares after optimal superimposition is reported as m², which ranges from 0 (perfect congruence) to 1 (complete incongruence). The Procrustes correlation r = √(1 − m²) quantifies the degree of match as an effect size. Statistical significance was assessed by permuting species labels 9,999 times to generate a null distribution. Per-species displacement magnitudes (Procrustes residuals) were computed as the row-wise Euclidean distance between the two superimposed configurations after optimal rotation.

This test addresses sensitivity at the most global level: it asks whether the overall geometric arrangement of the 239 species in the nine-dimensional PC space, that is, who is architecturally similar to whom and by how much, is preserved when the window or anchor is changed.

###### Results

All three comparisons yielded highly significant Procrustes correlations (p < 0.001 in all cases; Table S1; Figure S4). The ATG 500 vs. ATG 1000 bp comparison produced the highest congruence (r = 0.9943, m² = 0.0113), indicating near-perfect preservation of the species configuration when the window is widened to 1,000 bp. Widening the window further to 1,500 bp reduced congruence modestly (r = 0.9835, m² = 0.0328). The ATG 500 bp vs. TSS 500 bp comparison yielded r = 0.9858 (m² = 0.0282), a level of congruence intermediate between the two ATG-window comparisons.

The per-species Procrustes residuals (Figure S5, panel B) were right-skewed in all three comparisons, with the large majority of species showing very small displacements. Median residuals were smallest for ATG 500 vs. ATG 1000 bp (median = 0.0050), followed by ATG 500 vs. ATG 1500 bp (median = 0.0072) and ATG 500 bp vs. TSS 500 bp (median = 0.0080). The right-skewed distributions indicate that a small subset of species shows greater displacement across windows than the majority, but no comparison produced bimodal residual distributions, which would be indicative of a systematic clade-level reorganization.

| **Comparison** | **Procrustes *r*** | ***m*²** | **Median residual** |
| --- | --- | --- | --- |
| ATG 500 vs. ATG 1,000 bp | 0.9943 | 0.0113 | 0.0050 |
| ATG 500 vs. ATG 1,500 bp | 0.9835 | 0.0328 | 0.0072 |
| ATG 500 vs. TSS 500 bp | 0.9858 | 0.0282 | 0.0080 |

Table S1: Procrustes correlations and m^2^ values for sensitivity comparisons.

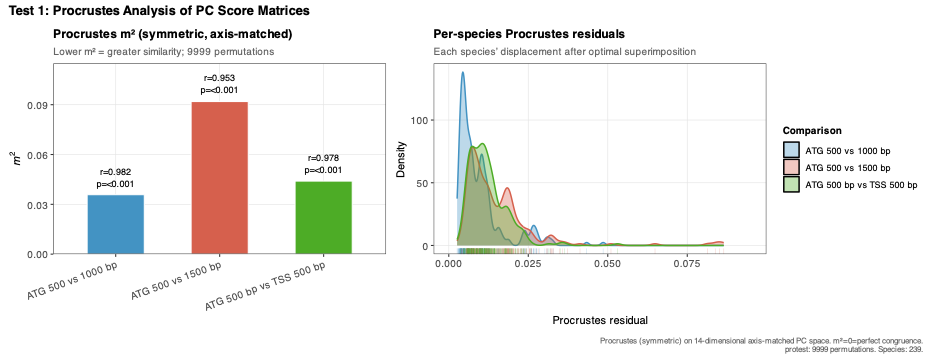

**Figure S4.** Test 1: Procrustes analysis of PC score matrices across window sizes and anchor positions. (A) Symmetric Procrustes m² statistic for three pairwise comparisons between the ATG 500 bp reference solution and the ATG 1000 bp, ATG 1500 bp, and TSS 500 bp alternatives. All axes were content-matched via the Hungarian algorithm prior to superimposition. Lower m² indicates greater similarity between configurations; annotated values show the Procrustes correlation r and permutation p-value (9,999 permutations). (B) Kernel density distributions of per-species Procrustes residuals for each comparison. Each residual represents the Euclidean displacement of one species in the nine-dimensional PC space after optimal rotation, reflection, and scaling. Distributions are right-skewed, indicating that the large majority of species are closely matched across windows, with a small number of species, corresponding to phylogenetically isolated taxa at the extremes of the regulatory architecture space, showing greater displacement. Rug marks show individual species residuals. Median residuals: ATG 500 vs. ATG 1000 bp = 0.0050; ATG 500 vs. ATG 1500 bp = 0.0072; ATG 500 vs. TSS 500 bp = 0.0080.

###### Interpretation

The results demonstrate that the nine-dimensional regulatory architecture space of 239 mammals is highly conserved across all four window conditions. The modest decline in congruence from ATG 1000 bp to ATG 1500 bp, despite the larger absolute window difference, is consistent with a pattern in which the 500–1,000 bp region adds a small amount of real regulatory signal (slightly widening inter-species distances in a consistent way, hence high r), while the 1,000–1,500 bp region adds progressively more lineage-specific distal TFBS that are weakly shared across species, introducing minor noise that reduces global congruence. The fact that the TSS 500 bp comparison achieves a congruence value (r = 0.9858) that falls between the two ATG window comparisons, rather than being lower than both, demonstrates that switching anchor from ATG to TSS does not systematically distort the species configuration. The slightly higher median residual for the TSS comparison reflects the expected consequence of anchor displacement in genes where the TSS is appreciably upstream of the ATG, but this effect is small and distributed smoothly across species rather than concentrated in specific clades. Together, the Procrustes results establish that the primary biological conclusions derived from the ATG 500 bp solution are highly robust to both window width and anchor choice.

##### Test 2: Mantel Test — Pairwise Species Distance Matrices

###### Method

Euclidean pairwise distance matrices were computed for each window in the nine-dimensional PC space. The Mantel test assessed the Pearson correlation between the lower triangles of the ATG 500 bp distance matrix and each comparison-window distance matrix, with significance evaluated by 9,999 permutations of species labels. Because Euclidean distance is rotation-invariant, this test is unaffected by axis permutation or sign flips. To complement the Mantel correlation, which captures rank fidelity, ordinary least-squares (OLS) regression was applied to each pairwise comparison (comparison distance ~ ATG 500 bp distance), providing slope and intercept estimates that characterize the metric (scale) relationship between windows. A slope of 1 and intercept of 0 indicates that the two windows are metrically equivalent; deviations from this benchmark reveal systematic inflation or compression of inter-species distances.

###### Results

All comparisons yielded highly significant Mantel correlations (p < 0.001; Figure S5). ATG 500 vs. ATG 1000 bp produced the highest rank correspondence (r = 0.9893), followed by ATG 500 bp vs. TSS 500 bp (r = 0.9808) and ATG 500 vs. ATG 1500 bp (r = 0.9731).

The scatter plots in Figure S5 reveal the metric relationships in detail:

ATG 500 vs. ATG 1000 bp (r = 0.9893, slope = 1.0887, intercept = −2.7905): pairwise species distances are closely aligned along the regression line, with a slope slightly above 1, indicating modest systematic inflation of distances by the wider window. A marked deviation from the regression line occurs in the upper right region of the plot, where species pairs with high ATG 500 bp distances (just below 200) are assigned appreciably larger distances under ATG 1000 bp (above 200). Two distinct clusters are visible in this upper-right region: one at approximately ATG 500 bp = 190, ATG 1000 bp = 220, and another at approximately ATG 500 bp = 190, ATG 1000 bp = 240. These clusters correspond to the most phylogenetically divergent cross-clade species pairs (e.g., monotremes paired with eutherians), whose already-large architectural distances become proportionally further inflated by the addition of lineage-specific distal TFBS in the wider window.

ATG 500 vs. ATG 1500 bp (r = 0.9731, slope = 1.1246, intercept = −2.9513): a similar pattern is evident, but more pronounced. The slope is higher than in the ATG 1000 bp comparison, confirming progressive distance inflation with window width. The upper-right deviation is also more pronounced, with two clusters visible at approximately ATG 500 bp = 190, ATG 1500 bp = 230 and ATG 500 bp = 190, ATG 1500 bp = 280. The increasing displacement of the upper-right clusters across ATG 1000 and ATG 1500, from ~220 to ~240, and from ~230 to ~280, is consistent with progressive accumulation of lineage-specific TFBS as window width increases, which disproportionately affects already-distant cross-clade pairs.

ATG 500 bp vs. TSS 500 bp (r = 0.9808, slope = 0.8775, intercept = 1.5200): the pattern here is qualitatively distinct from the ATG-window comparisons. The slope is below 1, indicating that inter-species distances are compressed under TSS anchoring relative to ATG anchoring. A single cluster is visible in the upper-right region, at approximately ATG 500 bp = 190, TSS 500 bp = 170, rather than the two clusters seen in the ATG-window comparisons. Importantly, these high-distance pairs fall above the OLS regression line but below the 1:1 identity line, meaning their TSS 500 bp distances, while compressed relative to ATG 500 bp, are still disproportionately large relative to the regression prediction. The positive intercept (1.52) combined with slope < 1 reflects a characteristic compression-with-offset geometry: close pairs show slightly larger TSS distances than expected (intercept effect), while distant pairs show smaller TSS distances than expected (slope effect).

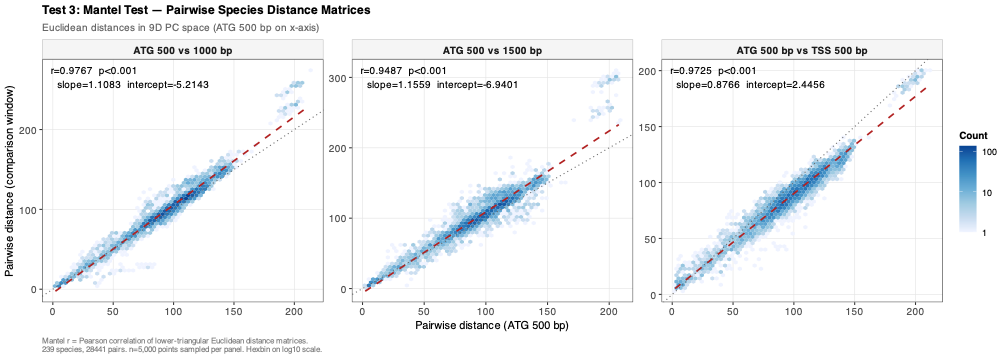

**Figure S5.** Test 2: Mantel test of pairwise species distance matrices across window sizes and anchor positions. Each panel shows a hexbin scatter plot of pairwise Euclidean species distances in nine-dimensional PC space, with ATG 500 bp distances on the x-axis and comparison-window distances on the y-axis (5,000 randomly sampled pairs per panel; hexbin colour on log₁₀ scale). The dashed red line shows the OLS regression fit (comparison ~ ATG 500 bp); the dotted grey line is the 1:1 identity line (slope = 1, intercept = 0). Deviation of the regression line from the identity line quantifies systematic distance inflation (slope > 1; ATG 1000 and ATG 1500 bp panels) or compression (slope < 1; TSS 500 bp panel) relative to the reference window. Annotated values give the Mantel r, OLS slope, and OLS intercept. The upper-right deviation from the regression line in all panels reflects the disproportionate displacement of the most phylogenetically distant species pairs (e.g., monotremes vs. eutherians) across window conditions; the two-cluster structure visible in the ATG 1000 bp and ATG 1500 bp panels corresponds to two groups of highly divergent cross-clade pairs whose distances are differentially amplified by the addition of lineage-specific distal TFBS.

###### Interpretation

The Mantel results confirm that the rank order of inter-species regulatory architecture distances is highly preserved across all four window conditions. The slope analysis adds a critical complementary layer: the ATG 500 bp window is the most parsimonious choice in terms of distance scaling. Widening the ATG window progressively inflates inter-species distances, disproportionately so for already-distant cross-clade pairs, due to the accumulation of lineage-specific distal TFBS that do not represent shared regulatory signal. The TSS 500 bp anchor produces a slope below 1, reflecting a modest compression of inter-species distances (the opposite effect) which arises because TSS-proximal TFBS counts are somewhat lower on average and vary less across species than ATG-proximal counts in this gene set. Crucially, the pattern of which species pairs are most architecturally distant is preserved across all four windows (Mantel r > 0.97 in all cases), confirming that the biological groupings and gradients identified in the primary analysis are not an artefact of the 500 bp ATG window choice.

##### Test 3: Variance Explained per PC Axis

###### Method

For each window, the percentage of total variance explained by each PC axis was computed as the column variance of the species scores divided by the sum of column variances across all nine axes. Cumulative variance profiles were also derived. This test does not assess species- or gene-level congruence but instead examines whether the window choice alters the dimensionality structure of regulatory architecture variation, specifically, whether the same proportion of total variation is captured by the same number of axes regardless of window.

###### Results

The variance profiles across all four windows were closely parallel (Figure S6). PC1 consistently captured the largest share of variance across all windows (ATG 500 bp: 19.6%; ATG 1000 bp: 19.1%; ATG 1500 bp: 19.5%; TSS 500 bp: 19.1%), followed by PC2 (ATG 500 bp: 16.2%; ATG 1000 bp: 16.7%; ATG 1500 bp: 16.7%; TSS 500 bp: 15.3%). The cumulative variance curves were nearly superimposable across windows, with all four solutions reaching 50% cumulative variance at PC3 and 90% cumulative variance at PC7–PC8.

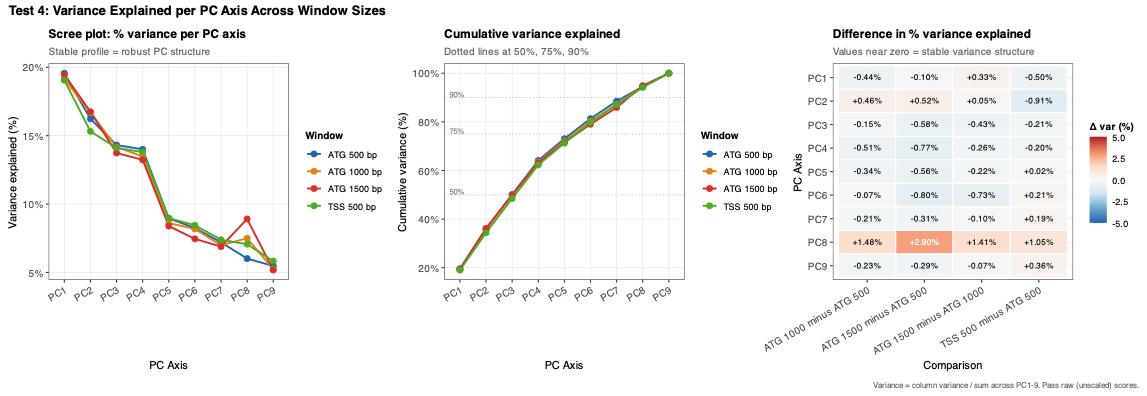

**Figure S6.** Test 3: Variance explained per PC axis (first nine axes considered) across window sizes and anchor positions. (A) Scree plot showing the percentage of total variance (sum of column variances across PC1–PC9) explained by each axis, separately for each window. (B) Cumulative variance profiles; dotted horizontal lines mark the 50%, 75%, and 90% thresholds. (C) Delta heatmap showing the difference in percentage variance between each comparison window and the ATG 500 bp reference (positive values = comparison window explains more variance on that axis; negative values = less). Near-zero values across all axes indicate that the variance structure is highly stable across window and anchor conditions.

###### Interpretation

The near-identical scree profiles across all four windows indicate that the dimensionality of mammalian regulatory architecture space is a robust property of the data, not a feature introduced by the specific window or anchor used. The consistently even spread of variance across nine axes, with no single axis dominating, is preserved regardless of whether the window is widened by up to 1,000 bp or the anchor is shifted from ATG to TSS. The only noteworthy deviation is the slight reduction in PC2 variance under TSS 500 bp. PC2 captures the deepest phylogenetic split in the dataset (monotremes and marsupials versus placentals); the modest reduction in its variance share under TSS anchoring suggests that this dimension of regulatory architecture variation is marginally less accentuated in the TSS-proximal window, consistent with the general distance compression seen in Test 2. Nonetheless, PC2 retains the second-largest variance share under all four windows, and its biological interpretation is unchanged. The variance partitioning results thus confirm that the choice of window size and anchor does not alter the fundamental structure of how regulatory architecture variation is distributed across the nine principal axes of mammalian diversity.

##### Test 4: Blomberg's K of PC Scores

###### Method

Blomberg's K statistic measures the degree of phylogenetic signal in a continuous trait relative to the expectation under Brownian motion (Blomberg et al., 2003, Evolution 57:717–745). A value of K = 1 indicates that close relatives are precisely as similar as expected under Brownian motion; K > 1 indicates stronger-than-expected phylogenetic conservatism; K < 1 indicates greater evolutionary lability than expected. K was computed independently for each of the nine PC axes and for each window using phytools::phylosig (method = "K") with 1,000 permutations for significance testing. The trait in each case was the species score on a given PC axis, that is, the species' position in regulatory architecture space along that dimension. This test addresses whether the phylogenetic signal embedded in the regulatory architecture is sensitive to window or anchor choice.

###### Results

Phylogenetic signal was statistically significant (p < 0.001) for all nine PC axes under all four window conditions without exception (Figure S7). Blomberg's K values were substantially greater than 1 across all axes and all windows, indicating strong phylogenetic conservatism of regulatory architecture throughout the regulatory architecture space. The magnitude and rank ordering of K values across axes was largely preserved across windows, with PC2 consistently displaying the highest K by a wide margin (ATG 500 bp: K = 41.09; ATG 1000 bp: K = 53.66; ATG 1500 bp: K = 59.74; TSS 500 bp: K = 30.72), followed by PC8 (ATG 500 bp: K = 11.74; ATG 1500 bp: K = 12.39) and PC4 (ATG 500 bp: K = 8.92).

All K values are significant at p < 0.001 (1,000 permutations).

Two patterns merit particular attention. First, the TSS 500 bp window consistently yields lower K values across all axes, most notably on PC1 (ATG 500 bp: K = 6.17 vs. TSS 500 bp: K = 1.67), PC3 (7.24 vs. 4.10), and PC4 (8.92 vs. 3.09). Second, PC6 shows marked instability across ATG windows (K = 1.15 at ATG 500 bp, 4.41 at ATG 1000 bp, 1.60 at ATG 1500 bp), though it remains highly significant under all conditions.

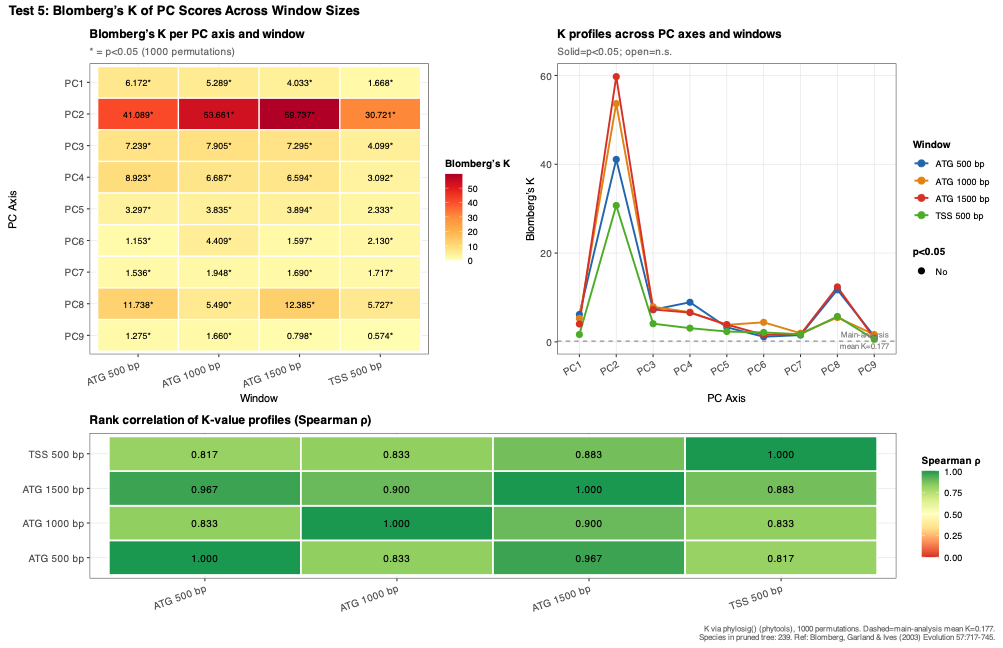

**Figure S7.** Test 4: Blomberg's K of PC scores across window sizes and anchor positions. (A) Heatmap of Blomberg's K per PC axis and window. Asterisks indicate p < 0.05 (1,000 permutations; all axes are significant at p < 0.001 across all windows). (B) Line profiles of K across PC axes for each window; filled symbols indicate p < 0.05, open symbols indicate non-significant values. The dashed horizontal line marks the mean K from the primary ATG 500 bp analysis for reference. (C) Spearman rank correlation matrix of K-value profiles (PC1–PC9) between all pairs of windows, quantifying the degree to which the phylogenetic signal hierarchy across axes is conserved irrespective of window or anchor choice.

###### Interpretation

The universal significance of phylogenetic signal across all nine axes under all four window conditions is the central finding of Test 4 and confirms the robustness of one of the study's core claims: that regulatory architecture variation across mammals is not random with respect to phylogeny but reflects evolutionarily conserved patterns of TF assignment. This result is not trivially expected; if window or anchor choice were introducing substantial noise, phylogenetic signal would be expected to decay on lower-variance axes (PC6–PC9), which carry smaller and more diffuse patterns of inter-species variation. The preservation of significant and supra-Brownian K on these axes across all windows is therefore strong evidence against noise accumulation as a confound.

#### Overall Conclusion: Sensitivity of Results to Window size and Anchor Choice

Across four independent and methodologically complementary tests, the regulatory architecture of the mammalian oxygen-sensing network, as captured by the nine-dimensional TFBS-PCA solution, is highly robust to variation in both scanning window width and anchor position. The overall geometric configuration of 239 species in PC space was preserved to a Procrustes correlation of r ≥ 0.984 under all three alternative window conditions (Test 1), with per-species displacements that were small, continuously distributed, and free of any clade-level discontinuity. The rank ordering of inter-species regulatory architecture distances was maintained with Mantel r ≥ 0.973 across all comparisons (Test 2), and the dimensionality structure of regulatory variation, the proportion of total variance distributed across nine axes, was stable to within ~1–2 percentage points per axis across all windows (Test 3). Most critically, phylogenetic signal was significant at p < 0.001 on all nine PC axes under all four window conditions, with Blomberg's K consistently and substantially exceeding 1 (Test 4), confirming that the core finding of the main text, that mammalian regulatory architecture variation is deeply structured by phylogenetic history, is not a product of the specific window or anchor used.

The two sources of variation introduced by wider windows and TSS anchoring are well-characterized and mechanistically interpretable. Widening the ATG window to 1,000 or 1,500 bp progressively inflates inter-species distances (slopes of 1.09 and 1.12 respectively; Test 2) due to the accumulation of lineage-specific distal TFBS that contribute noise rather than shared regulatory signal; this inflation is proportional and disproportionately affects already-distant cross-clade pairs, leaving within-clade structure intact. TSS anchoring produces the opposite metric effect, a slope of 0.88 relative to ATG 500 bp (Test 2) and modestly lower Blomberg's K values across axes (Test 4), consistent with the known lower average density of TFBS in the TSS-proximal window for this gene set, which compresses inter-species distances without reorganizing the species configuration. Neither effect alters the identity of the principal axes, the groupings they define, the ecological and phylogenetic gradients they capture, or the significance of any biological claim made in the main text. The ATG 500 bp window thus represents the most parsimonious choice: it maximizes the signal-to-noise ratio by focusing on the functionally validated proximal promoter region while avoiding the distance inflation introduced by wider windows, and it produces a species configuration that is metrically and structurally equivalent to the TSS-anchored solution while being more reliably annotated across the full 239-species comparative dataset. The species configuration equivalence of the PCA space across windows and anchors is exemplified in Figure S8a-c.

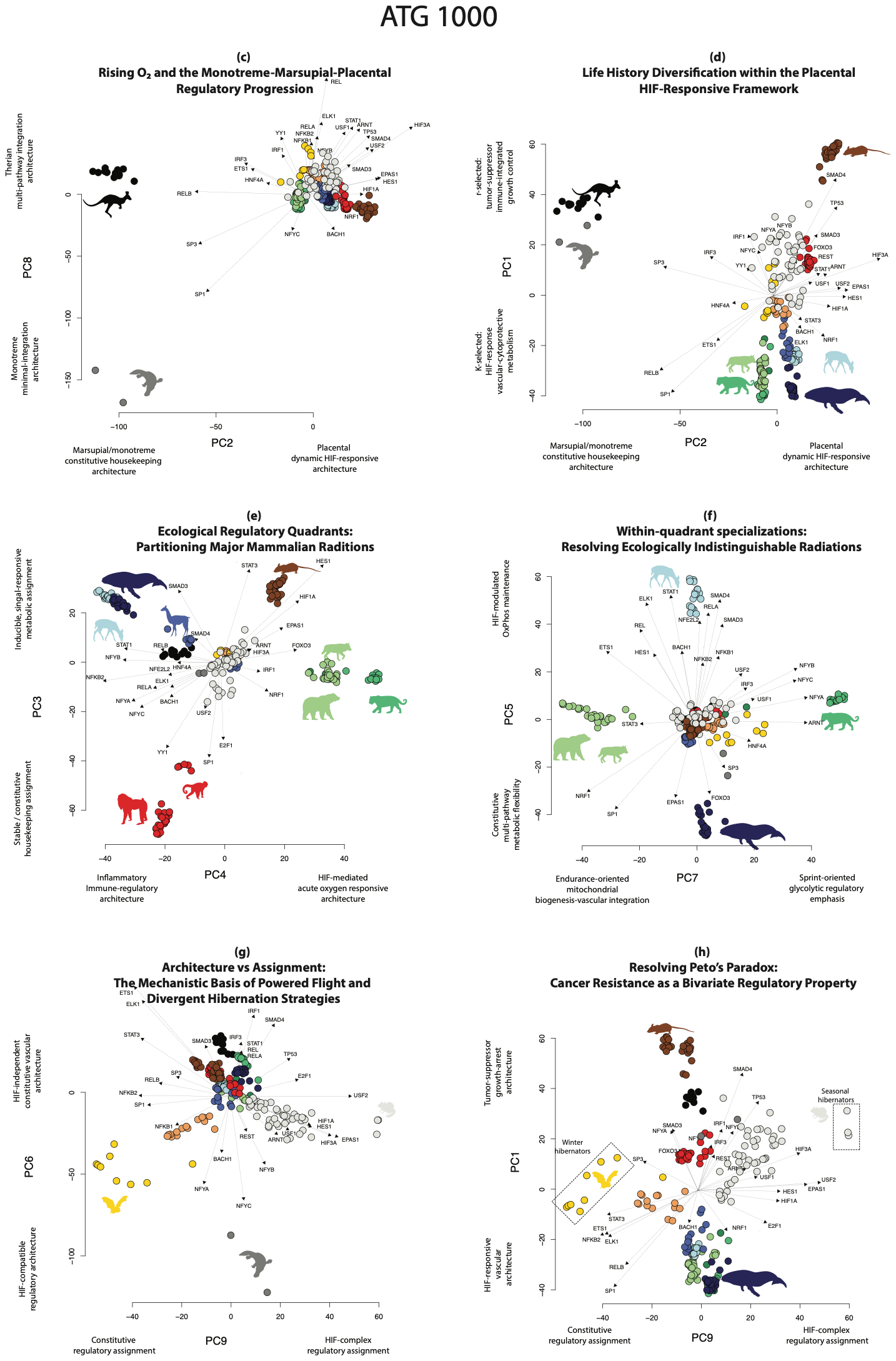

**Figure S8a.** As in Figure 1c-h, but for ATG 1000.

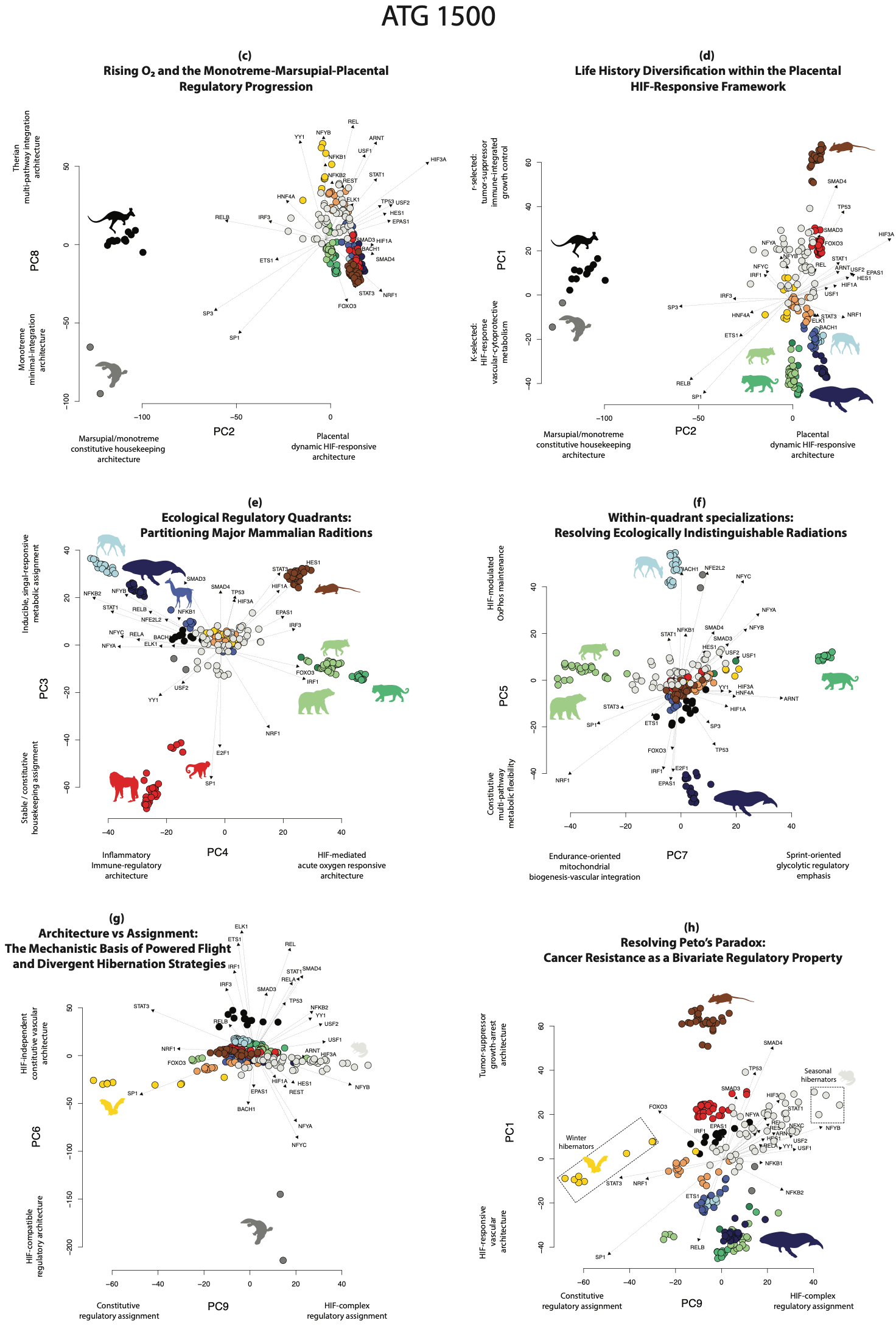

**Figure S8b.** As in Figure 1c-h, but for ATG 1500.

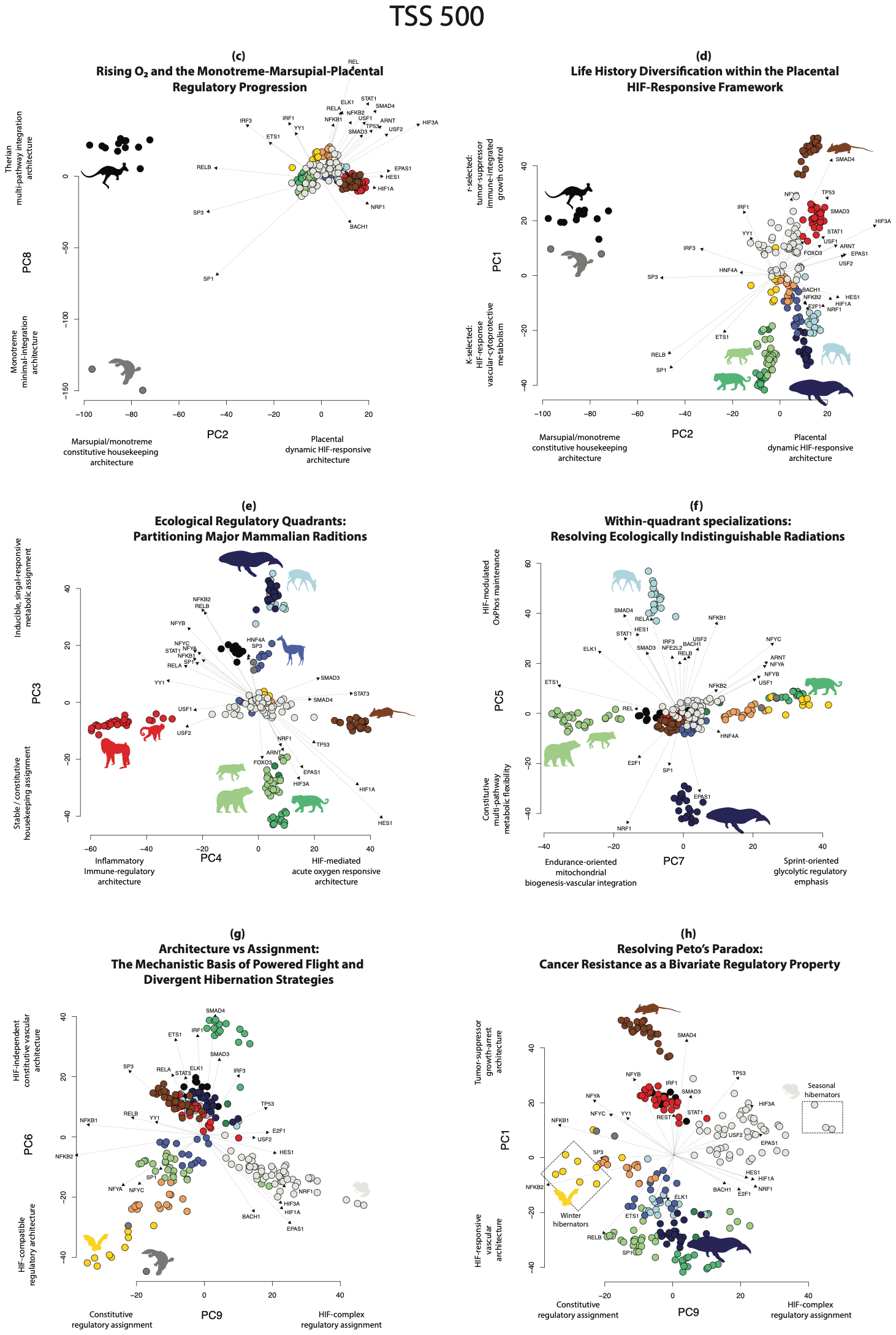

**Figure S8c.** As in Figure 1c-h, but for TSS 500.

### Sensitivity of the regulatory-architecture PCA to genome-annotation method: OrthoFinder/NCBI versus TOGA

#### Overview

The primary analyses in this study derive transcription factor binding site (TFBS) architecture from NCBI gene models, with orthogroups inferred using OrthoFinder on NCBI genome assemblies. To assess whether the biological conclusions depend on this annotation pipeline, we compared the primary solution against an independent dataset in which the same oxygen-sensing gene–TF architecture was quantified from TOGA (Tool to infer Orthologs from Genome Alignments) annotations^8^. Because TOGA and OrthoFinder/NCBI differ in the set of species recovered, the set of gene–TF features recovered, and the quantitative binding-site values assigned to shared features, a valid comparison must control the first two so that any residual incongruence is attributable to the annotation method itself. We therefore restricted both datasets to the species and gene–TF features they share and re-computed the PCA on that common basis. Both datasets use the 500 bp window anchored at the transcription start site (TSS 500 bp); the primary comparison holds this anchor constant, and a secondary comparison additionally relates the TOGA solution to the study’s published main analysis (ATG 500 bp).

#### Part 1: Scope of the comparison

The OrthoFinder/NCBI dataset comprises 239 species and 12,918 gene–TF pairs (main text analysis); the TOGA dataset comprises 504 species and 13,876 gene–TF pairs. The two share 193 species and 11,123 gene–TF pairs. Critically, TOGA contains no marsupials or monotremes: all 14 such species present in the NCBI dataset are absent, so the 193 shared species are all placental (of the 46 NCBI species not shared, 14 are marsupials/monotremes and 32 are placentals absent from the TOGA assembly set).

This has a direct and unavoidable consequence for two axes. In the published analysis PC2 separates monotremes and marsupials from placentals (group means: monotremes −86, marsupials −84, placentals +5) and PC8 is dominated by monotremes (monotreme mean −142 versus ≈0 for all other taxa). These axes are defined by the very taxa that TOGA lacks. Quantitatively, the fraction of each published axis’s variance that survives into the placental subset (the ratio of its standard deviation among the 193 shared placentals to its standard deviation among all 239 species) is ≈1.0 for every axis except PC2 (0.52) and PC8 (0.31), which collapse (Table S2, Figure S9a). PC2 and PC8 therefore cannot be assessed against TOGA with the currently available taxon sampling, and are reported here as out of scope. The annotation-sensitivity analysis necessarily concerns the within-placental regulatory architecture captured by PC1, PC3–PC7, and PC9.

**Table S2.** Survival of each published PC axis into the placental (TOGA-shared) subset.

| Published axis | Variance (%, 239 sp) | Survival ratio | Testable vs TOGA | Biology |
| --- | --- | --- | --- | --- |
| PC1 | 6.3 | 0.98 | yes | r–K life history / body size / tumor-suppressor (cancer) |
| PC2 | 5.3 | **0.52** | **no (deep taxa)** | SP1→HIF; monotreme–marsupial–placental |
| PC3 | 4.6 | 1.04 | yes | constitutive vs inducible ecological control |
| PC4 | 4.5 | 1.03 | yes | NF-κB/IRF vs HIF ecological control |
| PC5 | 2.9 | 1.02 | yes | cetacean vs Pecora |
| PC6 | 2.7 | 0.99 | yes | HIF-independent vs HIF-compatible architecture (flight) |
| PC7 | 2.3 | 1.04 | yes | feliform vs caniform |
| PC8 | 2.0 | **0.31** | **no (deep taxa)** | multi-pathway integration; monotreme-dominated |
| PC9 | 1.8 | 1.04 | yes | HIF-complex assignment (hibernation; cancer) |

#### Part 2: Methods

Both raw species × gene–TF matrices were restricted to the 193 shared species and 11,123 shared gene–TF pairs, so that species set, feature set, phylogeny, and missing-value imputation were held identical and the only difference between the two matrices was the annotation-derived binding-site values. Each restricted matrix was then processed through the study’s standard pipeline independently: Brownian-motion phylogenetic imputation of missing values on the shared 193-tip ultrametric tree, followed by principal component analysis on the standardized features (prcomp, scale = TRUE). This re-fitting is essential: subsetting the scores of a PCA fitted on a different sample would project the shared species onto axes optimised for a different species cloud, confounding annotation differences with sampling artefacts. Re-fitting on the common basis avoids this.

Because principal axes are defined only up to sign and permutation, and because removing the deep taxa renumbers the placental solution relative to the published one, axes were matched across solutions over a wide range (up to 20 components) using the Hungarian algorithm on Tucker congruence coefficients, allowing an axis in one solution to correspond to a differently-numbered axis in the other. Each published axis was additionally bridged to its counterpart in the re-fitted solutions by maximal score congruence over the 193 shared placentals, so that every tested axis retains its published biological identity. Congruence was quantified with (i) symmetric Procrustes analysis of the species score configurations (vegan::protest, 9,999 permutations), (ii) the Mantel correlation of Euclidean species-distance matrices (9,999 permutations), (iii) per-axis Tucker congruence coefficients, and (iv) Blomberg’s K per axis (phytools::phylosig, 1,000 permutations). For reference, the study’s window/anchor sensitivity analysis established robustness at Procrustes r ≥ 0.984 and Mantel r ≥ 0.973.

#### Part 3: Results

##### Test 1: Global configuration congruence (Procrustes and Mantel).

The species configuration in the within-placental regulatory space is strongly congruent between annotation methods, and the leading dimensions match the window/anchor benchmark (Table S3, Figure S9b). For the primary annotation-isolated comparison (NCBI-TSS vs TOGA, same anchor), Procrustes r = 0.991 over the first six matched axes and remained at or above the 0.984 window/anchor benchmark through all nine matched axes (r = 0.984 at nine). The secondary comparison against the published main analysis (NCBI-ATG vs TOGA) behaved almost identically (r = 0.995 at six axes; 0.977 at nine). Crucially, switching annotation method perturbed the placental configuration by no more than switching anchor did within NCBI: the anchor-only reference (NCBI-ATG vs NCBI-TSS on the same 193 placentals) gave r = 0.991 to 0.968 over the same range; comparable to, and no more congruent than, the annotation comparisons. Mantel correlations followed the same pattern (at nine axes: 0.97, 0.98, and 0.96 for the primary, secondary, and anchor comparisons, respectively). All tests were significant at p < 0.001.

**Table S3.** Global Procrustes correlation (r) over matched axes, by number of axes (# axes).

| # axes | NCBI-TSS vs TOGA (annotation) | NCBI-ATG vs TOGA (vs headline) | NCBI-ATG vs NCBI-TSS (anchor ref.) |
| --- | --- | --- | --- |
| 6 | 0.991 | 0.995 | 0.991 |
| 7 | 0.984 | 0.992 | 0.984 |
| 8 | 0.986 | 0.984 | 0.981 |
| 9 | 0.984 | 0.977 | 0.968 |

##### Test 2: Per-axis reproduction of the published biology.

Each testable published axis has a clear TOGA counterpart, though at a shifted position because PC2 and PC8 drop out among placentals (Table S4, Figure S9c). The two axes carrying the strongest claims, PC1 (life history, body size and the tumor-suppressor pole of the cancer framework) and PC7 (the caniform–feliform divergence), are reproduced excellently (Tucker φ = 0.97 each). The within-quadrant axis PC5 (cetacean vs Pecora), the HIF-complex assignment axis PC9 (the second axis of the cancer framework), and the flight axis PC6 are reproduced strongly (φ = 0.93, 0.92, and 0.91), whereas the two ecological-control axes PC3 and PC4 are the weakest, reproduced only moderately (φ = 0.78 and 0.80). The renumbering is systematic (PC4→TOGA PC2, PC5→PC4, PC6→PC5, PC7→PC6, PC9→PC7), reflecting the excision of the two deep-taxon axes rather than any reorganisation of the biology.

**Table S4.** Reproduction of each testable published axis by TOGA.

| Published axis (biology) | TOGA counterpart | Tucker φ |
| --- | --- | --- |
| PC1 — life history / body size / cancer | TOGA PC1 | 0.97 |
| PC3 — constitutive vs inducible ecology | TOGA PC3 | 0.78 |
| PC4 — NF-κB vs HIF ecology | TOGA PC2 | 0.80 |
| PC5 — cetacean vs Pecora | TOGA PC4 | 0.93 |
| PC6 — flight architecture | TOGA PC5 | 0.91 |
| PC7 — feliform vs caniform | TOGA PC6 | 0.97 |
| PC9 — HIF assignment (hibernation; cancer) | TOGA PC7 | 0.92 |

##### Test 3: Phylogenetic signal (Blomberg’s K).

Phylogenetic signal is preserved on every testable axis and is, on most axes, stronger under TOGA (Table S5). Blomberg’s K exceeds 1 for all seven testable axes in all datasets, and is higher in TOGA than in NCBI for six of the seven testable axes (e.g. PC1: 10.1 in TOGA vs 7.7/2.7 in NCBI-ATG/TSS; PC3: 12.4 vs 11.1/4.1); the sole exception is PC7, where the NCBI-TSS value (5.0) marginally exceeds TOGA (4.7). This confirms that the core finding — that regulatory architecture variation is deeply structured by phylogeny — does not depend on the annotation pipeline.

**Table S5.** Blomberg’s K per testable axis (all p < 0.05).

| Published axis | NCBI-ATG | NCBI-TSS | TOGA |
| --- | --- | --- | --- |
| PC1 | 7.67 | 2.74 | 10.10 |
| PC3 | 11.08 | 4.12 | 12.44 |
| PC4 | 12.82 | 5.92 | 13.10 |
| PC5 | 4.67 | 3.75 | 4.95 |
| PC6 | 3.70 | 3.29 | 5.97 |
| PC7 | 3.50 | 4.98 | 4.70 |
| PC9 | 2.58 | 1.14 | 4.09 |

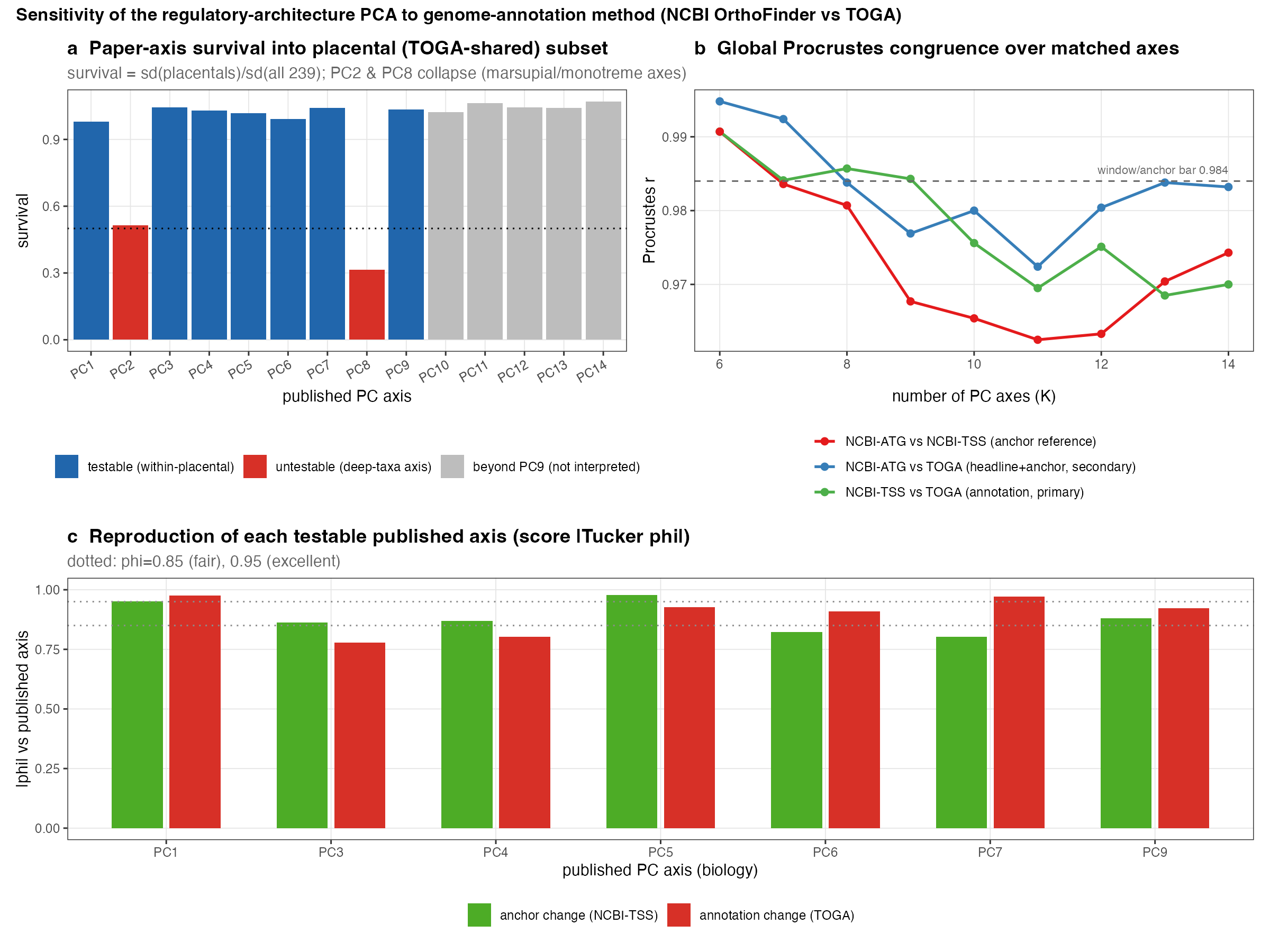

Figure S9. Robustness of the regulatory-architecture PCA to genome-annotation method (OrthoFinder/NCBI vs TOGA). (a) Survival of each published PC axis into the placental subset shared with TOGA; PC2 and PC8, defined by marsupials and monotremes, collapse and are untestable. (b) Global Procrustes congruence of the species configuration over matched axes, for the annotation comparison (green), the comparison against the published main analysis (blue), and the within-NCBI anchor reference (red); dashed line, the window/anchor robustness benchmark (r = 0.984). (c) Per-axis reproduction of each testable published axis (score Tucker φ) under anchor change (green) and annotation change (red); dotted lines mark φ = 0.85 (fair) and 0.95 (excellent).

#### Overall conclusion

Within its testable scope, the oxygen-sensing regulatory-architecture space is robust to genome-annotation method. The axes carrying the study’s principal claims (the life-history/body-size/cancer axis (PC1), the ecological-quadrant axes (PC3/PC4), the carnivoran (PC7) and cetacean/ruminant (PC5) specialization axes, and the HIF-assignment/hibernation axis (PC9)) are recovered by an independent TOGA annotation (the two ecological-quadrant axes, PC3/PC4, more moderately, φ ≈ 0.78–0.80), with the leading dimensions matching the same congruence benchmark established for window and anchor choice, and with phylogenetic signal preserved or strengthened. Switching annotation method perturbs the placental configuration no more than (and, over the nine matched axes, slightly less than) the already-validated change of anchor from ATG to TSS. Two axes cannot be evaluated with the current data: PC2 (the monotreme–marsupial–placental transition underlying the atmospheric-oxygen narrative) and PC8 (a monotreme-dominated axis) are defined by taxa that the TOGA dataset does not include, and their robustness to annotation would require a TOGA annotation that samples marsupials and monotremes. The species configuration equivalence of the PCA space across annotation pipeline is exemplified in Figure S10a-b.

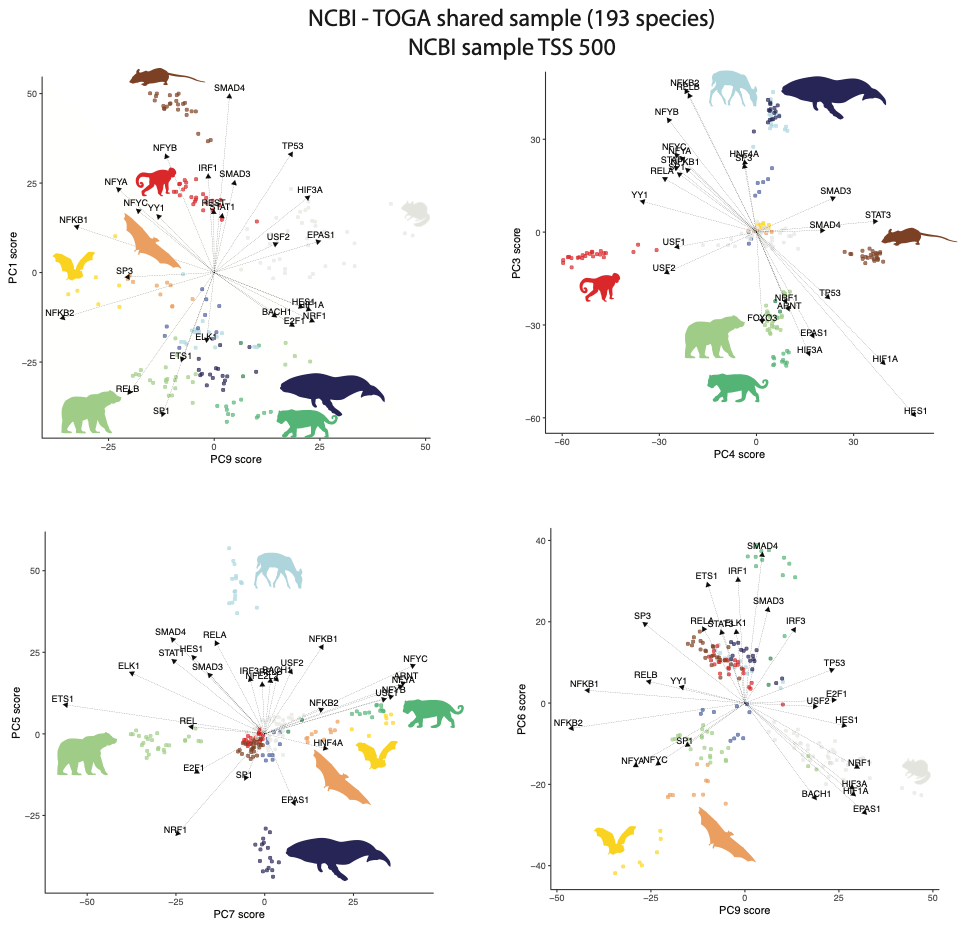

**Figure S10a.** The PCA spaces of the comparable axes between the NCBI-Orthofinder pipeline and the TOGA annotation pipeline for the NCBI-Orthofinder sample.

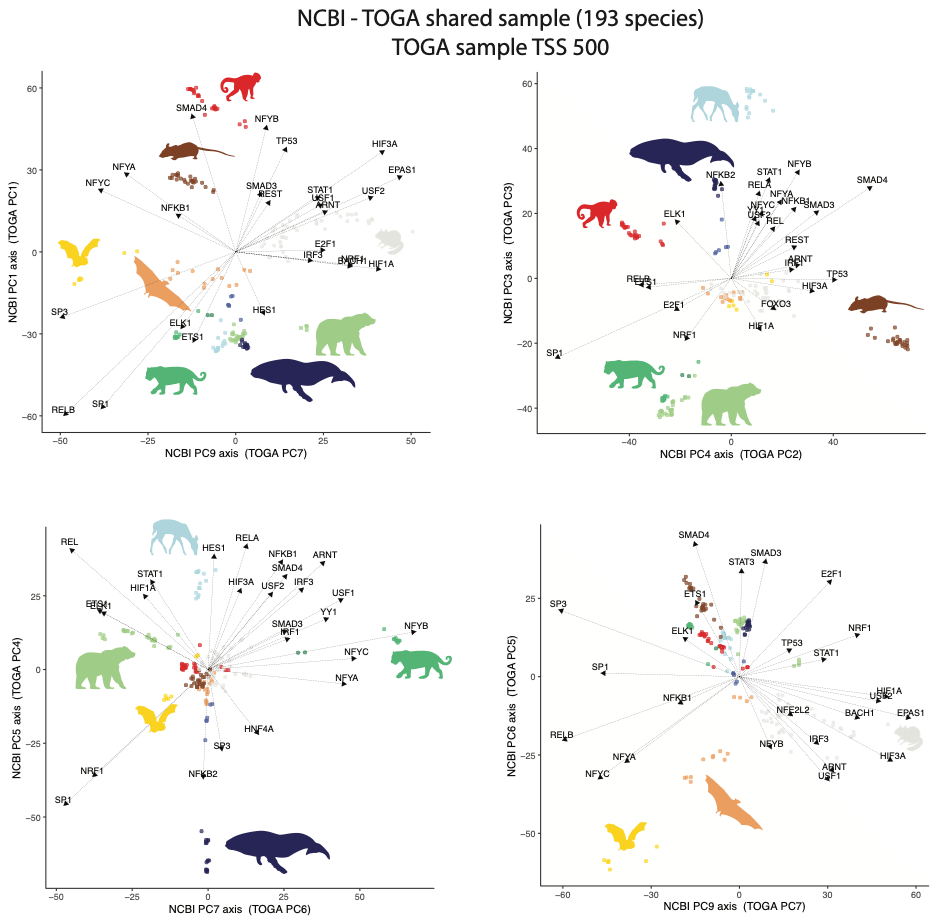
 **Figure S10b.** The PCA spaces of the comparable axes between the NCBI-Orthofinder pipeline and the TOGA annotation pipeline for the TOGA sample.

1	Haberle, V. & Stark, A. Eukaryotic core promoters and the functional basis of transcription initiation. *Nature reviews Molecular cell biology* **19**, 621-637 (2018).

2	Mudge, J. M. *et al.* GENCODE 2025: reference gene annotation for human and mouse. *Nucleic acids research* **53**, D966-D975 (2025).

3	Morales, J. *et al.* A joint NCBI and EMBL-EBI transcript set for clinical genomics and research. *Nature* **604**, 310-315 (2022).

4	Sample, P. J. *et al.* Human 5′ UTR design and variant effect prediction from a massively parallel translation assay. *Nature biotechnology* **37**, 803-809 (2019).

5	Park, S. G., Hannenhalli, S. & Choi, S. S. Conservation in first introns is positively associated with the number of exons within genes and the presence of regulatory epigenetic signals. *BMC genomics* **15**, 526 (2014).

6	Hammal, F., De Langen, P., Bergon, A., Lopez, F. & Ballester, B. ReMap 2022: a database of Human, Mouse, Drosophila and Arabidopsis regulatory regions from an integrative analysis of DNA-binding sequencing experiments. *Nucleic acids research* **50**, D316-D325 (2022).

7	Feng, J., Liu, T., Qin, B., Zhang, Y. & Liu, X. S. Identifying ChIP-seq enrichment using MACS. *Nature protocols* **7**, 1728-1740 (2012).

8	Malovichko, Y. V. *et al.* Accurate, comprehensive gene annotation and ortholog identification across thousands of vertebrate genomes with TOGA2. *bioRxiv* (2026).
