## Supplementary figures and images for "Oxygen-sensing regulatory architecture structures mammalian diversification"

### Fig. ED1

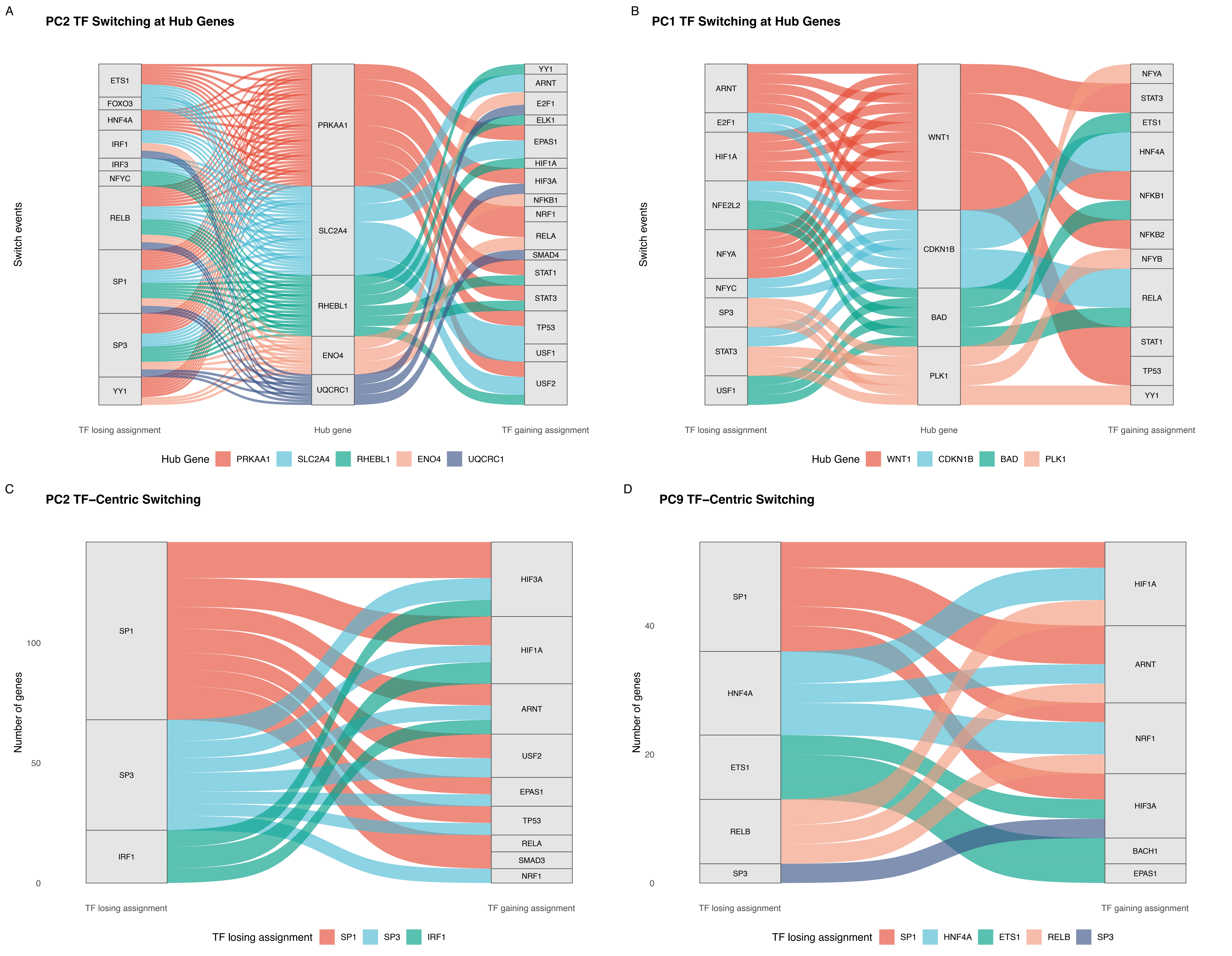

### Fig. ED2

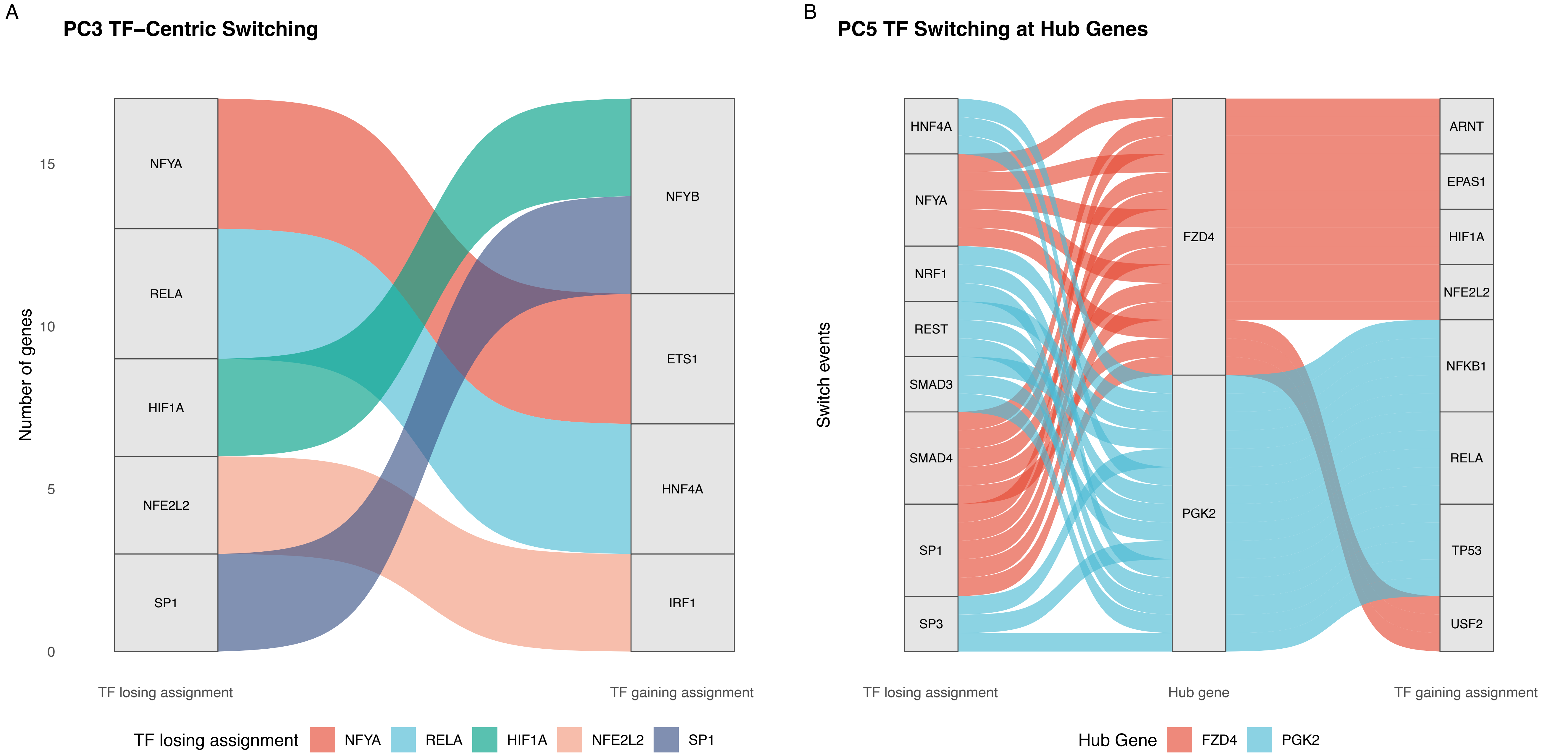

### Fig. ED3

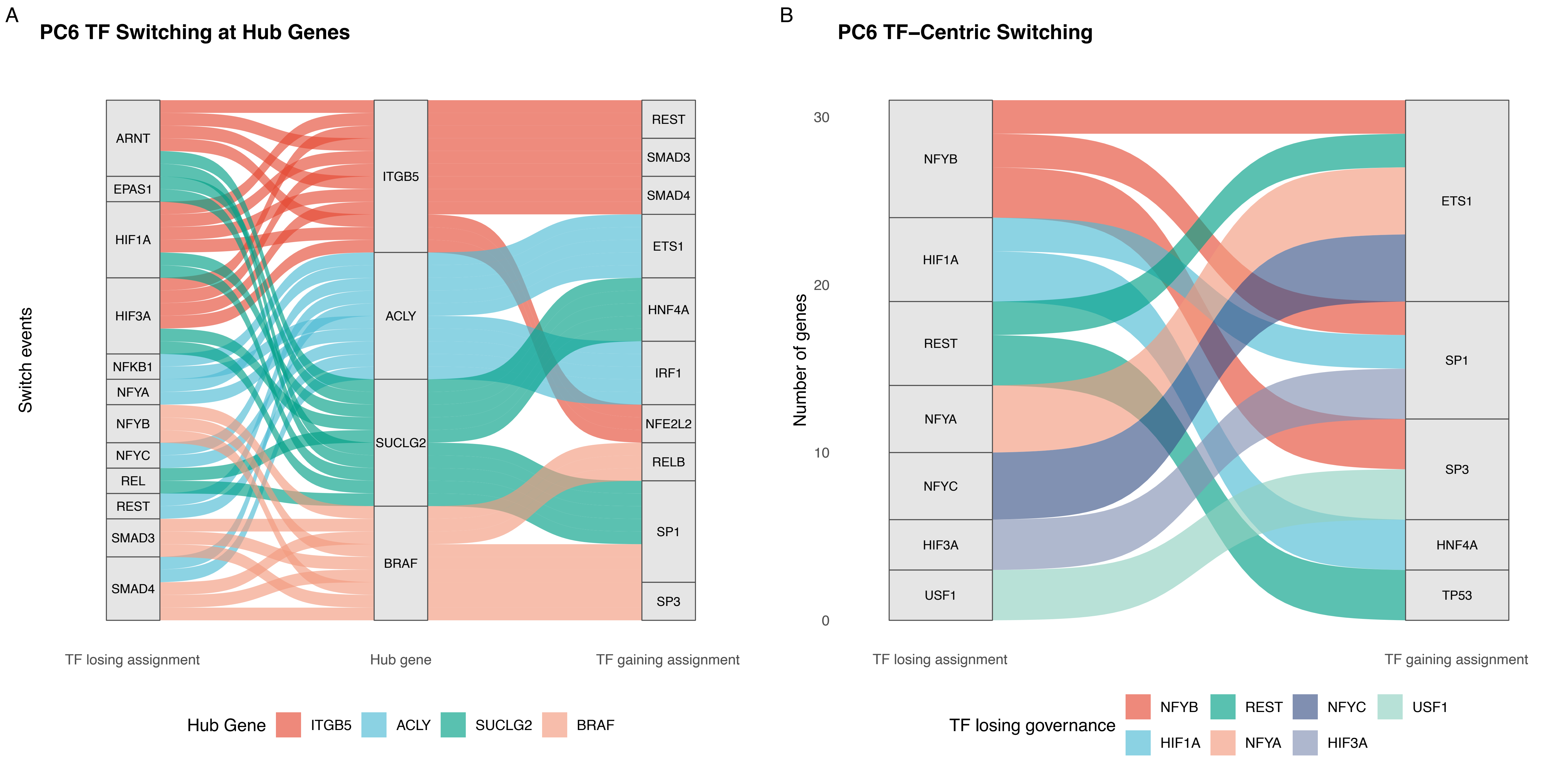

### Fig. ED4

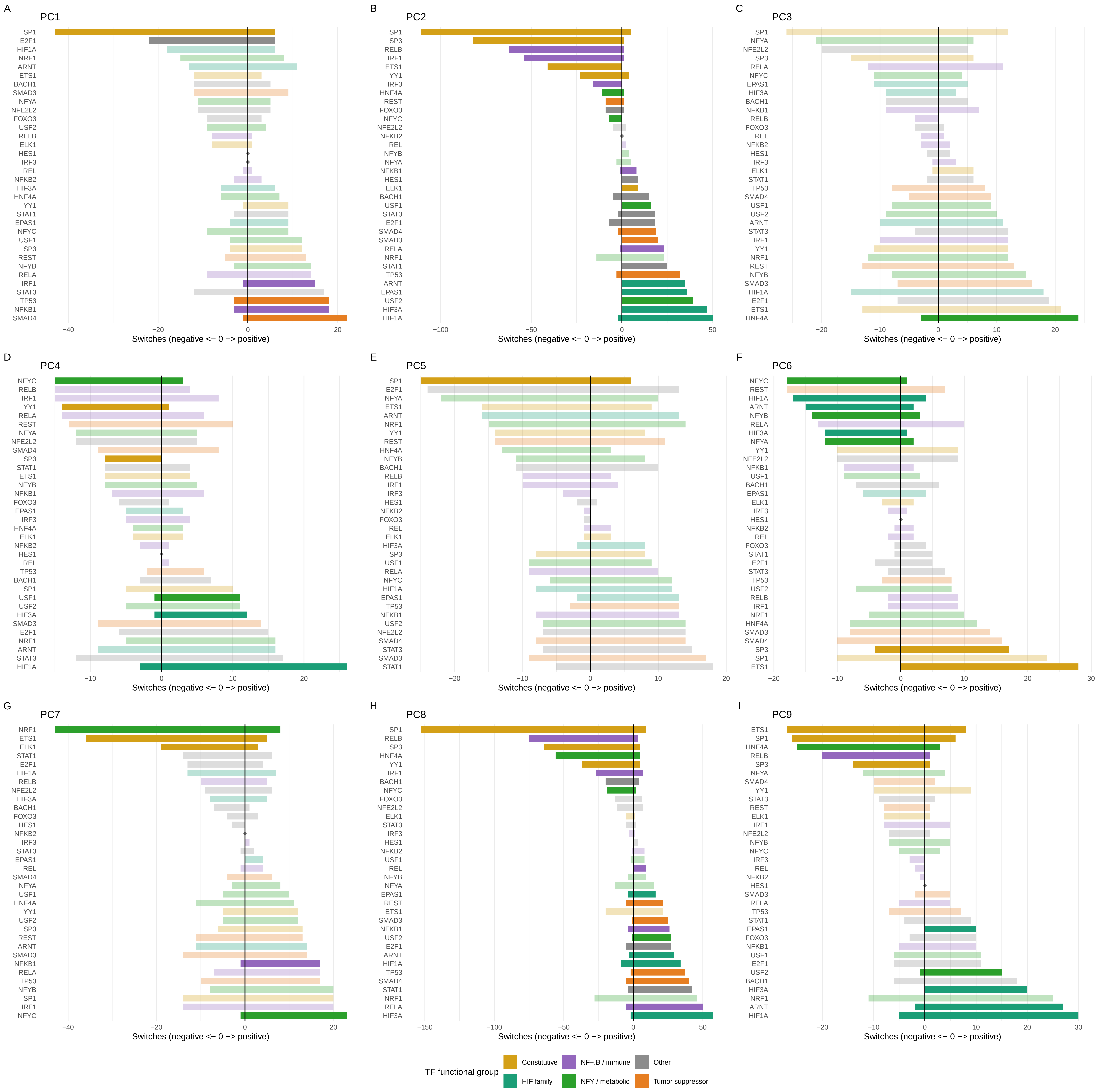
