## Supplementary material for "Oxygen-sensing regulatory architecture structures mammalian diversification": Table ED1

Table ED1 a: KEGG pathways included in the study

| **#** | **KEGG ID** | **Pathway Name** | **Tier** | **Core Biological Program** | **Primary Systems Influenced** | **Connection to O₂ Sensing** |
| --- | --- | --- | --- | --- | --- | --- |
| **1** | 04066 | HIF-1 signaling | 1 | Master hypoxia transcriptional response | Erythropoiesis, angiogenesis, metabolism, immunity | **Is the canonical oxygen-sensing pathway; PHD enzymes are direct O₂ sensors^1,2^** The HIF-1 signaling pathway is the canonical molecular oxygen-sensing system in animals. Its PHD-based sensing mechanism constitutes a “biochemical oxygen meter”, with PHD2 identified as the dominant enzyme controlling cellular HIF-1α levels. The pathway is conserved across all thirteen metazoan phyla, traceable to the last metazoan common ancestor approximately 800 million years ago, predating the Cambrian explosion. HIF-1 is a nexus where oxygen availability, metabolic state, and developmental programming converge, since the HIF-1 pathway integrates signals of hypoxia, nitric oxide, growth factors, and oncogenic signaling. |
| **2** | 00010 | Glycolysis/  Gluconeogenesis | 2 | Carbohydrate energy metabolism | All tissues; brain, muscle, liver especially | **Direct HIF-1 transcriptional output; all glycolytic genes bear HREs^3,4^** The glycolysis pathway is a direct and primary transcriptional output of HIF-1 activation. Under hypoxia, HIF-1α binds HREs in the promoters of nearly every glycolytic enzyme gene, including hexokinase 1 and 2 (HK1/2), phosphoglucose isomerase (PGI), phosphofructokinase (PFKL), aldolase A (ALDOA), glyceraldehyde-3-phosphate dehydrogenase (GAPDH), phosphoglycerate kinase 1 (PGK1), phosphoglycerate mutase 1 (PGAM1), enolase (ENO1), pyruvate kinase M (PKM), and lactate dehydrogenase A (LDHA). To increase glucose supply for enhanced glycolytic flux, HIF-1 can upregulate glucose transporters GLUT1 and GLUT3. This coordinated regulation represents the most conserved and direct metabolic output of the oxygen-sensing system. Cells under hypoxia are metabolically reprogrammed to glycolysis without requiring oxygen. The coupling between HIF-1 and glycolytic gene expression is so tight that the "Warburg effect" in cancer (aerobic glycolysis) can in part be explained by constitutive HIF-1 activation through oncogenic PI3K/Akt/mTOR signaling or VHL loss. Here, variation in how transcription factors govern glycolytic gene promoters across mammalian species reflects the evolutionary diversity of metabolic strategies under differing oxygen regimes. |
| **3** | 00020 | Citrate cycle (TCA) | 2 | Central oxidative catabolism | Heart, brain, kidney, liver | **HIF-1 suppresses via PDK1; α-KG is PHD co-substrate linking TCA to O₂ sensing^5,6^** The TCA cycle is intrinsically oxygen-dependent: its reduced products (NADH, FADH₂) are only useful if oxygen-dependent oxidative phosphorylation is operating. HIF-1 directly suppresses TCA cycle activity during hypoxia through multiple mechanisms. The induction of PDK1 by HIF-1α prevents pyruvate from entering the cycle by inactivating the pyruvate dehydrogenase complex, thereby reducing acetyl-CoA supply. Under severe or prolonged hypoxia, mitochondrial respiration becomes untenable, and cells must rely on glycolytic ATP generation, a transition mechanistically coordinated by HIF-1 suppressing TCA flux while simultaneously upregulating glycolysis. Critically, TCA cycle intermediates, particularly α-ketoglutarate, are obligate co-substrates for the PHD enzymes that hydroxylate HIF-1α. Thus, the TCA cycle directly feeds back to regulate the sensitivity of the HIF oxygen sensor itself. Hence, the TCA cycle is not merely a downstream target of HIF signaling but a direct biochemical participant in oxygen sensing. |
| **4** | 00190 | Oxidative phosphorylation | 2 | ATP generation via ETC | High-energy tissues: heart, brain, kidney | **O₂ is terminal ETC electron acceptor; HIF-1 suppresses OXPHOS in hypoxia^6^** Oxidative phosphorylation is inseparable from oxygen sensing: oxygen is the terminal electron acceptor at Complex IV, any fall in cellular oxygen directly impairs electron flow and ATP synthesis. This bioenergetic consequence of hypoxia (rising ADP/ATP and AMP/ATP ratios) triggers AMPK activation (see AMPK pathway, below), creating a direct link between OXPHOS dysfunction and energy-sensing of the oxygen-sensing network. HIF-1 actively suppresses OXPHOS gene expression during sustained hypoxia, reducing mitochondrial oxygen consumption to lower the intracellular oxygen demand and protect against mitochondrial reactive oxygen species (ROS) overproduction. This is achieved through PDK1-mediated inactivation of the TCA/OXPHOS substrate supply, downregulation of mitochondrial biogenesis, and induction of mitophagy. Conversely, EPAS1 (HIF-2α) regulates the chronic tolerance of OXPHOS activity, making the balance between HIF-1-mediated OXPHOS suppression (acute hypoxia response) and EPAS1-mediated OXPHOS maintenance (chronic hypoxia adaptation) a central axis of oxygen-sensing regulatory diversity across mammalian lineages. Here, the transcription factor governance of OXPHOS pathway genes, HIF-1-directed (suppressive) or NRF1/NFY-directed (biogenesis/maintenance), directly reflects the metabolic strategy of each lineage for managing oxygen availability. |
| **5** | 04630 | JAK-STAT signaling | 2 | Cytokine/immune signal transduction | Immune system, haematopoiesis, tissue homeostasis | **STAT3 physically interacts with HIF-1α; EPO signals via JAK2-STAT5^7,8^** STAT3 is the JAK-STAT member with the most direct and experimentally established connection to HIF signaling. Multiple studies have demonstrated that STAT3 physically interacts with HIF-1α in the nucleus under hypoxic conditions. STAT3 cooperatively activates a subset of HIF-1 target genes and is required for the full amplitude of the hypoxic transcriptional response in several cell types, including natural killer cells, where STAT3 is the essential regulator of HIF-1α accumulation, glycolysis, and oxygen consumption. Mechanistically, STAT3 enhances HIF-1α transcriptional activity by facilitating its interaction with hypoxia response elements, rather than by controlling HIF-1α stability. The JAK-STAT pathway also feeds into the oxygen-sensing network through erythropoiesis: EPO, a direct HIF-1 target gene, signals through JAK2-STAT5 in erythroid progenitors to drive red blood cell production in response to systemic hypoxia. Additionally, hypoxic suppression of STAT1 and interferon represents an immunomodulatory consequence of reduced oxygen, connecting the JAK-STAT pathway to immunosuppression of hypoxic tissue environments. |
| **6** | 04068 | FoxO signaling | 2 | Stress resistance, longevity, metabolism | Metabolism, immunity, aging, apoptosis | **FOXO3 is direct HIF-1 transcriptional target; FOXO3 also inhibits HIF-1α feedback^9,10^** FOXO3 is a direct transcriptional target of HIF-1α: under hypoxia, HIF-1α binds HREs in the FOXO3 promoter and drives its transcriptional induction, with multiple independent studies confirming that FOXO3 upregulation during hypoxia is attenuated by HIF-1α knockdown. This HIF-1-FOXO3 axis creates an important regulatory circuit: HIF-1 activates glycolysis and angiogenesis and simultaneously induces FOXO3-mediated programs of cell cycle arrest, apoptotic priming, and ROS management. Conversely, FOXO3 can inhibit HIF-1α through c-Myc-dependent mechanisms, with FOXO3 activity shown to abolish HIF-1α induction by hypoxia, establishing a negative feedback loop limiting duration and amplitude of the hypoxic transcriptional response. By interacting with the AMPK and mTOR pathways, FOXOs modulate the metabolic response to hypoxia. Were, the identification of the governance of FoxO pathway genes by HIF-family versus PI3K-Akt-associated transcription factors across mammalian lineages suggest an evolutionary balance between acute hypoxia-responsive adaptation and constitutive stress-resistance programs. |
| **7** | 04064 | NF-κB signaling | 2 | Immunity, inflammation, cell survival | Innate/adaptive immunity, all tissues | **NF-κB directly transcribes *HIF1A*; bidirectional hypoxia–inflammation coupling^11,12^** NF-κB directly regulates transcription of HIF-1α, representing one of the most functionally defined links between immune signaling and oxygen-sensing. Multiple NF-κB subunits bind an NF-κB response element in the HIF1A promoter and activate its transcription. Loss of both IKKα and IKKβ severely impairs HIF-1α stabilization under hypoxia. This shows that NF-κB-driven HIF1A transcription is required for the full hypoxic HIF response. In line with this, hypoxia activates NF-κB through mechanisms including suppression of IκB synthesis and increased mitochondrial ROS production, creating a bidirectional reinforcement loop between the two pathways. This inflammation-hypoxia axis has profound physiological consequences: at sites of infection or tissue injury (inflammation and local hypoxia are present) NF-κB-driven HIF-1 activation amplifies glycolytic metabolism in immune cells upregulating inflammatory cytokines and promotes angiogenesis for immune cell recruitment. In the evolutionary context of our study, the extent to which NF-κB family members govern oxygen-sensing pathway genes across mammalian lineages reflects how deeply immune and inflammatory programs have been integrated into the oxygen-sensing architecture, which could suggest a coupling particularly pronounced in lineages where recurrent hypoxia coincides with infectious or inflammatory challenge. |
| **8** | 04151 | PI3K-Akt signaling | 2 | Growth factor/survival signaling | Cell growth, survival, metabolism, immunity | **Akt activates mTORC1 to enhance HIF-1α translation; constitutive activation drives normoxic HIF-1^13,14^** A major upstream regulator of HIF-1α is the PI3K-Akt signaling pathway. Akt activates mTORC1 (by phosphorylating and inhibiting TSC2), which enhances HIF-1α mRNA translation through phosphorylation of the translational regulators 4E-BP1 and S6K1. Additionally, PI3K/Akt signaling activates NF-κB thereby upregulating HIF-1α transcription. In cancer cells with constitutively active PI3K/Akt (e.g., due to PTEN loss or PIK3CA mutation), HIF-1α is stabilized even under normoxic conditions, driving the Warburg effect and tumor angiogenesis independently of hypoxia. Under hypoxia, the relationship is bidirectional: hypoxia activates PI3K/Akt signaling in many cell types, and Akt-mediated HIF-1α stabilization contributes to the full hypoxic response in contexts where mRNA translation augments the stabilization achieved by PHD inhibition alone. PTEN, a main negative regulator of PI3K signaling, functions as a tumor suppressor because its loss constitutively activates HIF-1α through Akt/mTOR, coupling loss of growth control to metabolic reprogramming. Based on our study, the degree to which PI3K-Akt pathway gene promoters are governed by growth-factor-responsive versus HIF-family transcription factors across mammalian lineages reflects the evolutionary integration of growth signaling architecture with oxygen-sensing regulatory programs, particularly relevant to the resolution of Peto's paradox through analysis of tumor suppressor gene regulation. |
| **9** | 04150 | mTOR signaling | 2 | Nutrient/growth sensing and anabolism | Cell growth, metabolism, aging, immunity | **mTORC1 controls HIF-1α translation; hypoxia suppresses mTOR via REDD1/AMPK^15,16^** mTOR acts upstream of the translational machinery that determines how much HIF-1α protein accumulates under hypoxia. To amplify the hypoxic HIF response under growth-factor-stimulated or nutrient-replete conditions, mTORC1 phosphorylates 4E-BP1 and S6K1, enhancing cap-dependent translation of HIF-1α mRNA. Hypoxia inhibits mTORC1 through multiple converging mechanisms: as oxygen falls, mTORC1 is suppressed to reduce the energetically costly processes of protein synthesis and ribosome biogenesis. This suppression reduces the major consumer of cellular ATP (protein synthesis accounts for ~25–30% of basal ATP consumption) at the moment when ATP production is most compromised. The HIF-mTOR axis also regulates the metabolic switch from oxidative to glycolytic metabolism: mTORC1 normally promotes mitochondrial biogenesis and oxidative metabolism, and its hypoxic suppression therefore reinforces the HIF-1-mediated shift away from OXPHOS. Our results suggest that variation in how oxygen-sensing pathway genes are governed by mTOR-associated versus HIF-family transcription factors across mammals reflects the evolutionary architecture of the growth–oxygen coupling, a relationship that is particularly critical for understanding the large-body-size, slow-life-history axis of regulatory variation (PC1 in this study). |
| **10** | 04152 | AMPK signaling | 2 | Cellular energy homeostasis | Metabolism, autophagy, aging, ventilatory control | **Activated by hypoxia-driven ATP depletion; coordinates with HIF-1 in the cellular hypoxia response^16,17^** A hypoxia-induced fall in mitochondrial oxidative phosphorylation, depletes cellular ATP and raises the AMP:ATP ratio, which leads to activation of AMPK. This activation occurs even at moderate hypoxia levels where ATP depletion may be modest, because AMPK's sensitivity to AMP:ATP changes is amplified by the adenylate kinase reaction. On systemic level, AMPK in oxygen-sensing cells of the carotid body couples mitochondrial inhibition by hypoxia to the activation of potassium channels that trigger the ventilatory response to low oxygen. On cellular level, AMPK and HIF-1 operate as a coordinated pair: AMPK activation inhibits mTORC1, reducing HIF-1α translation, while simultaneously providing alternative energy sources (fatty acid oxidation, autophagy-released amino acids) that support cell survival during hypoxia. AMPK activity is important for HIF-1α transcriptional activity under hypoxic conditions, and the two pathways interact: HIF-1 governs the transcriptional adaptation program, while AMPK handles the immediate enzymatic and metabolic responses. Our results suggest that the governance of AMPK pathway genes by HIF-family or FOXO/NF-κB transcription factors could reflect how different mammalian lineages have wired their energy-sensing and oxygen-sensing networks together, whether treating AMPK as an acute emergency response (HIF-governed) or a constitutively integrated metabolic regulator (SP/NFY-governed). |

Table ED1 b: TFs included in the study

| **TF** | **Tier** | **Functional Group** | **Primary Biological Program** | **Oxygen-Sensing Connection** |
| --- | --- | --- | --- | --- |
| **HIF1A** | 1 | HIF family | **Acute hypoxia response, glycolysis, angiogenesis** The adaptive response to acute and severe reductions in oxygen is mediated by HIF-1α. Hundreds (200+) of oxygen-related genes involved in metabolism, angiogenesis, erythropoiesis and cell survival are directly or indirectly up-regulated by HIF-1α, a regulatory cascade that comprises cell survival adaptive program defined as the Warburg effect and which allows adaptation to low oxygen by switching on key genes required for angiogenesis, including VEGF, or modifying the general metabolism to ensure energy homeostasis. Thus, cells have developed a universal regulatory pathway under the control of a single transcription factor that serves as the hub to respond to acute hypoxia or severe reduction in ambient oxygen. | **Master oxygen sensor; PHD/VHL-regulated ^1,18^** HIF-1α is tagged for rapid degradation in an oxygen-dependent manner by prolyl hydroxylases (PHD) using oxygen-derived hydroxyl groups to hydroxylate proline residues. The modified HIF-1α is then recognized by the E3 ligase VHL (Von-Hippel-Lindau) and ubiquitinated and degraded. In low oxygen (hypoxia), PHD activity is reduced, and HIF-1α is stabilized. The stabilized alpha subunit of HIF-1 then dimerizes with the beta subunit of HIF-1 (aryl hydrocarbon receptor nuclear translocator, HIF-1β) (ARNT) and induces the transcription of target genes through the binding to hypoxia-response elements (HREs). Oxygen-dependent regulation of HIF-1α is crucial for normal cellular metabolism. |
| **EPAS1** | 1 | HIF family | **Chronic hypoxia, erythropoiesis, iron metabolism** In physiological conditions, EPAS1 (HIF-2α) is the chronic oxygen sensor in the HIF family of transcription factors. It regulates the expression of genes that control the physiological responses of erythropoiesis (development of red blood cells) including EPO, lipid and iron metabolism and vascular integrity to ensure cellular and organism growth and survival. The expression of these target genes is important for survival of animals exposed to hypoxia, particularly for EPO gene expression in the liver which is a key regulatory step of the erythropoietic response to hypoxia. Thus, whereas HIF-1α regulates a rapid and glycolytic-based hypoxic response through primary activation of gene expression, the survival of animals exposed to low oxygen levels for physiological adaptations like erythropoiesis depends on the EPAS1-dependent transcription program. Furthermore, in addition to HIF1α-dependent activation, EPAS1 also cooperates with ETS-family proteins in order to control angiogenic-specific endothelial cell gene expression. | **Sustained adaptive response; EPO regulation ^19-21^** EPAS1, like HIF-1α, also undergoes oxygen-dependent hydroxylation and degradation by VHL. However, it contains different target gene sequences and is expressed at different time points compared to HIF-1α. Furthermore, EPAS1 functions as the predominant form of HIF under prolonged periods of hypoxia, when the level of HIF-1α diminishes. Evolutionary genetics studies of high-altitude populations, such as the Tibetan population, found variations in EPAS1 associated with high-altitude adaptation in humans. |
| **ARNT** | 1 | HIF family | **Obligate HIF dimerization partner** For any HIF-α subunit (HIF-1α, HIF-2α, HIF-3α) to bind DNA, dimerization with the constitutively expressed binding partner ARNT is required. ARNT also acts as the dimerization partner for the aryl hydrocarbon receptor (AhR), linking oxygen sensing to xenobiotic metabolism. Due to competition of binding partners, when ARNT is sequestered by AhR, HIF signaling is attenuated, and vice versa. | **Required for all HIF transcriptional output ^22,23^** ARNT is necessary for the function of the HIF pathway. Loss of ARNT abolishes both HIF-1 and HIF-2 transcriptional activity. To ensure that the HIF-dependent transcriptional output is based on oxygen-dependent stabilization of HIF-α and not limited by dimerization with its binding partner, ARNT is constitutively expressed. |
| **HIF3A** | 1 | HIF family | **Dominant-negative HIF regulation (IPAS)** IPAS (Inhibitory PAS domain protein) is a splice variant of HIF3A and functions as a dominant-negative regulator of HIF-1 and HIF-2. Attached to HIF-α, IPAS forms heterodimers preventing its DNA-binding, thereby establishing a negative feedback loop that fine-tunes hypoxic gene expression. HIF3A is expressed in a tissue-specific manner (cerebellum, cornea, heart, lung) where it prevents pathological HIF overactivation and regulates vascularization. | **Negative feedback; hypoxia-induced splicing ^24,25^**  Hypoxia triggers a shift of splicing of HIF3A from activating to inhibitory isoforms, creating a mechanism of hypoxia-dependent gene regulation through post-transcriptional splicing. During prolonged hypoxia, this negative feedback prevents excessive HIF-1/HIF-2 activation, which would otherwise cause pathological overexpression of VEGF and glycolytic enzymes**.** |
| **NFKB1** | 2 | NF-κB/immune | **Innate immunity, inflammation** NF-κB (nuclear factor kappa-light-chain-enhancer of activated B cells) proteins are a family of five transcription factors that form homo- and heterodimers to regulate a wide variety of biological processes such as innate and adaptive immunity, inflammation, cell survival and proliferation. The expression of pro-inflammatory cytokines (TNF-α, IL-1β and IL-6) and chemokines as well as antimicrobial peptides are controlled by the canonical NFKB1/RELA heterodimers while the non-canonical NFKB2/RELB pathway is important for lymphoid organ development and adaptive immunity. NF-κB has also emerged as a key metabolic regulator of immunometabolic functions that coordinate the metabolic changes required for the inflammatory state and the functional activity of immune cells. | **Directly regulates *HIF1A* transcription ^26-28^** NF-κB directly regulates HIF1A gene transcription. In hypoxia, it induces HIF-1α mRNA expression. In hypoxic innate immune cells, HIF-1α activates NF-κB signaling, creating a reciprocal positive-feedback loop. This bidirectional regulation indicates that NF-κB serves as an upstream inducer and a downstream effector of the HIF pathway, coupling oxygen sensing to inflammatory and immune responses. For cell signaling, NF-κB is required for maintaining reactive oxygen species (ROS) levels in hypoxic tissues. |
| **RELA** | 2 | NF-κB/immune | **Pro-inflammatory cytokines, cell survival**  See above. | **Bidirectional HIF-NF-κB feedback loop ^26,29^** |
| **REL** | 2 | NF-κB/immune | **Lymphocyte activation**  See above. | **HIF pathway crosstalk ^29^** |
| **RELB** | 2 | NF-κB/immune | **Non-canonical NF-κB, lymphoid development**  See above. | **Constitutive regulatory layer ^29^** |
| **NFKB2** | 2 | NF-κB/immune | **Adaptive immunity**  See above. | **Non-canonical NF-κB pathway ^29^** |
| **SP1** | 2 | Constitutive | **Housekeeping gene expression (~12,000 targets)** With an estimated 12,000 binding sites in the human genome, SP1 acts as housekeeper and ensures baseline transcription of genes essential for fundamental cellular processes including cell cycle progression (cyclins, MYC), growth factor signaling (IGF1R, EGFR), apoptosis, chromatin remodeling, and telomere maintenance. | **Drives constitutive *HIF1A* transcription ^30-32^**  The constitutive transcription of the HIF1A gene is SP1-dependent. SP1 also regulates glycolytic enzyme genes (β-enolase, pyruvate kinase-M) through GC-rich promoter elements that are modulated by hypoxia. On protein level and under hypoxia, SP1 can interact with HIF-1α, and this interaction sequesters SP1 from some of its constitutive target genes. Overall, SP1 regulation can be seen as a basal, oxygen-independent layer of transcriptional control. |
| **SP3** | 2 | Constitutive | **Housekeeping; activator/repressor balance** While it recognizes the same GC-rich binding sites as its homolog SP1, SP3 can function as an activator and repressor of transcription depending on post-translational modifications, particularly SUMOylation. SP3 competes with SP1 for DNA binding sites and modulates the balance between constitutive activation and repression at housekeeping gene promoters**.** | **Hypoxia-depleted; modulates glycolytic genes ^32,33^**  In hypoxic conditions, SP3 protein levels are declining while SP1 levels remain stable, shifting the SP1/SP3 balance especially at glycolytic enzyme promoters and contributing to the hypoxic induction of glycolysis. SP3 also functions as a transcriptional repressor through epigenetic repression of oxygen-sensing target genes under normoxia. |
| **ETS1** | 2 | Constitutive | **Vascular development, angiogenesis** ETS1 acts as a master regulator of vascular development and angiogenesis by activating genes encoding VEGF receptors (Flk-1/KDR), VE-cadherin, PDGF, and matrix metalloproteinases, essential for blood vessel formation. Additionally, it plays a role in immune cells, where it regulates T cell development and cytokine production**.** | **Physical cooperator with HIF-2α ^34,35^**  By interacting with HIF-2α (but not HIF-1α), ETS1 regulates endothelial-specific gene expression, including the VEGF receptor Flk-1, making it a critical cooperating factor that channels the chronic oxygen-sensing signal (EPAS1/HIF-2α) into vascular development programs. |
| **ELK1** | 2 | Constitutive | **MAPK/ERK effector; immediate-early genes** As a member of the ternary complex factor subfamily of ETS transcription factors, ELK1 functions downstream of the MAPK/ERK signaling cascade. It regulates immediate-early genes involved in cell proliferation, differentiation, and migration. | **Physical cooperator with HIF-2α (not HIF-1α) ^36-38^**  Similar to ETS1, ELK1 interacts with HIF-2α protein but not HIF-1α and regulates genes, including CITED2, a negative-feedback regulator of HIF-1. Hence, ELK1 mediates the specificity of the HIF-2α transcriptional response. |
| **YY1** | 2 | Constitutive | **Multifunctional activator/repressor** The zinc-finger transcription factor YY1 can activate or repress transcription. It is involved in the regulation of proliferation, differentiation, apoptosis, and chromatin structure across all hallmarks of cancer. | **HIF-1α-inducible; stabilizes HIF-1α protein ^39-41^**  Three HIF-1 binding are located in the YY1 promoter. YY1 can be upregulated by HIF-1α. In turn, a positive feedback loop that amplifies HIF signaling consists of YY1 stabilizing HIF-1α protein through SUMOylation and phase separation. YY1 also regulates PGC-1β-mediated HIF-1α-modulated fatty acid β-oxidation, linking it to metabolic adaptation under hypoxia. |
| **NFE2L2** | 2 | Other | **Antioxidant defense (NRF2/ARE pathway)** NRF2 acts as a master regulator of the cellular antioxidant defense system: under oxidative stress NRF2 avoids KEAP1-mediated degradation, translocating to the nucleus, and activating transcription of antioxidant response element (ARE)-containing genes encoding phase II detoxification enzymes (NQO1, GSTs), and antioxidant proteins (HO-1, thioredoxin). | **Required for optimal HIF-1α activation** ^42,43^ In hypoxic kidney tubular epithelial cells, NRF2 is necessary for optimal HIF-1α activation. In brief, hypoxia increases ROS production, which activates NRF2, which modulates HIF-1α stability and signaling. In tumor microenvironment and ischemic tissues, the HIF-NRF2 axis coordinates the cellular response to the challenges of oxygen deprivation and oxidative stress |
| **BACH1** | 2 | Other | **Heme-dependent repression of antioxidants** BACH1 functions as heme-regulated transcriptional repressor that competes with NRF2 for binding to ARE/MARE enhancers. At baseline, BACH1 represses heme oxygenase-1 and other antioxidant genes. When intracellular heme levels rise, heme binds directly to BACH1, triggering its nuclear export and de-repression of target genes, allowing NRF2 to activate antioxidant transcription. BACH1 target genes include cell cycle regulators, linking oxidative stress sensing to proliferative control. | **Hypoxia-inducible repressor; NRF2 antagonist ^44-46^**  BACH1 functions as a hypoxia-inducible repressor of HO-1 in human cells, preventing inappropriate HO-1 induction during hypoxia (when carbon monoxide production from heme degradation could be harmful) and provides a heme-sensitive checkpoint that coordinates oxygen sensing with heme/iron metabolism. |
| **STAT3** | 2 | Other | **Cytokine signaling; immune modulation** STAT3 regulates cell proliferation, survival, angiogenesis, and immune suppression and acts as central mediator of JAK-STAT signaling. Usually activated by cytokines (IL-6, IL-10), growth factors, and oncogenes, STAT3 is constitutively activated in many cancers, where it promotes tumor immune evasion and metabolic reprogramming**.** | **Cooperates with HIF-1α at VEGF promoter ^47-49^**  In response to hypoxia, STAT3 interacts with HIF-1α and binds to the VEGF promoter, providing additional enhancement of HIF-1-mediated transcription. Additionally, STAT3 regulates HIF-1α stability through transcriptional and post-translational mechanisms, creating a feedforward loop between cytokine signaling and hypoxic gene expression. |
| **STAT1** | 2 | Other | **Interferon signaling; antiviral defense** STAT1 serves as the primary transcription factor for the antiviral and antiproliferative effects of type I and type II interferons. Furthermore, STAT1 activates genes encoding MHC molecules, antigen-processing components, and apoptotic mediators, functioning as a tumor suppressor. | **Downregulated in hypoxia; immunosuppression ^50,51^**  Under hypoxia, STAT1 is transcriptionally and translationally downregulated and chromatin accessibility at STAT1 binding sites decreases independently of HIF. This STAT1 suppression contributes to the immunosuppressive microenvironment of hypoxic tumors by impairing interferon signaling and cytotoxic T cell activation. |
| **REST** | 2 | Tumor suppressor | **Neuronal gene repression** REST controls hundreds of genes involved in neurotransmission, ion channels, and synaptic function and is a critical repressor that silences neuronal genes in non-neuronal tissues by binding RE1/NRSE elements and recruiting corepressor complexes (CoREST, mSin3A) along with HDACs. | **Represses ~20% of hypoxia-downregulated genes ^52,53^**  Hypoxia induces REST nuclear localization and REST is responsible for repressing approximately 20% of all hypoxia-repressed genes. REST mediates a negative-feedback mechanism that prevents sustained HIF overactivation by binding the HIF-1α promoter and mediating the downregulation of HIF-dependent gene expression during prolonged hypoxia. |
| **HNF4A** | 2 | NFY/metabolic | **Liver metabolism; gluconeogenesis** HNF4A is crucial for pancreatic β-cell function and intestinal epithelial development. Furthermore, HNF4A is essential for hepatocyte differentiation, lipid and glucose metabolism, and liver function by controlling thousands of liver-specific genes, including those for gluconeogenesis, bile acid synthesis, lipid transport, and drug metabolism. | **Cooperates with HIF for EPO expression ^54-56^**  Interacting with HIF, HNF4A regulates erythropoietin expression in the liver, modulating the tissue-specific oxygen-sensing output. The promoters of HIF-2α target genes in the liver are enriched for HNF4A binding sites which contributes to the tissue-specific hypoxic transcriptional response, particularly for iron-regulatory and metabolic genes. |
| **FOXO3** | 2 | Other | **Apoptosis, oxidative stress defense, longevity** FOXO3 is involved in the regulation of apoptosis, cell cycle arrest, autophagy, oxidative stress resistance, and longevity. FOXO3 transcriptionally activates genes for cell cycle inhibition (p27, Bim), DNA repair (GADD45) and antioxidant defense (MnSOD). | **Direct HIF-1 target; suppresses mitochondria in hypoxia ^9,57,58^**  Under hypoxia, HIF-1α binds to the promoter of FOXO3. The hypoxia-induced activation of FOXO3 leads to promotion cell survival by repressing nuclear-encoded mitochondrial genes, suppressing mitochondrial mass, oxygen consumption, and ROS production. Additionally, FOXO3 can inhibit HIF-1α-induced apoptosis by activating CITED2, a negative-feedback regulator of HIF-1, creating a survival-promoting circuit during oxygen deprivation. |
| **NFYA** | 2 | NFY/metabolic | **CCAAT-binding pioneer factor; chromatin access** The highly conserved heterotrimeric transcription factor (NFYA + NFYB + NFYC) NF-Y binds CCAAT motifs in ~30% of eukaryotic promoters. NF-Y can act as a pioneer factor opening chromatin and facilitating access for other transcription factors, which is enabled by the NFYB/NFYC dimer which has histone-fold domains. NF-Y regulates genes involved in cell cycle progression (cyclins A, B, CDK1), metabolism (glycolytic and mitochondrial genes), and cholesterol biosynthesis (SREBP-mediated). NF-Y activates glycolytic gene promoters but is neutral or repressive toward mitochondrial respiratory genes. | **Enriched in hypoxia-downregulated promoters ^59-61^**  The promoter regions of multiple genes that are transcriptionally downregulated during hypoxia are enriched for NF-Y binding sites. This suggests that NF-Y-mediated constitutive expression is displaced during the hypoxic transcriptional response. The fact that the competition between NF-Y-mediated constitutive expression and HIF-mediated inducible expression represents a fundamental regulatory axis in oxygen-sensing gene control, is reflected by common co-occurrence of CCAAT (NF-Y) motifs and HREs (HIF-1α) in hypoxia-responsive promoters. |
| **NFYB** | 2 | NFY/metabolic | **NF-Y histone-fold subunit**  See above. | **Required for NF-Y trimer formation ^59-61^** |
| **NFYC** | 2 | NFY/metabolic | **NF-Y histone-fold subunit**  See above. | **Mediates metabolic gene regulation ^59-61^** |
| **USF1** | 2 | NFY/metabolic | **E-box binding; glucose/lipid metabolism** USF1 and USF2 are ubiquitous transcription factors that bind E-box (CACGTG) motifs. Both factors are involved in the regulation of glucose and lipid metabolism, stress responses, and cell cycle control. USFs are key metabolic regulators that activate genes involved in glucose-responsive transcription in the liver (fatty acid synthase, glucokinase). | **HIF cooperator at overlapping E-box/HRE motifs ^62-64^**  Since the HIF-binding HRE core sequence (RCGTG) overlaps with E-box motifs, USF proteins can bind to the same or adjacent elements as HIF. To enable tissue-specific metabolic responses to hypoxia, USF2 functions with HIF to fine-tune transcriptional output at shared target promoters, |
| **USF2** | 2 | NFY/metabolic | **E-box binding; metabolic regulation**  See above. | **HIF cooperator; tissue-specific metabolic modulation ^64^** |
| **HES1** | 2 | Other | **Notch target; stem cell maintenance** As key effector of Notch signaling, the transcriptional repressor HES1 controls cell fate determination, neural stem cell maintenance, and somitogenesis. In multiple tissues, the oscillatory expression of HES1 regulates the timing of cell differentiation versus self-renewal. | **HIF-1α-Notch cooperation; hypoxia-inducible ^65-67^**  In hypoxic conditions, HES1 expression is induced through cooperation of HIF-1α and the Notch intracellular domain (NICD) at the HES1 promoter. This interaction leads to potentiation of HES1 transcription, linking oxygen sensing to stem cell maintenance and differentiation fate decisions. |
| **E2F1** | 2 | Other | **G1/S cell cycle transition** As central controller of the G1/S cell cycle transition, E2F1 regulates the expression of genes required for DNA replication (thymidine kinase, DHFR, DNA polymerase α). Depending on context, E2F1 can function as both an oncogene and a tumor suppressor, by inducing apoptosis through p53-dependent and p53-independent pathways. | **Directly regulates *EPAS1* (HIF-2α) transcription ^68,69^**  Cell-cycle-dependently, E2F1 regulates EPAS1 (HIF-2α) transcription, which provides a mechanistic link between cell proliferation and chronic oxygen sensing. The cell cycle-regulated deubiquitinase Cezanne/OTUD7B regulates E2F1 protein stability, which in turn controls HIF-2α expression levels. Through this process, the E2F1-HIF-2α axis modulates cell cycle progression and, reciprocally, the cell cycle state modulates the chronic oxygen-sensing response. |
| **IRF1** | 2 | NF-κB/immune | **Type I interferon; antiviral immunity** IRF1 activates the transcription of type I interferon genes and interferon-stimulated genes (ISGs, essential for antiviral defense), MHC class I expression, and natural killer cell maturation. Through activation of p21 and caspase genes, IRF1 also acts as tumor-suppressor. | **Directly suppressed by HIF-1α ^70^**  Under hypoxia, HIF-1α functions as a direct suppressor of IRF1 expression, resulting in a block of type I interferon production, linking oxygen sensing and innate immunity. In low-oxygen tissues, while NF-κB-mediated inflammatory cytokine production is maintained, the antiviral/antitumor interferon response is attenuated |
| **IRF3** | 2 | NF-κB/immune | **Type I interferon induction** IRF3 serves as a first responder in antiviral innate immunity, by inducing type I interferon production downstream of pathogen recognition receptors (TLRs, RIG-I), | **Suppressed by hypoxia; retains HIF-α in cytoplasm ^70,71^**  Under oxygen deprivation, IRF3 is suppressed through transcriptional repression by HIF-1α and reduced chromatin accessibility (HIF-independently). Cytoplasmic (resting) IRF3 can physically interact with HIF-1α and HIF-2α, retaining them in the cytoplasm and attenuating hypoxia signaling, thus, IRF3 acts at the intersection of oxygen sensing and antiviral defense. |
| **NRF1** | 2 | NFY/metabolic | **Mitochondrial biogenesis (TFAM, ETC subunits)** As a well-known master transcriptional regulator of mitochondrial biogenesis NRF1 activates expression of mitochondrial transcription factor A the key factor for mitochondrial DNA replication and transcription as well as nuclear-encoded electron transport chain subunits, heme biosynthetic enzymes, and the mitochondrial protein import machinery. Hence, NRF1 is the crucial coordinator of nuclear and mitochondrial genomes to regulate oxidative phosphorylation. | **ATM-activated by hypoxic ATP depletion ^72,73^**  Hypoxia causes ATP depletion which activates ATM kinase. This kinase activates NRF1 triggering mitochondrial biogenesis to restore respiratory capacity, linking oxygen availability to mitochondrial function. The promoter of NRF1 is regulated by NF-κB and CREB, connecting inflammatory (TLR4) signaling to mitochondrial biogenesis as a damage-mitigation response. In the context of oxygen-sensing, NRF1 can be seen as hub of mitochondrial integration that couples the oxygen-utilization machinery to oxygen-sensing transcriptional programs. |
| **SMAD3** | 2 | Tumor suppressor | **TGF-β signaling; growth inhibition** Acting as the principal signal transducer of the TGF-β pathway, SMAD3 mediates growth inhibition, fibrosis, immune regulation, and epithelial-mesenchymal transition. Activation of the TGF-β receptor leads to SMAD3 phosphorylation and complex formation with SMAD4. This complex translocates to the nucleus to regulate target genes. | **Selectively represses HIF-2α iron-regulatory genes ^74-76^**  Hypoxia selectively activates SMAD3 by increasing the SMAD3-SARA (SMAD anchor for receptor activation) interaction and promoting SMAD3 nuclear translocation. The SMAD3/SMAD4 complex (typically a heterotimer) represses HIF-2α-dependent iron-regulatory genes (DMT1) without affecting HIF-2α glycolytic or angiogenic targets. Under iron deficiency, SMAD3 is degraded, derepressing HIF-2α-mediated iron absorption. |
| **SMAD4** | 2 | Tumor suppressor | **Co-SMAD; TGF-β/BMP signaling** As it is required for all canonical TGF-β/BMP signaling, SMAD4 can be seen as a central communication hub and is known as the common-mediator SMAD (co-SMAD). Since its loss disrupts growth inhibition and genomic integrity, SMAD4 is one of the most commonly inactivated tumor suppressors in pancreatic, colorectal, and gastric cancers. | **Essential repressor of HIF-2α activity ^74-76^** SMAD4 DNA binding is required for the SMAD-mediated repression of HIF-2α activity. While hypoxia activates the HIF-1α/TGF-β1/SMAD signaling axis, SMAD4 upregulation in hypoxia provides negative feedback to regulate HIF-1α, creating a bidirectional regulatory relationship. |
| **TP53** | 2 | Tumor suppressor | **Tumor suppression; apoptosis; DNA repair** Known as the "guardian of the genome" TP53 is the most frequently mutated gene in human cancer. The encoded transcription factor p53 integrates signals from DNA damage, oncogene activation, and metabolic stress to induce cell cycle arrest, DNA repair, senescence, or apoptosis. p53 regulated hundreds of critical target genes including p21 (cell cycle arrest), BAX (apoptosis), PUMA (apoptosis), and TIGAR (metabolic regulation). | **Reciprocal regulation with HIF-1α; promotes HIF degradation ^77-79^**  p53 and HIF-1α can regulate each other. To reduce HIF signaling, p53 promotes MDM2-mediated degradation of HIF-1α. Under severe hypoxia, HIF-1α can stabilize p53 via MDM2 interaction, triggering apoptosis of cells unable to adapt. To provide an additional layer of p53-mediated HIF pathway control, p53 transcriptionally regulates microRNA-107, which suppresses HIF-1β (ARNT) expression. The reciprocal regulation of HIF-1α and p53 determines how cells respond to severe hypoxia, while healthy cells undergo apoptosis, p53-deficient tumor cells survive and adapt through HIF-driven programs. |

1	Semenza, G. L. Hypoxia-inducible factors in physiology and medicine. *Cell* **148**, 399-408 (2012). <https://doi.org/10.1016/j.cell.2012.01.021>

2	Luo, Z. *et al.* Hypoxia signaling in human health and diseases: implications and prospects for therapeutics. *Signal Transduct Target Ther* **7**, 218 (2022). <https://doi.org/10.1038/s41392-022-01080-1>

3	Grandjean, G. *et al.* Definition of a Novel Feed-Forward Mechanism for Glycolysis-HIF1α Signaling in Hypoxic Tumors Highlights Aldolase A as a Therapeutic Target. *Cancer Res* **76**, 4259-4269 (2016). <https://doi.org/10.1158/0008-5472.Can-16-0401>

4	Kierans, S. J. & Taylor, C. T. Regulation of glycolysis by the hypoxia-inducible factor (HIF): implications for cellular physiology. *J Physiol* **599**, 23-37 (2021). <https://doi.org/10.1113/jp280572>

5	Alberghina, L. The Warburg Effect Explained: Integration of Enhanced Glycolysis with Heterogeneous Mitochondria to Promote Cancer Cell Proliferation. *Int J Mol Sci* **24** (2023). <https://doi.org/10.3390/ijms242115787>

6	Schönenberger, M. J. & Kovacs, W. J. Hypoxia signaling pathways: modulators of oxygen-related organelles. *Front Cell Dev Biol* **3**, 42 (2015). <https://doi.org/10.3389/fcell.2015.00042>

7	Coulibaly, A. *et al.* STAT3 governs the HIF-1α response in IL-15 primed human NK cells. *Sci Rep* **11**, 7023 (2021). <https://doi.org/10.1038/s41598-021-84916-0>

8	Dinarello, A. *et al.* STAT3 and HIF1α cooperatively mediate the transcriptional and physiological responses to hypoxia. *Cell Death Discovery* **9**, 226 (2023). <https://doi.org/10.1038/s41420-023-01507-w>

9	Ferber, E. C. *et al.* FOXO3a regulates reactive oxygen metabolism by inhibiting mitochondrial gene expression. *Cell Death Differ* **19**, 968-979 (2012). <https://doi.org/10.1038/cdd.2011.179>

10	Fasano, C., Disciglio, V., Bertora, S., Lepore Signorile, M. & Simone, C. FOXO3a from the Nucleus to the Mitochondria: A Round Trip in Cellular Stress Response. *Cells* **8** (2019). <https://doi.org/10.3390/cells8091110>

11	van Uden, P., Kenneth, N. S. & Rocha, S. Regulation of hypoxia-inducible factor-1alpha by NF-kappaB. *Biochem J* **412**, 477-484 (2008). <https://doi.org/10.1042/bj20080476>

12	Shakir, D. *et al.* NF-κB is a central regulator of hypoxia-induced gene expression. *EMBO Rep* **27**, 416-432 (2026). <https://doi.org/10.1038/s44319-025-00651-x>

13	Sakitani, M. Crosstalk Between the PI3K/AKT/mTOR Signaling Pathway And The HIF-1α/CA9 Axis and its Significance in Leukemogenesis of Acute Myeloid Leukemia: Review article. *International Journal of Research in Medical and Clinical Science* **2**, 103-115 (2025). <https://doi.org/10.70829/ijrmcs.v02.i02.014>

14	Tian, Y. *et al.* PI3K/AKT signaling activates HIF1α to modulate the biological effects of invasive breast cancer with microcalcification. *npj Breast Cancer* **9**, 93 (2023). <https://doi.org/10.1038/s41523-023-00598-z>

15	Saxton, R. A. & Sabatini, D. M. mTOR Signaling in Growth, Metabolism, and Disease. *Cell* **168**, 960-976 (2017). <https://doi.org/10.1016/j.cell.2017.02.004>

16	Li, H. *et al.* Interactions between HIF-1α and AMPK in the regulation of cellular hypoxia adaptation in chronic kidney disease. *Am J Physiol Renal Physiol* **309**, F414-428 (2015). <https://doi.org/10.1152/ajprenal.00463.2014>

17	Barialai, L. *et al.* AMPK activation protects astrocytes from hypoxia‑induced cell death. *Int J Mol Med* **45**, 1385-1396 (2020). <https://doi.org/10.3892/ijmm.2020.4528>

18	Wang, G. L., Jiang, B. H., Rue, E. A. & Semenza, G. L. Hypoxia-inducible factor 1 is a basic-helix-loop-helix-PAS heterodimer regulated by cellular O2 tension. *Proc Natl Acad Sci U S A* **92**, 5510-5514 (1995). <https://doi.org/10.1073/pnas.92.12.5510>

19	Rankin, E. B. *et al.* Hypoxia-inducible factor-2 (HIF-2) regulates hepatic erythropoietin in vivo. *J Clin Invest* **117**, 1068-1077 (2007). <https://doi.org/10.1172/JCI30117>

20	Tian, H., McKnight, S. L. & Russell, D. W. Endothelial PAS domain protein 1 (EPAS1), a transcription factor selectively expressed in endothelial cells. *Genes Dev* **11**, 72-82 (1997). <https://doi.org/10.1101/gad.11.1.72>

21	Patel, S. A. & Simon, M. C. Biology of hypoxia-inducible factor-2alpha in development and disease. *Cell Death Differ* **15**, 628-634 (2008). <https://doi.org/10.1038/cdd.2008.17>

22	Mandl, M. & Depping, R. Hypoxia-inducible aryl hydrocarbon receptor nuclear translocator (ARNT) (HIF-1beta): is it a rare exception? *Mol Med* **20**, 215-220 (2014). <https://doi.org/10.2119/molmed.2014.00032>

23	Wood, S. M., Gleadle, J. M., Pugh, C. W., Hankinson, O. & Ratcliffe, P. J. The role of the aryl hydrocarbon receptor nuclear translocator (ARNT) in hypoxic induction of gene expression. Studies in ARNT-deficient cells. *J Biol Chem* **271**, 15117-15123 (1996). <https://doi.org/10.1074/jbc.271.25.15117>

24	Makino, Y., Kanopka, A., Wilson, W. J., Tanaka, H. & Poellinger, L. Inhibitory PAS domain protein (IPAS) is a hypoxia-inducible splicing variant of the hypoxia-inducible factor-3alpha locus. *J Biol Chem* **277**, 32405-32408 (2002). <https://doi.org/10.1074/jbc.C200328200>

25	Maynard, M. A. *et al.* Human HIF-3alpha4 is a dominant-negative regulator of HIF-1 and is down-regulated in renal cell carcinoma. *FASEB J* **19**, 1396-1406 (2005). <https://doi.org/10.1096/fj.05-3788com>

26	Shakir, D. *et al.* NF-kappaB is a central regulator of hypoxia-induced gene expression. *EMBO Rep* **27**, 416-432 (2026). <https://doi.org/10.1038/s44319-025-00651-x>

27	Rius, J. *et al.* NF-kappaB links innate immunity to the hypoxic response through transcriptional regulation of HIF-1alpha. *Nature* **453**, 807-811 (2008). <https://doi.org/10.1038/nature06905>

28	Belaiba, R. S. *et al.* Hypoxia up-regulates hypoxia-inducible factor-1alpha transcription by involving phosphatidylinositol 3-kinase and nuclear factor kappaB in pulmonary artery smooth muscle cells. *Mol Biol Cell* **18**, 4691-4697 (2007). <https://doi.org/10.1091/mbc.e07-04-0391>

29	Liu, T., Zhang, L., Joo, D. & Sun, S. C. NF-kappaB signaling in inflammation. *Signal Transduct Target Ther* **2**, 17023- (2017). <https://doi.org/10.1038/sigtrans.2017.23>

30	O'Connor, L., Gilmour, J. & Bonifer, C. The Role of the Ubiquitously Expressed Transcription Factor Sp1 in Tissue-specific Transcriptional Regulation and in Disease. *Yale J Biol Med* **89**, 513-525 (2016).

31	Jeong, J. K. & Park, S. Y. Transcriptional regulation of specific protein 1 (SP1) by hypoxia-inducible factor 1 alpha (HIF-1alpha) leads to PRNP expression and neuroprotection from toxic prion peptide. *Biochem Biophys Res Commun* **429**, 93-98 (2012). <https://doi.org/10.1016/j.bbrc.2012.10.086>

32	Suske, G. The Sp-family of transcription factors. *Gene* **238**, 291-300 (1999). <https://doi.org/10.1016/s0378-1119(99)00357-1>

33	Discher, D. J., Bishopric, N. H., Wu, X., Peterson, C. A. & Webster, K. A. Hypoxia regulates beta-enolase and pyruvate kinase-M promoters by modulating Sp1/Sp3 binding to a conserved GC element. *J Biol Chem* **273**, 26087-26093 (1998). <https://doi.org/10.1074/jbc.273.40.26087>

34	Elvert, G. *et al.* Cooperative interaction of hypoxia-inducible factor-2alpha (HIF-2alpha ) and Ets-1 in the transcriptional activation of vascular endothelial growth factor receptor-2 (Flk-1). *J Biol Chem* **278**, 7520-7530 (2003). <https://doi.org/10.1074/jbc.M211298200>

35	Dittmer, J. The biology of the Ets1 proto-oncogene. *Mol Cancer* **2**, 29 (2003). <https://doi.org/10.1186/1476-4598-2-29>

36	Aprelikova, O., Wood, M., Tackett, S., Chandramouli, G. V. & Barrett, J. C. Role of ETS transcription factors in the hypoxia-inducible factor-2 target gene selection. *Cancer Res* **66**, 5641-5647 (2006). <https://doi.org/10.1158/0008-5472.CAN-05-3345>

37	Wang, V., Davis, D. A., Veeranna, R. P., Haque, M. & Yarchoan, R. Characterization of the activation of protein tyrosine phosphatase, receptor-type, Z polypeptide 1 (PTPRZ1) by hypoxia inducible factor-2 alpha. *PLoS One* **5**, e9641 (2010). <https://doi.org/10.1371/journal.pone.0009641>

38	Ohnishi, S. *et al.* hypoxia-inducible factors activate CD133 promoter through ETS family transcription factors. *PLoS One* **8**, e66255 (2013). <https://doi.org/10.1371/journal.pone.0066255>

39	Sarvagalla, S., Kolapalli, S. P. & Vallabhapurapu, S. The Two Sides of YY1 in Cancer: A Friend and a Foe. *Front Oncol* **9**, 1230 (2019). <https://doi.org/10.3389/fonc.2019.01230>

40	Meliala, I. T. S., Hosea, R., Kasim, V. & Wu, S. The biological implications of Yin Yang 1 in the hallmarks of cancer. *Theranostics* **10**, 4183-4200 (2020). <https://doi.org/10.7150/thno.43481>

41	Antonio-Andres, G. *et al.* Transcriptional Regulation of Yin-Yang 1 Expression through the Hypoxia Inducible Factor-1 in Pediatric Acute Lymphoblastic Leukemia. *Int J Mol Sci* **23** (2022). <https://doi.org/10.3390/ijms23031728>

42	Bae, T., Hallis, S. P. & Kwak, M. K. Hypoxia, oxidative stress, and the interplay of HIFs and NRF2 signaling in cancer. *Exp Mol Med* **56**, 501-514 (2024). <https://doi.org/10.1038/s12276-024-01180-8>

43	Potteti, H. R. *et al.* Nrf2 mediates hypoxia-inducible HIF1alpha activation in kidney tubular epithelial cells. *Am J Physiol Renal Physiol* **320**, F464-F474 (2021). <https://doi.org/10.1152/ajprenal.00501.2020>

44	Kitamuro, T. *et al.* Bach1 functions as a hypoxia-inducible repressor for the heme oxygenase-1 gene in human cells. *J Biol Chem* **278**, 9125-9133 (2003). <https://doi.org/10.1074/jbc.M209939200>

45	Reichard, J. F., Motz, G. T. & Puga, A. Heme oxygenase-1 induction by NRF2 requires inactivation of the transcriptional repressor BACH1. *Nucleic Acids Res* **35**, 7074-7086 (2007). <https://doi.org/10.1093/nar/gkm638>

46	Sun, J. *et al.* Hemoprotein Bach1 regulates enhancer availability of heme oxygenase-1 gene. *EMBO J* **21**, 5216-5224 (2002). <https://doi.org/10.1093/emboj/cdf516>

47	Jung, J. E. *et al.* STAT3 is a potential modulator of HIF-1-mediated VEGF expression in human renal carcinoma cells. *FASEB J* **19**, 1296-1298 (2005). <https://doi.org/10.1096/fj.04-3099fje>

48	Pawlus, M. R., Wang, L. & Hu, C. J. STAT3 and HIF1alpha cooperatively activate HIF1 target genes in MDA-MB-231 and RCC4 cells. *Oncogene* **33**, 1670-1679 (2014). <https://doi.org/10.1038/onc.2013.115>

49	Dinarello, A. *et al.* STAT3 and HIF1alpha cooperatively mediate the transcriptional and physiological responses to hypoxia. *Cell Death Discov* **9**, 226 (2023). <https://doi.org/10.1038/s41420-023-01507-w>

50	Miar, A. *et al.* Hypoxia Induces Transcriptional and Translational Downregulation of the Type I IFN Pathway in Multiple Cancer Cell Types. *Cancer Res* **80**, 5245-5256 (2020). <https://doi.org/10.1158/0008-5472.CAN-19-2306>

51	Wang, J., Ni, Z., Duan, Z., Wang, G. & Li, F. Altered expression of hypoxia-inducible factor-1alpha (HIF-1alpha) and its regulatory genes in gastric cancer tissues. *PLoS One* **9**, e99835 (2014). <https://doi.org/10.1371/journal.pone.0099835>

52	Cavadas, M. A. *et al.* REST mediates resolution of HIF-dependent gene expression in prolonged hypoxia. *Sci Rep* **5**, 17851 (2015). <https://doi.org/10.1038/srep17851>

53	Cavadas, M. A. *et al.* REST is a hypoxia-responsive transcriptional repressor. *Sci Rep* **6**, 31355 (2016). <https://doi.org/10.1038/srep31355>

54	Eeckhoute, J., Formstecher, P. & Laine, B. Hepatocyte nuclear factor 4alpha enhances the hepatocyte nuclear factor 1alpha-mediated activation of transcription. *Nucleic Acids Res* **32**, 2586-2593 (2004). <https://doi.org/10.1093/nar/gkh581>

55	Odom, D. T. *et al.* Control of pancreas and liver gene expression by HNF transcription factors. *Science* **303**, 1378-1381 (2004). <https://doi.org/10.1126/science.1089769>

56	Hayhurst, G. P., Lee, Y. H., Lambert, G., Ward, J. M. & Gonzalez, F. J. Hepatocyte nuclear factor 4alpha (nuclear receptor 2A1) is essential for maintenance of hepatic gene expression and lipid homeostasis. *Mol Cell Biol* **21**, 1393-1403 (2001). <https://doi.org/10.1128/MCB.21.4.1393-1403.2001>

57	Jensen, K. S. *et al.* FoxO3A promotes metabolic adaptation to hypoxia by antagonizing Myc function. *EMBO J* **30**, 4554-4570 (2011). <https://doi.org/10.1038/emboj.2011.323>

58	Bakker, W. J., Harris, I. S. & Mak, T. W. FOXO3a is activated in response to hypoxic stress and inhibits HIF1-induced apoptosis via regulation of CITED2. *Mol Cell* **28**, 941-953 (2007). <https://doi.org/10.1016/j.molcel.2007.10.035>

59	Dolfini, D., Imbriano, C. & Mantovani, R. The role(s) of NF-Y in development and differentiation. *Cell Death Differ* **32**, 195-206 (2025). <https://doi.org/10.1038/s41418-024-01388-1>

60	Dolfini, D., Gatta, R. & Mantovani, R. NF-Y and the transcriptional activation of CCAAT promoters. *Crit Rev Biochem Mol Biol* **47**, 29-49 (2012). <https://doi.org/10.3109/10409238.2011.628970>

61	Benatti, P. *et al.* NF-Y activates genes of metabolic pathways altered in cancer cells. *Oncotarget* **7**, 1633-1650 (2016). <https://doi.org/10.18632/oncotarget.6453>

62	Corre, S. & Galibert, M. D. Upstream stimulating factors: highly versatile stress-responsive transcription factors. *Pigment Cell Res* **18**, 337-348 (2005). <https://doi.org/10.1111/j.1600-0749.2005.00262.x>

63	Fujimi, T. J. & Aruga, J. Upstream stimulatory factors, USF1 and USF2 are differentially expressed during Xenopus embryonic development. *Gene Expr Patterns* **8**, 376-381 (2008). <https://doi.org/10.1016/j.gep.2008.05.003>

64	Howcroft, T. K. *et al.* Upstream stimulatory factor regulates major histocompatibility complex class I gene expression: the U2DeltaE4 splice variant abrogates E-box activity. *Mol Cell Biol* **19**, 4788-4797 (1999). <https://doi.org/10.1128/MCB.19.7.4788>

65	Hiyama, A. *et al.* Hypoxia activates the notch signaling pathway in cells of the intervertebral disc: implications in degenerative disc disease. *Arthritis Rheum* **63**, 1355-1364 (2011). <https://doi.org/10.1002/art.30246>

66	Pistollato, F. *et al.* Interaction of hypoxia-inducible factor-1alpha and Notch signaling regulates medulloblastoma precursor proliferation and fate. *Stem Cells* **28**, 1918-1929 (2010). <https://doi.org/10.1002/stem.518>

67	Sahlgren, C., Gustafsson, M. V., Jin, S., Poellinger, L. & Lendahl, U. Notch signaling mediates hypoxia-induced tumor cell migration and invasion. *Proc Natl Acad Sci U S A* **105**, 6392-6397 (2008). <https://doi.org/10.1073/pnas.0802047105>

68	Moniz, S. *et al.* Cezanne regulates E2F1-dependent HIF2alpha expression. *J Cell Sci* **128**, 3082-3093 (2015). <https://doi.org/10.1242/jcs.168864>

69	Polager, S., Kalma, Y., Berkovich, E. & Ginsberg, D. E2Fs up-regulate expression of genes involved in DNA replication, DNA repair and mitosis. *Oncogene* **21**, 437-446 (2002). <https://doi.org/10.1038/sj.onc.1205102>

70	Peng, T., Du, S. Y., Son, M. & Diamond, B. HIF-1alpha is a negative regulator of interferon regulatory factors: Implications for interferon production by hypoxic monocytes. *Proc Natl Acad Sci U S A* **118** (2021). <https://doi.org/10.1073/pnas.2106017118>

71	Deng, H. *et al.* IRF3 attenuates hypoxia signaling by retaining HIF-alpha in the cytoplasm. *Cell Rep* **45**, 116815 (2026). <https://doi.org/10.1016/j.celrep.2025.116815>

72	Massaro, M., Baudo, G., Lee, H., Liu, H. & Blanco, E. Nuclear respiratory factor-1 (NRF1) induction drives mitochondrial biogenesis and attenuates amyloid beta-induced mitochondrial dysfunction and neurotoxicity. *Neurotherapeutics* **22**, e00513 (2025). <https://doi.org/10.1016/j.neurot.2024.e00513>

73	Suliman, H. B., Sweeney, T. E., Withers, C. M. & Piantadosi, C. A. Co-regulation of nuclear respiratory factor-1 by NFkappaB and CREB links LPS-induced inflammation to mitochondrial biogenesis. *J Cell Sci* **123**, 2565-2575 (2010). <https://doi.org/10.1242/jcs.064089>

74	Ma, X. *et al.* SMAD family member 3 (SMAD3) and SMAD4 repress HIF2alpha-dependent iron-regulatory genes. *J Biol Chem* **294**, 3974-3986 (2019). <https://doi.org/10.1074/jbc.RA118.005549>

75	Zhang, H. *et al.* Cellular response to hypoxia involves signaling via Smad proteins. *Blood* **101**, 2253-2260 (2003). <https://doi.org/10.1182/blood-2002-02-0629>

76	Sanchez-Elsner, T. *et al.* Synergistic cooperation between hypoxia and transforming growth factor-beta pathways on human vascular endothelial growth factor gene expression. *J Biol Chem* **276**, 38527-38535 (2001). <https://doi.org/10.1074/jbc.M104536200>

77	Sermeus, A. & Michiels, C. Reciprocal influence of the p53 and the hypoxic pathways. *Cell Death Dis* **2**, e164 (2011). <https://doi.org/10.1038/cddis.2011.48>

78	Zhou, C. H., Zhang, X. P., Liu, F. & Wang, W. Modeling the interplay between the HIF-1 and p53 pathways in hypoxia. *Sci Rep* **5**, 13834 (2015). <https://doi.org/10.1038/srep13834>

79	Ravi, R. *et al.* Regulation of tumor angiogenesis by p53-induced degradation of hypoxia-inducible factor 1alpha. *Genes Dev* **14**, 34-44 (2000).
